# New *Hydra* genomes reveal conserved principles of hydrozoan transcriptional regulation

**DOI:** 10.1101/2022.06.21.496857

**Authors:** Jack F. Cazet, Stefan Siebert, Hannah Morris Little, Philip Bertemes, Abby S. Primack, Peter Ladurner, Matthias Achrainer, Mark T. Fredriksen, R. Travis Moreland, Sumeeta Singh, Suiyuan Zhang, Tyra G. Wolfsberg, Christine E. Schnitzler, Andreas D. Baxevanis, Oleg Simakov, Bert Hobmayer, Celina E. Juliano

## Abstract

The epithelial and interstitial stem cells of the freshwater polyp *Hydra* are the best characterized stem cell systems in any cnidarian, providing valuable insight into cell type evolution and the origin of stemness in animals. However, little is known about the transcriptional regulatory mechanisms that determine how these stem cells are maintained and how they give rise to their diverse differentiated progeny. To address such questions, a thorough understanding of transcriptional regulation in *Hydra* is needed. To this end, we generated extensive new resources for characterizing transcriptional regulation in *Hydra*, including new genome assemblies for *Hydra oligactis* and the AEP strain of *Hydra vulgaris*, an updated whole-animal single-cell RNA-seq atlas, and genome-wide maps of chromatin interactions, chromatin accessibility, sequence conservation, and histone modifications. These data revealed the existence of large chromatin interaction domains in the *Hydra* genome that likely influence transcriptional regulation in a manner distinct from topologically associating domains in bilaterians. We also uncovered the transcriptomic profiles of two previously molecularly uncharacterized cell types, isorhiza-containing nematocytes and somatic gonad ectoderm. We identified novel candidate regulators of cell-type-specific transcription, several of which have likely been conserved at least since the divergence of *Hydra* and the jellyfish *Clytia hemisphaerica* over 200 million years ago. The resources generated in this study, which collectively represent the most comprehensive characterization of transcriptional regulation in a cnidarian to date, are accessible through a newly created genome portal, available at <u>research.nhgri.nih.gov/HydraAEP/</u>.

## Introduction

The advent of highly specialized cell-type-specific transcriptional programs played a critical role in the emergence and subsequent diversification of animal life. Decades of research have greatly advanced our understanding of the mechanisms of transcriptional regulation that underly cell identity in metazoans. However, much of that understanding is based on findings from bilaterian species. Consequently, relatively little is known about transcriptional regulation in non-bilaterian metazoans. Cnidaria is the sister phylum to Bilateria (Dunn et al. 2014), and despite having diverged over 500 million years ago, the two clades exhibit extensive homology at the molecular level. These similarities include important aspects of transcriptional regulation: both cnidarians and bilaterians use combinatorial histone modifications and distal enhancer-like cis-regulatory elements (CREs) (Schwaiger et al. 2014; Reddy et al. 2020; Murad et al. 2021), many transcription factors (TFs) in bilaterians are also present in cnidarians (Putnam et al. 2007; Technau et al. 2005; Babonis and Martindale 2017b), and the target genes of developmentally significant transcription factors are at least partially conserved across the two clades (Münder et al. 2010; Gufler et al. 2018; Hartl et al. 2019). However, beyond these general similarities, little is known about cnidarian gene regulatory networks and the mechanisms they employ to specify and maintain cellular identity. Given Cnidaria’s phylogenetic position within Metazoa, research in cnidarians is uniquely positioned to shed light on the evolutionary origins of Bilateria. In addition, many cnidarians possess remarkable abilities of self-repair and self-renewal not found in most bilaterian model systems, with species capable of whole-body regeneration (Trembley 1744; Darling et al. 2005; Bradshaw et al. 2015) and potentially biological immortality (Piraino et al. 1996; Martinez 1998; Schaible et al. 2015). Thus, a thorough characterization of transcriptional regulation in cnidarians can contribute to our understanding of both the origins and fundamental principles of transcriptional regulation of cell type in metazoans and the molecular basis for cnidarian resilience.

Species belonging to the genus *Hydra* are among the longest-studied and best-characterized cnidarian models, with the first experiments in *Hydra* dating back to 1744 (Trembley 1744). *Hydra* has since been used to study patterning (Browne 1909; Gierer and Meinhardt 1972), stem cell biology (David 2012; David and Murphy 1977; Bosch and David 1987; Bode et al. 1987), aging (Martinez 1998; Schaible et al. 2015), regeneration (Trembley 1744), and symbiosis (Fraune and Bosch 2007; Hamada et al. 2018). One of the strengths of *Hydra* as a research organism is its simplicity. In contrast to the three life cycle stages found in their close cnidarian relatives—planula, polyp, and medusa—*Hydra* species possess only a polyp stage.

This polyp is organized along a single oral-aboral axis, with a head made up of a mouth surrounded by a ring of tentacles at the oral pole and an adhesive foot at the aboral pole. Between the head and foot lies the body column, which serves as both the gut and stem cell compartment. The body is made up of two epithelial layers—endoderm and ectoderm—separated by an extracellular matrix. Interspersed throughout both epithelial layers are interstitial cells, which include digestive gland cells, neurons, germ cells, and nematocytes—the specialized stinging cells unique to cnidarians. In adult polyps, ectodermal, endodermal, and interstitial cells constitute three different cell lineages, each supported by their own stem cell population. The simplicity of this system has allowed researchers to identify every cell type in *Hydra* as well as the developmental trajectories that give rise to them (David 2012; Siebert et al. 2019), constituting a level of understanding unmatched in any other cnidarian research organism. However, the gene regulatory networks that coordinate these differentiation trajectories remain poorly understood.

Over the past 15 years, the advent of powerful tools and resources—including a reference genome (Chapman et al. 2010), a single-cell gene expression atlas (Siebert et al. 2019), knock-down techniques (Khalturin et al. 2008; Boehm et al. 2012), and transgenesis (Wittlieb et al. 2006)—has dramatically changed the nature of *Hydra* research, allowing researchers to address topics such as regeneration and patterning at the molecular level.

However, complicating the effective use of these tools is the fact that these resources were developed using different genetic backgrounds. Specifically, the currently available and recently improved (Simakov et al. 2022) reference genome was generated using strain 105 of *Hydra vulgaris* (formerly *H. magnipapillata*), whereas all transgenic *Hydra* lines and the single-cell expression atlas were generated using the AEP strain. The AEP and 105 strains belong to two distinct lineages that split ∼16 million years ago and have since undergone extensive divergence, to the extent that some have hypothesized that these two lineages constitute cryptic species (Schenkelaars et al. 2020; Schwentner and Bosch 2015; Wong et al. 2019; Martínez et al. 2010). Furthermore, the sequence divergence between these strains markedly reduces cross-strain mapping rates for sequencing data (Siebert et al. 2019), compromising researchers’ ability to leverage high-throughput sequencing approaches in AEP-derived lines. This highlights the need for a reference genome of the AEP strain that would allow researchers to take full advantage of available transgenic lines and the single-cell expression atlas.

Another appealing, though currently underutilized, strength of *Hydra* as an informative research organism is that it is relatively closely related to several other established and emerging laboratory models belonging to the clade Hydrozoa, creating opportunities for comparative studies. Recently published genomic and transcriptomic resources, including reference genomes for the green *Hydra viridissima* (Hamada et al. 2020) and the jellyfish *Clytia hemisphaerica* (Leclère et al. 2019) as well as a single-cell gene expression atlas of the *C. hemisphaerica* medusa (Chari et al. 2021), provide valuable reference points for systematic comparative analyses. The *Hydra* genus is associated with several noteworthy evolutionary gains and losses, including the loss of a medusa stage, the acquisition of stably associated endosymbionts in *H. viridissima* (Schwentner and Bosch 2015), and the loss of certain types of aboral regeneration in *H. oligactis* (Grens et al. 1996; Weimer 1928) (Fig. 1A). Thus, effectively establishing a framework for systematic comparative approaches would greatly enhance our ability to identify genes for future functional studies and interrogate both the conserved and unique aspects of *Hydra* biology.

**Figure 1.**
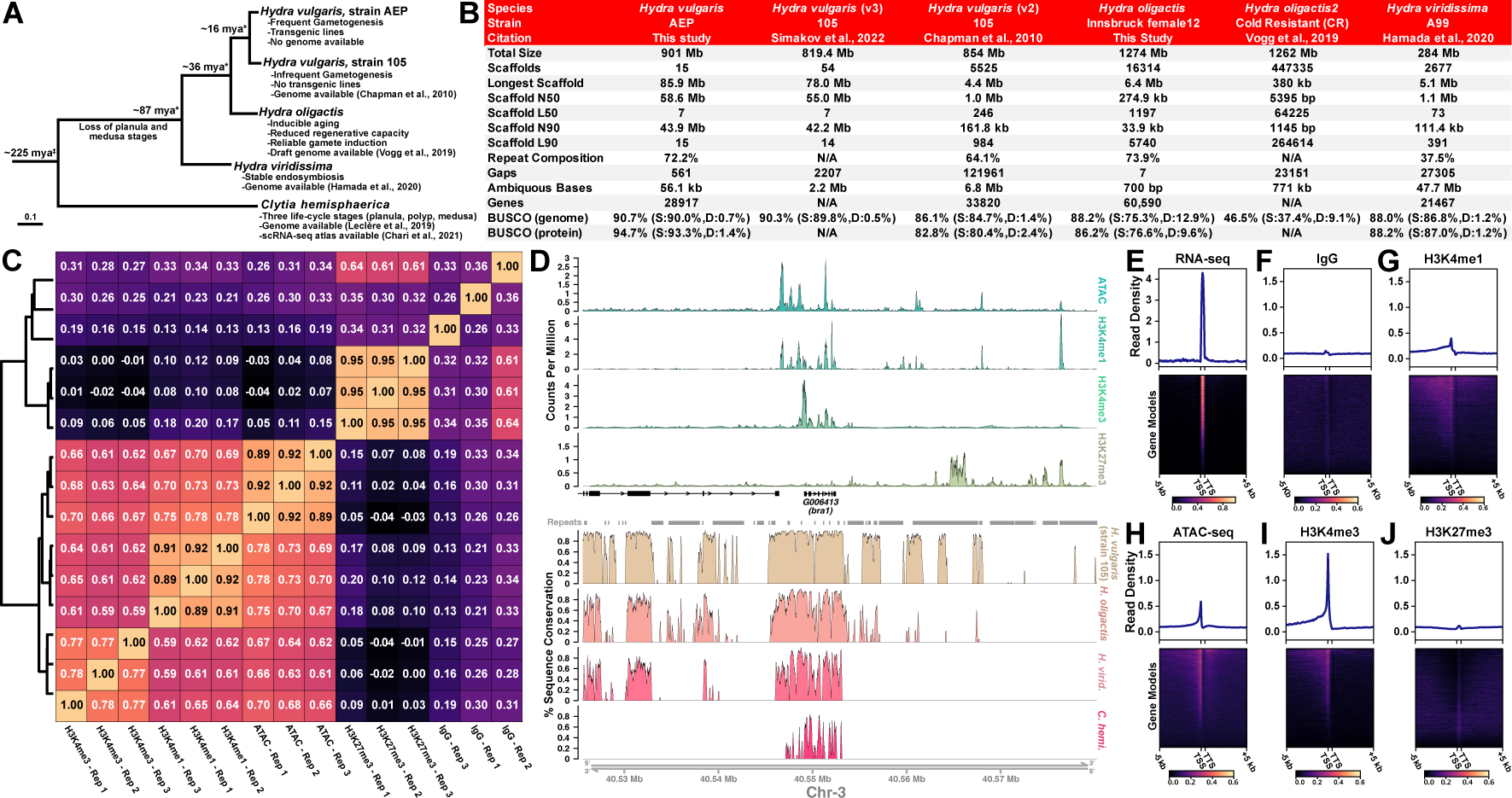
New genome assemblies provide improved resources for *Hydra* molecular biology research. *(A)* Phylogeny of hydrozoan research organisms highlighting currently available genomic and transcriptomic resources, divergence time estimates, and evolutionary gains and losses. * indicates divergence time estimates taken from Wong et al. (Wong et al. 2019). ‡ indicates divergence time estimate taken from Gold et al. (Gold et al. 2015). *(B)* The *Hydra vulgaris* strain AEP and *Hydra oligactis* genome assemblies presented in this study are marked improvements on the currently available reference genomes for their respective species. For the BUSCO statistics, S refers to the percentage of orthologs that had the expected copy number (i.e., single-copy), and D refers to the percentage of orthologs with a higher than expected copy number (i.e., the ortholog was duplicated). *(C)* Correlation analysis of genomic read distribution for *Hydra* ATAC-seq and CUT&Tag libraries shows reproducibility among biological replicates. Additionally, samples targeting active CREs (H3K4me1, H3K4me3, and ATAC-seq) were positively correlated with each other and showed no correlation to the repressive mark H3K27me3 or IgG controls. *(D)* Representative plot of CUT&Tag, ATAC-seq, and genomic conservation tracks centered on the *brachyury1* (*bra1*) gene. For the sequencing data, each track represents the signal from pooled biological replicates for the specified library type. A plot of the same locus that includes separate tracks for each CUT&Tag and ATAC-seq biological replicate is presented in Supplemental Fig. S2. (*E-J*) Read distribution for sequencing data centered on AEP assembly gene models. *(E)* Whole-animal RNA-seq data is strongly enriched in predicted coding sequences. *(F)* Control IgG CUT&Tag reads show minimal enrichment in or around genes. *(G)* H3K4me1 is enriched in promoter-proximal regions, but only weakly enriched at TSS. *(H)* ATAC-seq is enriched at TSS, but also shows some enrichment in more distal regions, likely because ATAC-seq targets both promoters and enhancers. *(I)* H3K4me3 is strongly enriched at TSS. *(J)* H3K27me3 shows minimal or no enrichment near transcribed genes.

To facilitate comparative genomic research in *Hydra*, we report two new high-quality genomes, a chromosome-level assembly for the AEP strain of *H. vulgaris* and a draft assembly for the *H. oligactis* Innsbruck female12 strain. To leverage these new references to better understand transcriptional regulation in *Hydra*, we used multiple independent approaches, such as ATAC-seq, CUT&Tag targeting histone modifications, and phylogenetic footprinting, to annotate CREs in the AEP genome. We also generated Hi-C data that revealed domains of elevated chromatin contact frequency that likely influence transcriptional regulation. To accompany these new genomic resources, we generated an updated and improved AEP-mapped version of the *Hydra* single-cell atlas, which includes two previously molecularly uncharacterized cell types. By combining our CRE annotations with the AEP single-cell atlas, we identified novel candidate regulators of cell-type-specific gene co-expression. Finally, we aligned the *Hydra* single-cell atlas with a *Clytia* medusa single-cell atlas and identified gene regulatory modules in the interstitial lineage that have been conserved over at least 200 million years of evolution. The resources generated in this study, which includes a genome browser for the *H. oligactis* and strain AEP *H. vulgaris* assemblies, a BLAST server, and an interactive portal for the AEP-mapped *Hydra* single-cell atlas, are available at <u>research.nhgri.nih.gov/HydraAEP/</u>.

## Results and Discussion

### Generation and annotation of two high-quality *Hydra* genome assemblies

We sequenced, assembled, and annotated a chromosome-level genome assembly for the AEP laboratory strain of *H. vulgaris* and a high-quality draft genome assembly for the Innsbruck female12 strain of *H. oligactis* (see Supplemental Material for details; Supplemental Fig. S1; Supplemental Data S1). The resulting assemblies were of comparable or greater completeness and contiguity when compared to other available hydrozoan genomes (Fig. 1B and Supplemental Table S1).

To augment the strain AEP *H. vulgaris* genome assembly as a resource for characterizing transcriptional regulation, we also generated genome-wide cis-regulatory element (CRE) annotations. To do this, we used the Assay for Transposase Accessible Chromatin using sequencing (ATAC-seq) (Buenrostro et al. 2013; Corces et al. 2017) to map accessible regions of chromatin. We also established a protocol for performing Cleavage Under Targets and Tagmentation (CUT&Tag) (Kaya-Okur et al. 2019) in *Hydra* to globally map multiple histone modifications, including the repressive histone modification H3K27me3 as well as the activating histone modifications H3K4me1 and H3K4me3 (Supplemental Data S2; see Supplemental Material for details). We validated the resulting data by confirming that they matched the expected distribution patterns of their associated genomic features (Fig. 1C-J and Supplemental Fig. S2).

To supplement our CRE annotations, we performed phylogenetic footprinting (Tagle et al. 1988; Gumucio et al. 1992) by using previously published genomes for the hydrozoans *Clytia hemisphaerica* (Leclère et al. 2019), *Hydra viridissima* (Hamada et al. 2020), and the 105 strain of *Hydra vulgaris* (Chapman et al. 2010)—along with our newly assembled genomes—to generate a cross-species whole-genome alignment that spans ∼225 million years of hydrozoan evolution (Fig. 1D). Importantly, our alignment yielded results that recapitulated the findings from previous manual cross-species alignments of individual *Hydra* promoter regions (Supplemental Fig. S3) (Vogg et al. 2019), supporting the accuracy of our genome-wide approach. We then used our whole-genome alignment to classify genomic features as either conserved or non-conserved based on sequence conservation across different hydrozoan genomes (see Supplemental Material for details). We provide lists of conserved non-coding genomic features in Supplemental Data S2.

### Prediction of conserved transcription factor binding sites using phylogenetic footprinting

Accurately identifying transcription factor (TF) binding sites in CREs is an essential albeit often challenging aspect of gene regulatory network characterization. This task is made especially difficult in non-bilaterian metazoans by the lack of specific antibodies needed for conventional TF mapping assays (e.g., ChIP-seq and CUT&RUN). The lack of binding data can even hinder computational approaches for predicting binding sites, as the binding preferences of cnidarian TFs typically must be inferred from data collected from distantly related bilaterians. We therefore sought to evaluate the functional relevance of bilaterian TF binding motifs in *Hydra* by leveraging phylogenetic footprinting to determine which motifs showed evidence of conservation in *Hydra*.

Of the 840 motifs considered in our analysis, 384 (45.7%) had significantly higher conservation rates than shuffled control motifs (Supplemental Data S3), including motifs for numerous conserved and developmentally significant TFs (Fig. 2A). In many cases we found that members of the same TF family, which often have similar overall binding preferences, had differing degrees of conservation. For example, we found that the TCF7L1 (JASPAR ID: MA1421.1) motif was conserved whereas the TCF7 (JASPAR ID: MA0769.2) motif was not, suggesting that the TCF7L1 motif is a better match for the binding preferences of *Hydra* TCF. By systematically differentiating between conserved and non-conserved motifs in this manner, we can improve future TF binding site predictions in *Hydra* by mitigating the risk of using functionally irrelevant motifs.

**Figure 2.**
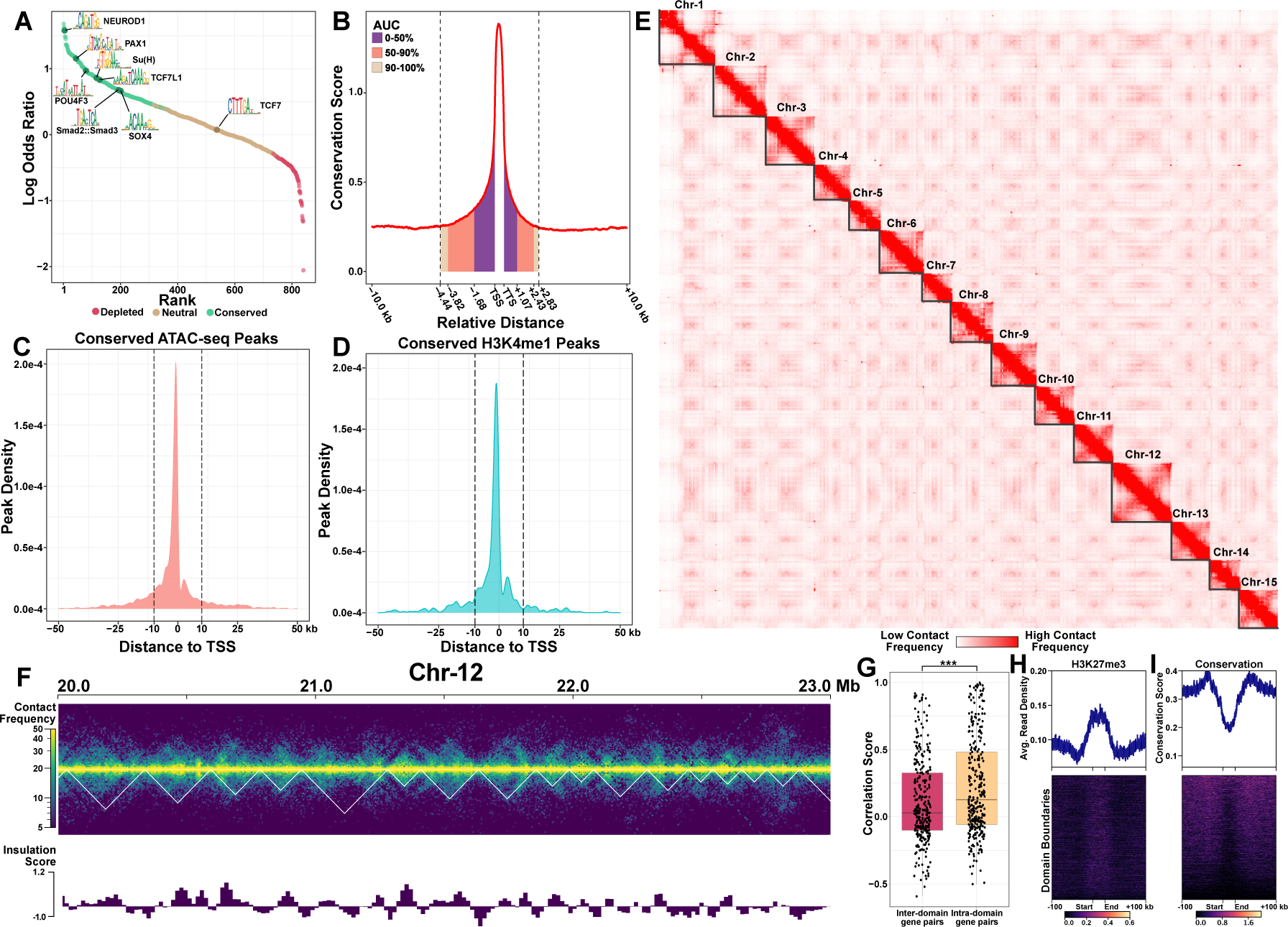
Genome sequence conservation and chromatin contact data suggest long-range chromatin interactions play a role in regulating transcription in *Hydra*. (*A*) Plot quantifying transcription factor (TF) binding motif conservation across four *Hydra* genomes. A positive log odds value indicates the non-shuffled motif had a higher conservation rate than its shuffled control. Statistical significance was evaluated using a chi-square test with an FDR cutoff of 0.01. *(B)* Plot depicting conservation patterns near genes suggests that a sizable minority of promoter-proximal CREs are farther than 2 kb from the nearest TSS. The conservation score represents the average number of non-AEP hydrozoan genomes that had the same base as the AEP assembly at a given locus. AUC, area under the curve. (*C* and *D*) Distribution of distances for conserved (*C*) ATAC-seq and (*D*) H3K4me1 CUT&Tag peaks to the nearest transcription start site (TSS) shows a strong promoter-proximal bias, although there is a non-trivial portion of peaks that are > 10 kb from the TSS. Dotted vertical lines indicate ±10 kb from the TSS. *(E)* Hi-C contact map for the *H. vulgaris* strain AEP assembly reveals 15 pseudo-chromosomes. High levels of both inter-chromosomal interactions between presumptive centromeric regions and intra-chromosomal interactions between centromeric and telomeric regions suggest a Rabl-like 3D genome conformation (Hoencamp et al. 2021). *(F)* Representative contact frequency heatmap of Hi-C data shows heterogeneity in contact frequency across distant loci. White lines delineate computationally predicted domain boundaries. The insulation score quantifies changes in local interaction frequency, with domain boundaries defined as regions with sharp changes in insulation score. *(G)* Boxplot/scatterplot depicting the correlation in expression for adjacent gene pairs show that gene pairs within the same domain (intra-domain pairs) were significantly more similar than pairs that spanned a domain boundary (inter-domain pairs; Welch two-sample t-test P-value=6.93e-05). *(H* and *I)* domain boundaries fall within regions with elevated H3K27me3 levels *(H)* and diminished sequence conservation rates *(I)*.

Another confounding issue for TF binding site prediction is that TF binding motifs are typically short and degenerate, leading to high false-positive rates when making predictions using genomic sequence alone. However, by filtering putative TF binding sites using both our ATAC-seq and phylogenetic footprinting data, we reduced the total number of predicted binding sites genome-wide by over 99%, from > 45 million to 210,122 (Supplemental Data S4). Thus, we dramatically simplified the landscape of putative TF binding sites by eliminating the multitudinous loci with a relatively low probability of being bona fide binding sites.

### Many *Hydra* genes are likely regulated by distal regulatory elements

In bilaterians, transcriptional regulation frequently involves long-range interactions between distal CREs and their target promoter, often spanning dozens of kilobases. However, numerous successful reporter lines have been generated in *Hydra* using only 500-2000 bp of flanking sequence upstream of a gene of interest, motivating some to hypothesize that transcriptional regulation in *Hydra* is simpler than in bilaterians and primarily regulated by promoter-proximal elements that typically fall within 2 kb of the TSS (Klimovich et al. 2019).

### However, this question has not been systematically investigated

To better understand the length of promoter-proximal CREs in *Hydra*, we used our cross-species whole-genome alignment to characterize sequence conservation patterns near genes in the AEP assembly (Fig. 2B). We found that flanking non-coding sequences around genes had elevated conservation rates that extended ∼4.4 kb upstream and ∼2.8 kb downstream before falling back to baseline levels. Importantly, although we found that most of the elevated sequence conservation fell within 2 kb upstream of the TSS, nearly half of the conservation signal fell outside of that boundary. This illustrates that there are many instances where conserved promoter-proximal CREs lie further than 2 kb upstream of their target gene’s TSS. In addition, although our ATAC-seq and H3K4me1 peaks showed a strong promoter-proximal bias, 5.3% (94/1782) of conserved H3K4me1 peaks and 7.8% (770/9847) of conserved ATAC-seq peaks were located further than 10 kb from the nearest gene (Fig. 2C,D), suggesting that long-range chromatin interactions play a role in regulating transcription in *Hydra*. Altogether, these observations suggest that generating a faithful reporter construct for many *Hydra* genes would require using sequence that lies further than 2 kb from the nearest TSS. The availability of the genomic resources generated by this study will enable future researchers to make more informed decisions when selecting candidate CRE sequences.

### *Hydra* chromatin is organized into localized contact domains

Our observations regarding the distribution of CREs around protein-coding genes, along with previous studies (Murad et al. 2021; Reddy et al. 2020), raised the possibility that distal enhancer-like elements play a role in regulating transcription in *Hydra*. In bilaterians, the 3D organization of the genome plays a critical role in facilitating and regulating these long-range interactions between promoters and distal CREs (Zheng and Xie 2019). However, little is known about this aspect of genome biology in cnidarians. We therefore interrogated our *Hydra* Hi-C data to better understand the 3D architecture of the *Hydra* genome.

We first examined chromatin interactions at the scale of whole chromosomes (Fig. 2E). We found that chromosomes had a Rabl-like conformation (Hoencamp et al. 2021), with interactions occurring between centromeres of different chromosomes as well as between centromeres and telomeres on individual chromosomes (Fig. 2E). Notably, these interactions appeared more prevalent in *Hydra* than in the anthozoan cnidarians *Acropora millepora* (Hoencamp et al. 2021) and *Nematostella vectensis* (Zimmermann et al. 2020), suggesting there is variability in chromosome-level nuclear architecture among cnidarians, although these differences could also be due to technical differences in Hi-C library preparation and sequencing. We noted that this apparent difference in inter-chromosome interaction correlated with the loss of multiple condensin II subunits in hydrozoans (Supplemental Fig. S4), which are required for the inhibition of a Rabl-like chromosomal conformation in bilaterians (Hoencamp et al. 2021).

Therefore, we hypothesize that the maintenance of distinct chromosome territories with few large-scale chromosome interactions is ancestral in cnidarians, and the Rabl-like conformation of *Hydra* chromatin is a derived trait caused by the loss of condensin II subunits in hydrozoans.

We next investigated local chromatin interactions in our Hi-C data to look for evidence of chromatin domains or loops, features prevalent in many bilaterian genomes that play an important role in transcriptional regulation (Zheng and Xie 2019). We observed clear heterogeneity in contact frequency between distal loci, with interaction patterns that in places resembled the triangle-shaped regions of elevated contact frequency associated with topologically associating domains (TADs) in bilaterians (Fig. 2F). Therefore, we used a previously established computational pipeline (Ramírez et al. 2018) to identify abrupt shifts in chromatin contact frequency, a feature distinctive of TAD boundaries. This analysis recovered 4,028 contact domains across the AEP assembly with a median size of ∼176 kb (Supplemental Data S5).

Notably, the expression of adjacent gene pairs that fell within the same contact domain was significantly more correlated in the *Hydra* single-cell RNA-seq atlas (described below) than adjacent gene pairs that spanned a domain boundary (Fig. 2G), suggesting that contact domains in *Hydra* are associated with transcriptional regulation.

Although *Hydra* chromatin contact domains resembled bilaterian TADs in some ways, there were important differences. Specifically, we found that domain boundaries were often located in stretches of heterochromatin with elevated H3K27me3 levels and decreased levels of sequence conservation (Fig. 2H,I). This is in stark contrast to bilaterian TAD boundaries, which are located in conserved regions of euchromatin (McArthur and Capra 2021; Akdemir et al. 2020). We also found no evidence of chromatin loops at domain boundaries, which often appear as puncta at the apex of TAD contact triangles in mammalian Hi-C data (Rao et al. 2014). Overall, our findings suggest that, although *Hydra* chromatin contact domains are associated with transcriptional regulation, domain location is likely not defined through the sequence-specific binding of regulatory factors at domain boundaries. Thus, we hypothesize that local chromatin organization in *Hydra* is achieved through mechanisms that are distinct from those that give rise to TADs in bilaterians.

In bilaterians, TADs are deeply conserved and have been linked to the preservation of local gene order (microsynteny) across the genomes of distantly related species (Harmston et al. 2017). The boundaries of these conserved TADs are associated with binding sites for CTCF, a mediator of TAD formation (Harmston et al. 2017; Heger et al. 2012). However, cnidarians and bilaterians, whose last common ancestor predates the emergence of CTCF, still exhibit extensive microsyntenic conservation (Robert et al. 2022; Heger et al. 2012). This raises the possibility that the chromatin contact domains in *Hydra* may reflect an ancestral state of chromatin organization that predates the split of Bilateria and Cnidaria.

### An updated scRNA-seq atlas for *H. vulgaris* uncovers the transcriptional profiles of additional cell types

We next focused on transcriptional regulation in the context of cell type specification in *Hydra*. This required access not only to CRE annotations but also to the transcriptomic profiles associated with different *Hydra* cell types. We previously published a whole-animal single-cell RNA-seq (scRNA-seq) dataset for the AEP strain of *H. vulgaris* that provides a molecular map of cell states in adult polyps (Siebert et al. 2019). However, the currently available versions of this dataset use either the strain 105 genome or an AEP strain transcriptome as a reference. Both are suboptimal, as the transcriptome does not provide information about genomic context—thus hindering any research into transcriptional regulation—and the 105 genome gene models are less complete and have reduced mapping rates when using AEP RNA-seq data (Supplemental Fig. S5). In addition, there have been substantial improvements in normalization (Hafemeister and Satija 2019), batch-correction (Stuart et al. 2019), and visualization techniques (McInnes et al. 2018) for scRNA-seq data since the *Hydra* single-cell atlas was initially published. Therefore, we addressed these limitations in previous iterations of the atlas by re-analyzing the scRNA-seq data using the AEP assembly as a reference.

Following mapping, we recovered 29,339 single-cell transcriptomes that passed our quality control cutoffs, an increase of ∼17.4% compared to the 24,985 transcriptomes presented in the originally published atlas (Siebert et al. 2019). We then used Seurat to perform a Louvain clustering analysis and visualized the results using a uniform manifold approximation and projection (UMAP) dimensional reduction (Hao et al. 2021; McInnes et al. 2018; Waltman and Van Eck 2013) (see Supplemental Materials for details; Fig. 3A and Supplemental Fig. S6,S7). We then annotated the resulting clusters using established cell-type markers (Siebert et al. 2019) (Supplemental Fig. S8). While generating these annotations, we identified two cell types that were not found in previous iterations of the single-cell atlas, isorhiza-containing nematocytes and ectodermal male and female somatic gonad cells, which we validated using colorimetric in-situ hybridization (Fig. 3B-H).

**Figure 3.**
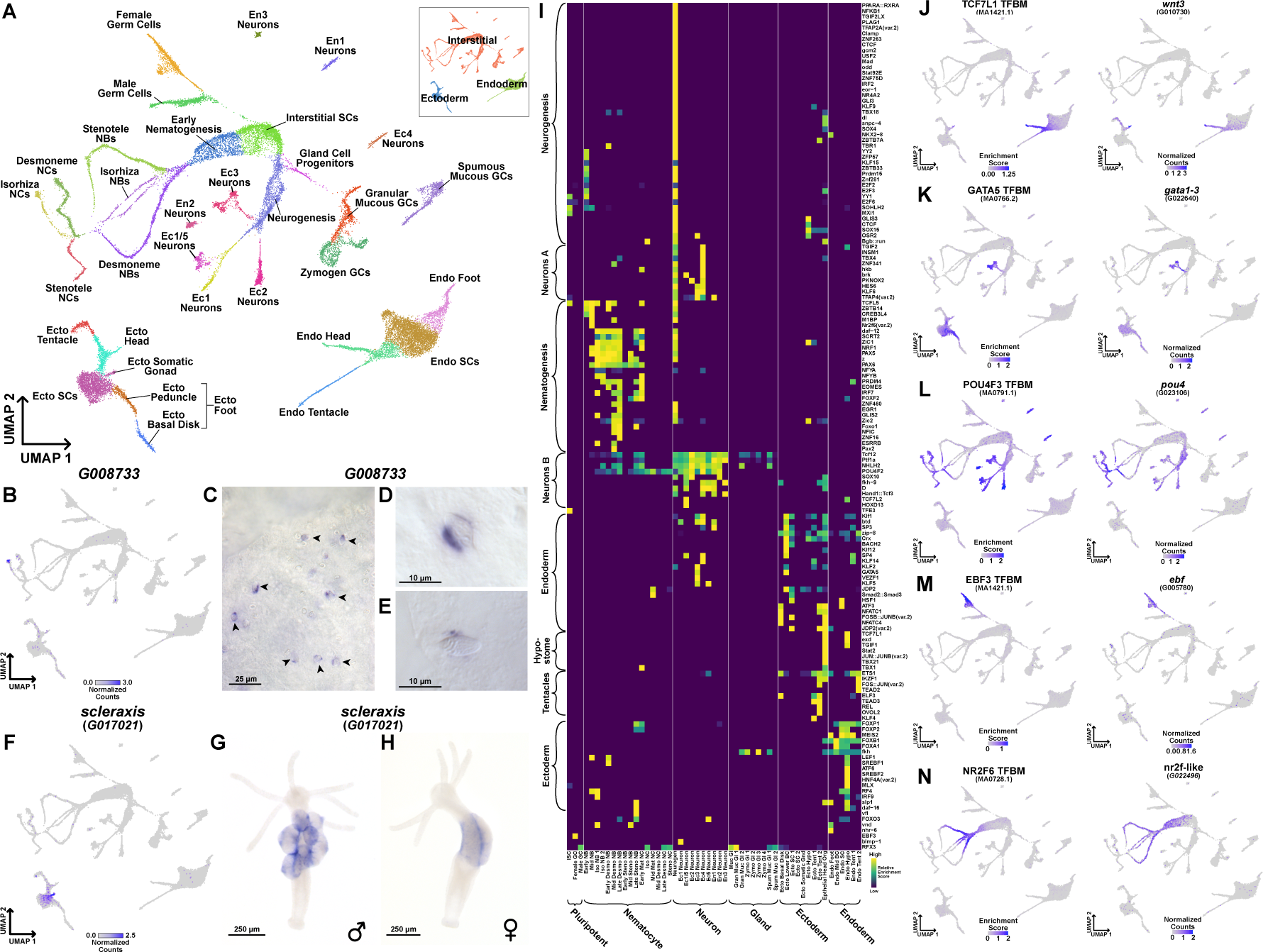
An updated *Hydra* single-cell RNA-seq atlas reveals novel regulators of gene co-expression in *Hydra*. (*A*) Uniform Manifold Approximation and Projection (UMAP) dimensional reduction of the *Hydra* single-cell RNA-seq atlas mapped to the AEP reference genome captures virtually all known cell states in adult polyps. Inset shows UMAP colored by the three stem cell lineages in adult *Hydra*. NCs, nematocytes; NBs, nematoblasts; SCs, stem cells; Ecto, ectodermal epithelial cells; Endo, endodermal epithelial cells; GCs, gland cells; Ec, neuron subtypes found in the ectoderm; En, neuron subtypes found in the endoderm. (*B*) The gene *G008733* is a specific marker for isorhiza nematocytes. (*C-E*) In-situ hybridization targeting *G008733* labels isorhiza nematocytes (black arrowheads) in upper body column tissue. (*F*) *scleraxis* is a specific marker for ectodermal somatic gonad cells. (*G*) In-situ hybridization targeting *scleraxis* in male polyps labels ectodermal testes cells. (*H*) In-situ hybridization reveals *scleraxis* is expressed in egg patches in female polyps. (*I*) Heatmap depicting TF binding motif enrichment patterns in the *Hydra* single-cell atlas. Motif redundancy was reduced by selecting a single representative from groups of motifs with similar sequence composition and enrichment patterns. A heatmap that includes all motifs is presented in Supplemental Fig. S12. (*J-N*) Motif enrichment and gene expression patterns reveal candidate regulators of cell state. (*J*) TCF motif enrichment and *wnt3* expression data corroborate the role of TCF/Wnt signaling in epithelial head tissue. (*K*) GATA motif enrichment and expression data corroborate the role of *gata1-3* in aboral epithelial tissue and suggest an additional function in Ec3 neurons. (*L*) Pou4 motif enrichment and expression data suggest *pou4* regulates transcription in differentiating and mature neurons and nematocytes. (*M*) Ebf motif enrichment and expression data suggest *ebf* regulates transcription during oogenesis. (*N*) NR2F motif enrichment and expression data suggest *nr2f-like* regulates transcription during nematogenesis. TFBM, transcription factor binding motif.

In summary, we provide an updated scRNA-seq atlas for whole adult *Hydra* that can now be used in conjunction with the AEP genome assembly. This comprehensively annotated atlas, which incorporates two additional cell types, contains virtually all known cell types in an adult *Hydra*. We also provide exhaustive lists of marker genes for all clusters (Supplemental Data S6) as well as sets of co-expressed genes (Supplemental Fig. S9; Supplemental Data S7,S8).

### Characterizing the evolutionary history of *Hydra* cell-type-specific transcriptomes

Our single-cell atlas captures the transcriptional signatures of virtually all cell states in an adult *Hydra*, which presents a unique opportunity to gain new insight into the evolutionary history of the transcriptional programs that define cnidarian cell types. The acquisition of novel cellular traits is often accompanied by a concurrent period of genetic innovation (Arendt 2008; Khalturin et al. 2009). This can leave a phylogenetic signature in a cell’s transcriptome in the form of an overrepresentation of novel genes that arose during periods of evolutionary change in a cell type’s transcriptional program (Domazet-Lošo et al. 2007). Thus, characterizing the age distribution of genes expressed in different cell types can shed light on when those genetic programs arose.

To analyze the relationship between gene age and transcriptional specificity, we first assigned phylostratigraphic ages to *Hydra* gene families using orthology predictions generated from an OrthoFinder analysis of 44 metazoan proteomes (Supplemental Fig. S10, Supplemental Table S2, and Supplemental Data S9) (Emms and Kelly 2015, 2019). We then characterized the relative enrichment of genes of a given age across different cell types in our scRNA-seq atlas, revealing clear cell-type-specific enrichment patterns (Supplemental Fig. S11A). We also calculated a holistic score, the transcriptome age index (TAI) (Domazet-Lošo and Tautz 2010), for each cell cluster (Supplemental Fig. S11B,C). Consistent with previous reports (Hemmrich et al. 2012), we found that ancient gene families predating Metazoa were most strongly associated with interstitial cells that have a high degree of potency, namely interstitial stem cells, early neuron and nematocyte progenitors, and germ cells—with interstitial stem cells having the least derived transcriptomic profile (Supplemental Fig. S11). Thus, although interstitial stem cells are likely a derived cell type unique to hydrozoans (Gold and Jacobs 2013), they make use of an evolutionarily ancient transcriptional program. This observation is consistent with the proposed existence of a deeply conserved genetic program underlying pluripotency in metazoans (Juliano et al. 2010; Sogabe et al. 2019).

Among differentiated interstitial cell types, both gland cells and neurons were enriched for genes that originated at the base of Metazoa, likely reflecting the ancient origins of their respective transcriptional programs (Supplemental Fig. S11) (Smith and Mayorova 2019; Musser et al. 2021). Neurons also showed enrichment for younger genes, suggesting the existence of cnidarian-specific modifications to neuronal transcription. In contrast, nematocyte transcriptional profiles were younger, with nematoblasts (i.e. developing nematocytes) showing stark enrichment for gene families that originated either at the base of Cnidaria or Medusozoa (Supplemental Fig. S11), consistent with the more recent evolutionary origin of nematocytes (Jung et al. 2007; David et al. 2008).

The two epithelial lineages were both associated with genes predating Cnidaria, although endodermal cell transcriptomes appeared somewhat older than those in ectodermal cells (Supplemental Fig. S11). Similar to neurons, both epithelial lineages were also enriched for younger hydrozoan-specific gene families. Little is known about the evolution of epithelial cells within hydrozoans, but the topic may merit further study as our analysis suggests these cell types may be a major driver of recent genetic novelty.

### Prediction of *Hydra* cell fate regulators

We next sought to leverage both the scRNA-seq atlas and the AEP assembly CRE annotations to identify TFs involved in coordinating *Hydra* cell-type-specific transcriptional programs. We had previously explored this question as part of the initial publication of the *Hydra* atlas using an analysis that combined ATAC-seq from strain 105 polyps with the strain AEP scRNA-seq data (Siebert et al. 2019). Broadly, our approach was first to identify TF binding motifs that were enriched in promoter-proximal CREs associated with a set of co-expressed genes, collectively referred to as a metagene. Then, we predicted candidate regulators by identifying TFs that both had similar expression to the metagene of interest and could plausibly bind one of the enriched motifs. We were motivated to revisit this analysis for two reasons: first, we could use our improved AEP-mapped version of the scRNA-seq atlas, and second, our phylogenetic footprinting data would improve our enrichment analysis by eliminating potential TF binding sites that were unlikely to be functionally relevant.

Our motif enrichment analysis identified 336 motifs that were enriched in at least one metagene in the AEP-mapped *Hydra* single-cell atlas (Fig. 3I; Supplemental Fig. S9 and S12; Supplemental Data S10), and our subsequent co-expression analysis identified 115 TFs as candidate regulators (Fig. 3J-N, Supplemental Fig. S13, and Supplemental Data S11). These candidates spanned diverse cell states and included multiple regulators whose function had been previously validated in *Hydra*, such as TCF/Wnt signaling as a regulator of oral tissue (Fig. 3J) (Lengfeld et al. 2009; Hobmayer et al. 2000; Gee et al. 2010; Broun et al. 2005), *gata1-3* as a regulator of aboral tissue (Fig. 3K) (Ferenc et al. 2021), and *zic4* as a regulator of epithelial tentacle tissue (Supplemental Fig. S13J) (Vogg et al. 2021). These results validate our analysis as a method for detecting functionally important regulatory relationships underlying cell fate decisions in *Hydra*. For *gata1-3* and *zic4,* our analysis also suggests that these TFs have uncharacterized roles in neurons, with *gata1-3* likely regulating Ec3 neurons (Fig. 3K) and *zic4* regulating neural progenitors and Ec4 neurons (Supplemental Fig. S13J). In addition to providing independent validation of previous findings, our analysis also identified uncharacterized candidate regulators. These included *pou4* as a regulator of late stage nematogenesis and neurogenesis (Fig. 3L), *ebf* as a regulator of oogenesis (Fig. 3M), and *nr2f-like* as a regulator of early nematogenesis (Fig. 3N). Our findings on these uncharacterized genes present promising opportunities for future research aimed at understanding the transcriptional regulation of cell state in *Hydra*.

### Systematic comparison of cell-type-specific transcription in *Hydra vulgaris* and *Clytia hemisphaerica*

We next extended our analysis of cell-type-specific transcription to another hydrozoan, the jellyfish *Clytia hemisphaerica*. *Hydra* and *Clytia*, while both hydrozoans, nonetheless exhibit dramatic differences at both the genomic and phenotypic level. The most recent common ancestor of *Hydra* and *Clytia* lived over 200 million years ago (Gold et al. 2015), and the protein sequence divergence between the two species is roughly equivalent to that of humans and lampreys (Supplemental Fig. S10). *Hydra* and *Clytia* also have markedly different life cycles: *Hydra* have a derived and simplified life cycle that consists only of a polyp stage, while *Clytia* have planula, colonial polyp, and medusa stages, each with distinct morphologies. Because of the extensive divergence between these lineages, identifying molecular commonalities between these two systems provides strong evidence of conservation and, by extension, functional significance.

To identify conserved cell-type-specific transcriptional patterns in *Hydra* and *Clytia*, we used reciprocal principal component analysis to align our *Hydra* scRNA-seq atlas to a recently published scRNA-seq atlas generated from whole *Clytia* medusae (Chari et al. 2021). The resulting UMAP representation accurately grouped equivalent cell types from the two species (Fig. 4A). To assess transcriptional similarities between cell types more quantitatively, we calculated an alignment score (Tarashansky et al. 2021) for all pairwise cross-species cell-type comparisons. This revealed extensive similarities between the two species, providing strong evidence of transcriptional conservation across homologous cell types (Fig. 4B). We also calculated a distance metric that quantified the overall degree of transcriptional equivalence between a given cell and similar cells in the other species (Supplemental Fig. S14)

**Figure 4.**
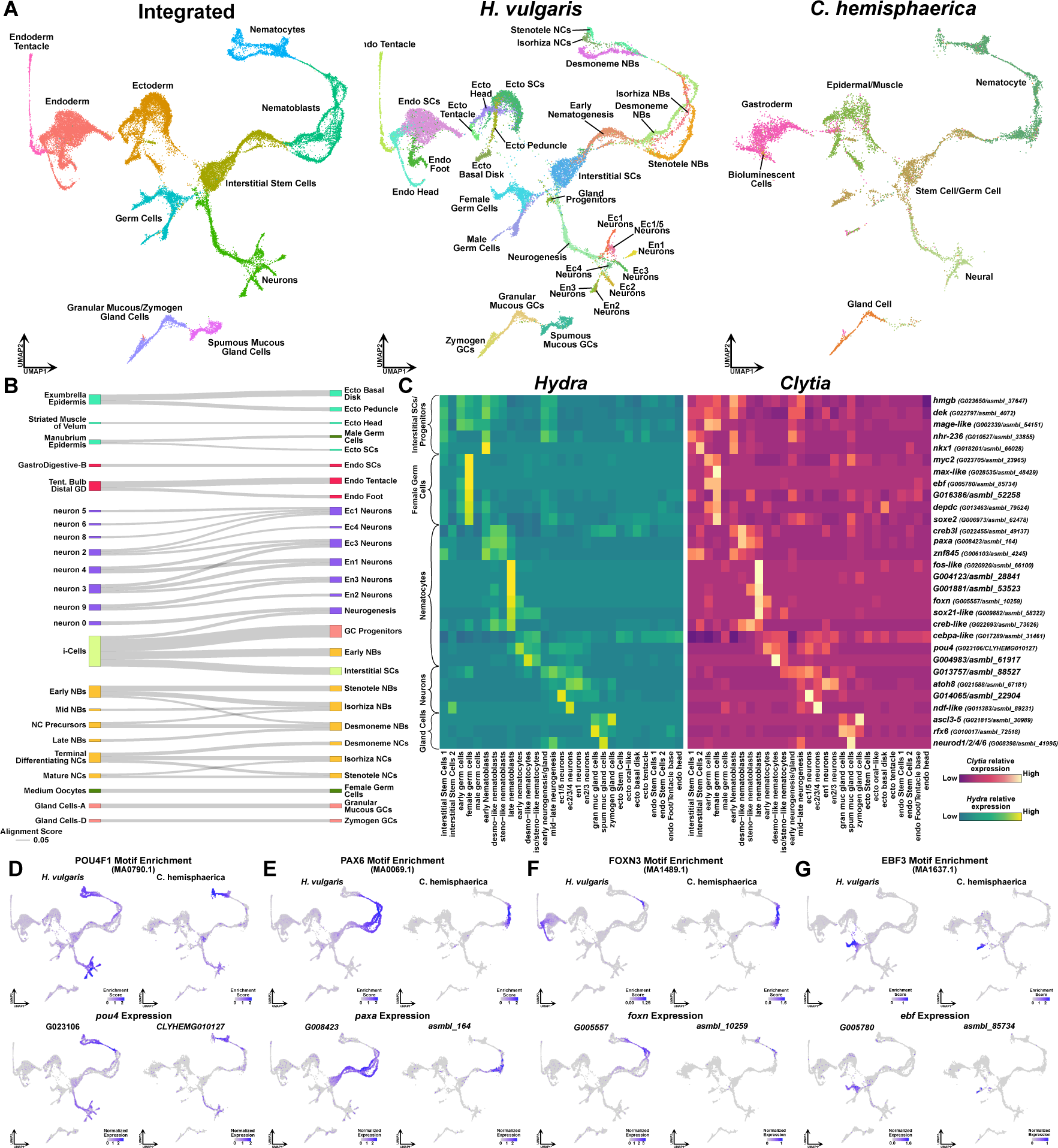
Aligned *Hydra* and *Clytia* single cell atlases reveal conserved cell-type-specific transcriptional regulation. (*A*) UMAP dimensional reduction of aligned *Hydra* and *Clytia* medusa single-cell atlases clusters together equivalent cell types from the two species. (*B*) Sankey plot showing transcriptional similarities between *Hydra* (right column) and *Clytia* (left column) cell types highlights extensive similarities amongst interstitial cell types. The alignment score quantifies the proportion of mutual nearest neighbors for one cell type that are made up of members of another cell type. An alignment score threshold of 0.05 was used to exclude poorly aligned cell types. NCs, nematocytes; NBs, nematoblasts; SCs, stem cells; Ecto, ectodermal epithelial cells; Endo, endodermal epithelial cells; GCs, gland cells; Ec, neuron subtypes found in the ectoderm; En, neuron subtypes found in the endoderm; Tent., tentacles; GD, gastroderm. (*C*) Heatmap of predicted transcription factors (TFs) with similar cell-type-specificity in *Hydra* and *Clytia*. TFs were predicted based on the presence of a predicted DNA binding domain. Orthologous gene pairs were classified as having similar expression patterns based on correlated expression (correlation score > 0.65) in the aligned cross-species principal component space. The heatmap column names refer to a fine-resolution cross-species Louvain clustering analysis presented in Supplemental Fig. S18. Heatmap values are normalized by row. (*D-G*) Conserved motif enrichment and gene expression patterns reflect gene regulatory network conservation in hydrozoans. (*D*) *pou4* is a conserved regulator of late stage and mature neurons and nematocytes. (*E*) *paxA* is a conserved regulator of nematoblasts. (*F*) *foxN* is a conserved regulator of nematocyte maturation. (*G*) *ebf* is a conserved regulator of oogenesis.

Among the three lineages, epithelial cells showed fewer cell-type similarities than interstitial cells (Fig. 4B), consistent with the dramatic differences in epithelial morphology between polyp and medusa body plans. Nonetheless, we did identify some cell-type-specific transcriptional similarities among epithelial cells, suggesting that medusa and polyp body plans in hydrozoans are created at least in part through the redeployment of shared transcriptional programs. In addition, *Hydra* epithelial stem cells had low transcriptional distance scores (Supplemental Fig. S14), potentially indicating the conservation of general epithelial transcriptional signatures despite the lack of direct homologies to individual *Clytia* epithelial cell types.

Interstitial cell types showed more robust conservation, with nearly all *Hydra* interstitial cell populations showing similarity to at least one *Clytia* cell type (Fig 4B). In some cases, there was clear one-to-one homology, such as female germline cells and some gland cell subtypes. In contrast, neuron and nematocyte cell types had either one-to-many or many-to-many patterns of homology. Thus, although neurons and nematocytes likely use a core set of conserved genes, how these transcriptional programs are deployed in a cell-type-specific manner appears to vary between *Hydra* and the *Clytia* medusa. However, because our analysis used an atlas of the *Clytia* medusa stage, as opposed to the polyp stage, it is unclear how many of the transcriptional differences between the *Hydra* and *Clytia* atlases are driven by species-specific differences versus life-cycle-specific differences. The generation of a single-cell *Clytia* polyp atlas would clarify the nature of the divergence in cell-type-specific transcription between *Clytia* and *Hydra*.

The *Hydra* genus has undergone extensive gene loss, likely as a consequence of its simplified life cycle (Chapman et al. 2010; Leclère et al. 2019; Hamada et al. 2020), but the ancestral function of these lost genes has gone largely unexplored. We sought to leverage the *Clytia* cell atlas to systematically characterize the potential function of genes lost in *Hydra*. To do this, we calculated a holistic score for each *Clytia* cell cluster that represented the degree to which that cell type expressed these lost genes (Supplemental Fig. S15). Unsurprisingly, the tentacle GFP cell—a cell type lost in *Hydra*—had one of the highest gene loss scores. Among other cell types, gland cell scores were clear outliers, with exceptionally high values across all subtypes. Notably, our cross-species cell-type comparison found that three of the five *Clytia* gland cell subtypes did not exhibit strong homology to any *Hydra* cell types (Fig. 4B). Thus, gland cell-specific genes are likely a major contributor to gene loss in the *Hydra* lineage, possibly reflecting either the loss or transcriptional simplification of gland cell subtypes.

### Interstitial cell-specific gene regulatory modules are conserved between *Hydra* and *Clytia*

Transcriptional similarities between *Hydra* and *Clytia* cell types imply the existence of conserved gene regulatory networks. Therefore, we sought to identify the regulators underlying conserved cell-type-specific expression in these two species. To that end, we re-applied the approach we used to identify candidate gene module regulators in *Hydra* to the *Clytia* single-cell atlas, albeit with some modifications because of the lack of epigenetic data in *Clytia* (Supplemental Fig. S16 and Supplemental Data S12,S13). We then compared the results from each species to identify commonalities. We found 13 motifs that had similar enrichment patterns in the two species (enrichment correlation score > 0.5) (Supplemental Data S14). Thus, despite the high level of divergence in non-coding sequence between the *Clytia* and *Hydra* genomes, we see significant overlap in the motifs associated with conserved gene co-expression modules.

To find candidate regulators of conserved gene co-expression modules, we first sought to identify TFs with similar cell-type specificity in *Hydra* and *Clytia*. To do this, we identified one-to-one ortholog pairs with correlated expression in the aligned cross-species principal component space (see Supplemental Material for details). This approach recovered 409 orthologs with highly conserved expression patterns (correlation score > 0.65), including markers for most cell types in the cross-species atlas (Supplemental Fig. S17,S18 and Supplemental Data S15). From these 409 orthologs, we identified 30 putative TFs with conserved cell-type-specific expression (Fig. 4C). While our analysis did not recover any conserved TFs in epithelial cells—likely because of the relatively poor alignability of the epithelial cell clusters— we did find conserved regulators for all interstitial cell types. We then determined which of these 30 TFs were likely to have retained their function from *Clytia* to *Hydra* by manually cross-referencing their expression patterns with our cross-species motif enrichment analysis to identify cases where both the TF expression pattern and its binding motif enrichment were conserved.

In both the *Clytia* and *Hydra* datasets, we identified *pou4* as a regulator of terminally differentiating neurons and nematocytes (Fig. 4D), *paxA* as a regulator of nematogenesis (Fig. 4E), and *atoh8*—whose orthologs are referred to as *atonal-like* (Richards and Rentzsch 2015) and *hlh6* (Chari et al. 2021) in *Nematostella* and *Clytia* respectively—as a regulator of neurogenesis (Supplemental Fig. S19). Importantly, functional data from *Nematostella*, a cnidarian that diverged from hydrozoans over 600 million years ago, are entirely consistent with our analysis’ predicted regulatory functions for these genes (Tournière et al. 2020; Babonis and Martindale 2017a; Richards and Rentzsch 2015). Our findings are also consistent with the role of *atoh8* and *pou4* in bilaterian nervous systems (Inoue et al. 2001; Gan et al. 1996). Thus, our analysis identified regulators with deeply conserved functions that either arose concurrent with or prior to the emergence of Cnidaria.

Our analysis also identified novel regulators. For example, we found that *foxN*, a TF uncharacterized in cnidarians, is likely involved in nematocyte maturation in *Hydra* and *Clytia* (Fig. 4F). We also found evidence that *ebf* is a conserved regulator of oogenesis (Fig. 4G). We noted that recently published bulk RNA-seq data from the hydrozoan *Hydractinia symbiolongicarpus* showed that *ebf* was specifically expressed in polyps undergoing oogenesis (DuBuc et al. 2020) and that an *ebf* ortholog, *ebf3b*, is a marker of female germline stem cells in zebrafish (Liu et al. 2021). Therefore, *ebf* regulation of oogenesis may predate the split of Bilateria and Cnidaria. In summary, by taking a comparative approach and leveraging the genomic and transcriptomic data available in *Clytia* and *Hydra*, we identified both conserved gene co-expression modules and the TFs that likely regulate them, providing new insight into the transcriptional programs underlying cell identity in hydrozoans.

## Conclusions

Characterizing transcriptional regulation in non-bilaterian metazoans presents significant challenges. In this study, we generated new genomes for *H. oligactis* and strain AEP *H. vulgaris*—with the latter being among the most contiguous and best-annotated cnidarian genomes currently available—to facilitate the investigation of hydrozoan transcriptional regulation. By combining our AEP strain assembly with data covering single-cell expression, chromatin accessibility, histone modifications, sequence conservation, and chromatin contact frequency, we present the most in-depth characterization of transcriptional regulation in a cnidarian to date. These data revealed that while *Hydra* CREs show a clear promoter proximal bias, there is still strong evidence of distal enhancer-like regulatory elements and large chromatin contact domains playing a role in regulating transcription. In addition, we performed an integrative analysis that predicted regulators of gene co-expression in *Hydra*, identifying both previously known and novel candidates. By extending this analysis to include a single-cell atlas of the *Clytia* medusa, we found strong evidence that the function of several candidate regulators in *Hydra* are conserved in *Clytia*, spanning over 200 million years of evolution. The resources generated by this study, available at <u>research.nhgri.nih.gov/HydraAEP/</u>, provide powerful new tools for future research aimed at unraveling hydrozoan transcriptional regulation.

## Materials and Methods

### *Hydra* Strains and Culturing Conditions

The following *Hydra vulgaris* strains were used in this study: AEP (Martin et al. 1997), 105 (Chapman et al. 2010), inverse watermelon (Glauber et al. 2015), watermelon (Glauber et al. 2015), enGreen1, and operon (Dana et al. 2012). The Innsbruck female12 strain was used for generating the *H. oligactis* genome assembly. All animals were maintained using standard procedures (Lenhoff and Brown 1970). See Supplemental Material for details.

### *Hydra vulgaris* Strain AEP Genome Sequencing, Assembly, and Annotation

High molecular weight (HMW) genomic DNA (gDNA) was extracted from strain AEP *H. vulgaris* using a Qiagen Gentra Puregene kit. The DNA was then used for the generation of 10X chromium library using a Chromium Genome Library & Gel Bead Kit v.2, an Oxford Nanopore library using an Oxford Nanopore Ligation Sequencing Kit, and a PacBio library using a SMRTbell Express Template Prep Kit 2.0. The 10X libraries were sequenced on an Illumina HiSeq X10. The Nanopore libraries were sequenced using an Oxford Nanopore PromethION. The PacBio libraries were sequenced using a PacBio Sequel II. Hi-C libraries were prepared using an Arima Hi-C kit and were sequenced on an Illumina NovaSeq 6000. The initial draft assembly was generated with the Nanopore data using Canu (Koren et al. 2017), scaffolding was performed using the 10X and Hi-C data, and error correction and gap-filling was performed using the 10X, PacBio, and Nanopore data. Gene models were generated using BRAKER2 (Brůna et al. 2021) and exonerate (Slater and Birney 2005). See Supplemental Material for details.

### *Hydra oligactis* Genome Sequencing, Assembly, and Annotation

HMW gDNA was extracted from Innsbruck female12 strain *H. oligactis* polyps using a Circulomics NanoBind BigTissue kit. Sequencing libraries were prepared using an Oxford Nanopore Ligation Sequencing Kit and sequenced using an Oxford Nanopore MinION. The assembly was generated using Flye (Kolmogorov et al. 2019). Gene models were generated using BRAKER2 (Brůna et al. 2021) See Supplemental Material for details.

### ATAC-seq

Whole-animal ATAC-seq was performed on strain AEP *H. vulgaris* polyps using a previously described protocol (Siebert et al. 2019). The data were analyzed using a previously described pipeline (Siebert et al. 2019). See Supplemental Material for details.

### CUT&Tag

Whole-animal CUT&Tag was performed on strain AEP *H. vulgaris* polyps using a modified version of the originally published protocol (Kaya-Okur et al. 2019). The data were mapped to the AEP genome assembly using Bowtie 2 (Langmead and Salzberg 2012) and peaks were called using SEACR (Meers et al. 2019). See Supplemental Material for details.

### Whole-Mount In Situ Hybridization

Whole-mount in situ hybridization was performed on strain AEP *H. vulgaris* polyps using a previously described protocol (Bode et al. 2009). See Supplemental Material for details.

### Sequence Conservation Analysis

The whole-genome cross-species alignment was generated using Progressive Cactus (Armstrong et al. 2020). Transcription factor binding sites were predicted using FIMO (Grant et al. 2011). See Supplemental Material for details.

### *Hydra* and *Clytia* Single-Cell Atlas Analysis

Previously published *Hydra* scRNA-seq data was aligned to the AEP genome assembly using a previously described pipeline (Siebert et al. 2019). The *Clytia* scRNA-seq data was mapped to gene models for the updated version of the *Clytia* genome assembly (marimba.obs-vlfr.fr) using the Cell Ranger pipeline. Normalization, clustering, cross-species alignment, and visualization was performed using Seurat (Hao et al. 2021). Gene co-expression analyses were performed using cNMF (Kotliar et al. 2019). See Supplemental Material for details.

### Data Access

We have generated a new genome portal, available at <u>research.nhgri.nih.gov/HydraAEP/</u> that allows users to interact with and download the data generated by this study. The raw sequencing data and assembled genomic sequences generated by this study has been deposited to NCBI under the BioProject ID PRJNA816482. Step-by-step descriptions of all computational analyses conducted as part of this study, including all relevant code, formatted both as markdown and HTML documents are available at <u>github.com/cejuliano/brown_hydra_genomes</u>.

## Supporting information

Cazet_Supplement

Cazet_Supplemental_DataS1

Cazet_Supplemental_DataS2

Cazet_Supplemental_DataS3

Cazet_Supplemental_DataS4

Cazet_Supplemental_DataS5

Cazet_Supplemental_DataS6

Cazet_Supplemental_DataS7

Cazet_Supplemental_DataS8

Cazet_Supplemental_DataS9

Cazet_Supplemental_DataS10

Cazet_Supplemental_DataS11

Cazet_Supplemental_DataS12

Cazet_Supplemental_DataS14

Cazet_Supplemental_DataS15

Cazet_Supplemental_DataS16

## Acknowledgments

We thank members of the C.E.J., B.H., O.S., and A.D.B. labs for valuable research discussions; Ruta Sahasrabudhe, Oanh Nguyen, Diana Burkart-Waco, and Lutz Froenicke from the DNA Technologies and Expression Analysis Core at the UC Davis Genome Center (supported by NIH Shared Instrumentation Grant 1S10OD010786-01) for technical advice and assistance with library preparation and sequencing; Bruce Draper for assistance with imaging; Anh-Dao Nguyen for assistance with generating the genome portal; Vijay Ramani for insight into metazoan chromatin organization; and Birte Mertens for assistance in establishing long-read sequencing in *H. oligactis*. This work was supported by a National Institutes of Health (NIH) Grant R35 GM133689 (to C.E.J.), an Austrian Science Fund (FWF) and Deutsche Forschungsgemeinschaft (DFG) DACH Grant I4353 (to O.S.), an Austrian Science Fund (FWF) Grant P30347 (to P.L.), and a Fonds National de la Recherche Luxembourg Grant 13569708 (to P.B.). This work was also supported in part by the Intramural Research Program of the National Human Genome Research Institute, National Institutes of Health (ZIA HG000140 to A.D.B.).

## Author Contributions

J.F.C, S.S.^a,b^, B.H., and C.E.J designed research; J.F.C, S.S. ^a,b^, H.M.L., P.B., A.S.P., M.A., P.L., and O.S. performed research; J.F.C., H.M.L., M.T.F., R.T.M., S.S.^d^, S.Z., T.G.W., C.E.S., and A.D.B contributed new analytic tools/resources; J.F.C., S.S. ^a,b^, P.B., A.S.P, and O.S. analyzed data; and J.F.C. and C.E.J wrote the paper with input from O.S., B.H., S.S. ^a,b^, and A.D.B.

## References

1. Akdemir KC, Le VT, Chandran S, Li Y, Verhaak RG, Beroukhim R, Campbell PJ, Chin L, Dixon JR, Futreal PA, et al. 2020. Disruption of chromatin folding domains by somatic genomic rearrangements in human cancer. Nat Genet 52: 294–305.

2. Arendt D. 2008. The evolution of cell types in animals: Emerging principles from molecular studies. Nat Rev Genet 9: 868–882.

3. Armstrong J, Hickey G, Diekhans M, Fiddes IT, Novak AM, Deran A, Fang Q, Xie D, Feng S, Stiller J, et al. 2020. Progressive Cactus is a multiple-genome aligner for the thousand-genome era. Nature 587: 246–251. http://dx.doi.org/10.1038/s41586-020-2871-y.

4. Babonis LS, Martindale MQ. 2017a. PaxA, but not PaxC, is required for cnidocyte development in the sea anemone Nematostella vectensis. Evodevo 8: 1–20.

5. Babonis LS, Martindale MQ. 2017b. Phylogenetic evidence for the modular evolution of metazoan signalling pathways. Philos Trans R Soc B Biol Sci 372.

6. Bode H, Lengfeld T, Hobmayer B, Holstein TW. 2009. Detection of Expression Patterns in Hydra Pattern Formation. In Wnt Signaling (ed. E. Vincan), pp. 69–84, Humana Press, Totowa, NJ https://doi.org/10.1007/978-1-60327-469-2_7.

7. Bode HR, Heimfeld S, Chow MA, Huang LW. 1987. Gland cells arise by differentiation from interstitial cells in Hydra attenuata. Dev Biol 122: 577–585.

8. Boehm AM, Khalturin K, Anton-Erxleben F, Hemmrich G, Klostermeier UC, Lopez-Quintero JA, Oberg HH, Puchert M, Rosenstiel P, Wittlieb J, et al. 2012. FoxO is a critical regulator of stem cell maintenance in immortal Hydra. Proc Natl Acad Sci U S A 109: 19697–19702. https://www.ncbi.nlm.nih.gov/pmc/articles/PMC3511741/pdf/pnas.201209714.pdf.

9. Bosch TCG, David CN. 1987. Stem cells of Hydra magnipapillata can differentiate into somatic cells and germ line cells. Dev Biol 121: 182–191. https://linkinghub.elsevier.com/retrieve/pii/0012160687901515.

10. Bradshaw B, Thompson K, Frank U. 2015. Distinct mechanisms underlie oral vs aboral regeneration in the cnidarian Hydractinia echinata. Elife 4: e05506. http://www.ncbi.nlm.nih.gov/pubmed/25884246.

11. Broun M, Gee L, Reinhardt B, Bode HR. 2005. Formation of the head organizer in hydra involves the canonical Wnt pathway. Development 132: 2907–2916. https://www.ncbi.nlm.nih.gov/pubmed/15930119.

12. Browne EN. 1909. The production of new hydranths in Hydra by the insertion of small grafts. J Exp Zool 7: 1–23. http://doi.wiley.com/10.1002/jez.1400070102.

13. Brůna T, Hoff KJ, Lomsadze A, Stanke M, Borodovsky M. 2021. BRAKER2: Automatic eukaryotic genome annotation with GeneMark-EP+ and AUGUSTUS supported by a protein database. NAR Genomics Bioinforma 3: 1–11.

14. Buenrostro JD, Giresi PG, Zaba LC, Chang HY, Greenleaf WJ. 2013. Transposition of native chromatin for fast and sensitive epigenomic profiling of open chromatin, DNA-binding proteins and nucleosome position. Nat Methods 10: 1213–1218. https://www.ncbi.nlm.nih.gov/pubmed/24097267.

15. Chapman JA, Kirkness EF, Simakov O, Hampson SE, Mitros T, Weinmaier T, Rattei T, Balasubramanian PG, Borman J, Busam D, et al. 2010. The dynamic genome of Hydra. Nature 464: 592–596. https://www.ncbi.nlm.nih.gov/pubmed/20228792.

16. Chari T, Weissbourd B, Gehring J, Ferraioli A, Leclère L, Herl M, Gao F, Chevalier S, Copley RR, Houliston E, et al. 2021. Whole-animal multiplexed single-cell RNA-seq reveals transcriptional shifts across Clytia medusa cell types. Sci Adv 7: 2021.01.22.427844. https://www.biorxiv.org/content/10.1101/2021.01.22.427844v1 https://www.biorxiv.org/content/10.1101/2021.01.22.427844v1.abstract.

17. Corces MR, Trevino AE, Hamilton EG, Greenside PG, Sinnott-Armstrong NA, Vesuna S, Satpathy AT, Rubin AJ, Montine KS, Wu B, et al. 2017. An improved ATAC-seq protocol reduces background and enables interrogation of frozen tissues. Nat Methods 14: 959–962. https://www.ncbi.nlm.nih.gov/pubmed/28846090.

18. Dana CE, Glauber KM, Chan TA, Bridge DM, Steele RE. 2012. Incorporation of a Horizontally Transferred Gene into an Operon during Cnidarian Evolution. PLoS One 7.

19. Darling JA, Reitzel AR, Burton PM, Mazza ME, Ryan JF, Sullivan JC, Finnerty JR. 2005. Rising starlet: The starlet sea anemone, Nematostella vectensis. BioEssays 27: 211–221.

20. David CN. 2012. Interstitial stem cells in Hydra: multipotency and decision-making. Int J Dev Biol 56: 489–497. http://www.intjdevbiol.com/paper.php?doi=113476cd.

21. David CN, Murphy S. 1977. Characterization of interstitial stem cells in hydra by cloning. Dev Biol 58: 372–383.

22. David CN, Özbek S, Adamczyk P, Meier S, Pauly B, Chapman J, Hwang JS, Gojobori T, Holstein TW. 2008. Evolution of complex structures: minicollagens shape the cnidarian nematocyst. Trends Genet 24: 431–438.

23. Domazet-Lošo T, Brajković J, Tautz D. 2007. A phylostratigraphy approach to uncover the genomic history of major adaptations in metazoan lineages. Trends Genet 23: 533–539. https://linkinghub.elsevier.com/retrieve/pii/S0168952507002995.

24. Domazet-Lošo T, Tautz D. 2010. A phylogenetically based transcriptome age index mirrors ontogenetic divergence patterns. Nature 468: 815–819.

25. DuBuc TQ, Schnitzler CE, Chrysostomou E, McMahon ET, Febrimarsa, Gahan JM, Buggie T, Gornik SG, Hanley S, Barreira SN, et al. 2020. Transcription factor AP2 controls cnidarian germ cell induction. Science (80-) 367: 757–762. https://www.sciencemag.org/lookup/doi/10.1126/science.aay6782.

26. Dunn CW, Giribet G, Edgecombe GD, Hejnol A. 2014. Animal phylogeny and its evolutionary implications. Annu Rev Ecol Evol Syst 45: 371–395.

27. Emms DM, Kelly S. 2019. OrthoFinder: phylogenetic orthology inference for comparative genomics. Genome Biol 20: 238. https://genomebiology.biomedcentral.com/articles/10.1186/s13059-015-0721-2.

28. Emms DM, Kelly S. 2015. OrthoFinder: solving fundamental biases in whole genome comparisons dramatically improves orthogroup inference accuracy. Genome Biol 16: 1–14. http://dx.doi.org/10.1186/s13059-015-0721-2.

29. Ferenc J, Papasaikas P, Ferralli J, Nakamura Y, Smallwood S, Tsiairis CD. 2021. Mechanical oscillations orchestrate axial patterning through Wnt activation in Hydra. Sci Adv 7: eabj6897. http://www.ncbi.nlm.nih.gov/pubmed/34890235.

30. Fraune S, Bosch TCG. 2007. Long-term maintenance of species-specific bacterial microbiota in the basal metazoan Hydra. Proc Natl Acad Sci U S A 104: 13146–13151.

31. Gan L, Xiang M, Zhou L, Wagner DS, Klein WH, Nathans J. 1996. POU domain factor Brn-3b is required for the development of a large set of retinal ganglion cells. Proc Natl Acad Sci U S A 93: 3920–3925.

32. Gee L, Hartig J, Law L, Wittlieb J, Khalturin K, Bosch TC, Bode HR. 2010. β-catenin plays a central role in setting up the head organizer in hydra. Dev Biol 340: 116–124. https://www.ncbi.nlm.nih.gov/pubmed/20045682.

33. Gierer A, Meinhardt H. 1972. A Theory of Biological Pattern Formation. Kybernetik 12: 30–39. https://link.springer.com/content/pdf/10.1007%2FBF00289234.pdf.

34. Glauber KM, Dana CE, Park SS, Colby DA, Noro Y, Fujisawa T, Chamberlin AR, Steele RE, Glauber KM, Dana CE, et al. 2015. A small molecule screen identifies a novel compound that induces a homeotic transformation in Hydra, (Development (Cambridge), (2015) 142, 4788-4796). Dev 142: 2081.

35. Gold DA, Jacobs DK. 2013. Stem cell dynamics in Cnidaria: Are there unifying principles? Dev Genes Evol 223: 53–66.

36. Gold DA, Runnegar B, Gehling JG, Jacobs DK. 2015. Ancestral state reconstruction of ontogeny supports a bilaterian affinity for Dickinsonia. Evol Dev 17: 315–324.

37. Grant CE, Bailey TL, Noble WS. 2011. FIMO: Scanning for occurrences of a given motif. Bioinformatics 27: 1017–1018.

38. Grens A, Gee L, Fisher DA, Bode HR. 1996. CnNK-2, an NK-2 homeobox gene, has a role in patterning the basal end of the axis in hydra. Dev Biol 180: 473–488. http://ac.els-cdn.com/S0012160696903218/1-s2.0-S0012160696903218-main.pdf?_tid=c2f38386-99a4-11e6-ba5000000aacb360&acdnat=1477284541_12d6755924f9c5db97b6dbcbaf1816af.

39. Gufler S, Artes B, Bielen H, Krainer I, Eder M-K, Falschlunger J, Bollmann A, Ostermann T, Valovka T, Hartl M, et al. 2018. β-Catenin acts in a position-independent regeneration response in the simple eumetazoan Hydra. Dev Biol 433: 310–323. https://www.sciencedirect.com/science/article/pii/S0012160617302907?via%3Dihub (Accessed February 7, 2019).

40. Gumucio DL, Heilstedt-Williamson H, Gray TA, Tarlé SA, Shelton DA, Tagle DA, Slightom JL, Goodman M, Collins FS. 1992. Phylogenetic footprinting reveals a nuclear protein which binds to silencer sequences in the human gamma and epsilon globin genes. Mol Cell Biol 12: 4919–4929.

41. Hafemeister C, Satija R. 2019. Normalization and variance stabilization of single-cell RNA-seq data using regularized negative binomial regression. Genome Biol 20: 296. http://www.ncbi.nlm.nih.gov/pubmed/31870423.

42. Hamada M, Satoh N, Khalturin K. 2020. A Reference Genome from the Symbiotic Hydrozoan, Hydra viridissima. G3 (Bethesda) 10: 3883–3895.

43. Hamada M, Schröder K, Bathia J, Kürn U, Fraune S, Khalturina M, Khalturin K, Shinzato C, Satoh N, Bosch TCG. 2018. Metabolic co-dependence drives the evolutionarily ancient Hydra– Chlorella symbiosis. Elife 7: 1–37.

44. Hao Y, Hao S, Andersen-Nissen E, Mauck WM, Zheng S, Butler A, Lee MJ, Wilk AJ, Darby C, Zager M, et al. 2021. Integrated analysis of multimodal single-cell data. Cell 184: 3573–3587.e29. https://doi.org/10.1016/j.cell.2021.04.048.

45. Harmston N, Ing-Simmons E, Tan G, Perry M, Merkenschlager M, Lenhard B. 2017. Topologically associating domains are ancient features that coincide with Metazoan clusters of extreme noncoding conservation. Nat Commun 8. http://dx.doi.org/10.1038/s41467-017-00524-5.

46. Hartl M, Glasauer S, Gufler S, Raffeiner A, Puglisi K, Breuker K, Bister K, Hobmayer B. 2019. Differential regulation of myc homologs by Wnt/β-Catenin signaling in the early metazoan Hydra. FEBS J 286: 2295–2310.

47. Heger P, Marin B, Bartkuhn M, Schierenberg E, Wiehe T. 2012. The chromatin insulator CTCF and the emergence of metazoan diversity. Proc Natl Acad Sci U S A 109: 17507–17512.

48. Hemmrich G, Khalturin K, Boehm AM, Puchert M, Anton-Erxleben F, Wittlieb J, Klostermeier UC, Rosenstiel P, Oberg HH, Domazet-Loso T, et al. 2012. Molecular Signatures of the Three Stem Cell Lineages in Hydra and the Emergence of Stem Cell Function at the Base of Multicellularity. Mol Biol Evol 29: 3267–3280. http://mbe.oxfordjournals.org/content/29/11/3267.full.pdf.

49. Hobmayer B, Rentzsch F, Kuhn K, Happel CM, von Laue CC, Snyder P, Rothbacher U, Holstein TW. 2000. WNT signalling molecules act in axis formation in the diploblastic metazoan Hydra. Nature 407: 186–189. https://www.ncbi.nlm.nih.gov/pubmed/11001056.

50. Hoencamp C, Dudchenko O, Elbatsh AMO, Brahmachari S, Raaijmakers JA, van Schaik T, Cacciatore ÁS, Contessoto VG, van Heesbeen RGHP, van den Broek B, et al. 2021. 3D genomics across the tree of life reveals condensin II as a determinant of architecture type. Science (80-) 372: 984–989.

51. Inoue C, Bae SK, Takatsuka K, Inoue T, Bessho Y, Kageyama R. 2001. Math6, a bHLH gene expressed in the developing nervous system, regulates neuronal versus glial differentiation. Genes to Cells 6: 977–986.

52. Juliano CE, Swartz SZ, Wessel GM. 2010. A conserved germline multipotency program. Development 137: 4113–4126. http://www.ncbi.nlm.nih.gov/pubmed/21098563.

53. Jung SH, Ohyanagi H, Hayakawa S, Osato N, Nishimiya-Fujisawa C, Ikeo K, David CN, Fujisawa T, Gojobori T. 2007. The evolutionary emergence of cell type-specific genes inferred from the gene expression analysis of Hydra. Proc Natl Acad Sci U S A 104: 14735–14740.

54. Kaya-Okur HS, Wu SJ, Codomo CA, Pledger ES, Bryson TD, Henikoff JG, Ahmad K, Henikoff S. 2019. CUT&Tag for efficient epigenomic profiling of small samples and single cells. Nat Commun 10: 1–10.

55. Khalturin K, Anton-Erxleben F, Sassmann S, Wittlieb J, Hemmrich G, Bosch TCG. 2008. A novel gene family controls species-specific morphological traits in Hydra. PLoS Biol 6: 2436–2449.

56. Khalturin K, Hemmrich G, Fraune S, Augustin R, Bosch TC. 2009. More than just orphans: are taxonomically-restricted genes important in evolution? Trends Genet 25: 404–413. https://www.ncbi.nlm.nih.gov/pubmed/19716618.

57. Klimovich A, Wittlieb J, Bosch TCG. 2019. Transgenesis in Hydra to characterize gene function and visualize cell behavior. Nat Protoc 14: 2069–2090. http://dx.doi.org/10.1038/s41596-019-0173-3.

58. Kolmogorov M, Yuan J, Lin Y, Pevzner PA. 2019. Assembly of long, error-prone reads using repeat graphs. Nat Biotechnol 37: 540–546. http://dx.doi.org/10.1038/s41587-019-0072-8.

59. Koren S, Walenz BP, Berlin K, Miller JR, Bergman NH, Phillippy AM. 2017. Canu: Scalable and accurate long-read assembly via adaptive κ-mer weighting and repeat separation. Genome Res 27: 722–736.

60. Kotliar D, Veres A, Nagy MA, Tabrizi S, Hodis E, Melton DA, Sabeti PC. 2019. Identifying gene expression programs of cell-type identity and cellular activity with single-cell RNA-Seq. Elife 8: 1–26.

61. Langmead B, Salzberg SL. 2012. Fast gapped-read alignment with Bowtie 2. Nat Methods 9: 357–359. https://www.ncbi.nlm.nih.gov/pubmed/22388286.

62. Leclère L, Horin C, Chevalier S, Lapébie P, Dru P, Peron S, Jager M, Condamine T, Pottin K, Romano S, et al. 2019. The genome of the jellyfish Clytia hemisphaerica and the evolution of the cnidarian life-cycle. Nat Ecol Evol 3: 801–810. http://dx.doi.org/10.1038/s41559-019-0833-2.

63. Lengfeld T, Watanabe H, Simakov O, Lindgens D, Gee L, Law L, Schmidt HA, Ozbek S, Bode H, Holstein TW. 2009. Multiple Wnts are involved in Hydra organizer formation and regeneration. Dev Biol 330: 186–199. http://www.ncbi.nlm.nih.gov/pubmed/19217898.

64. Lenhoff HM, Brown RD. 1970. Mass culture of hydra: an improved method and its application to other aquatic invertebrates. Lab Anim 4: 139–154.

65. Liu Y, Kossack ME, Mcfaul ME, Christensen LN, Siebert S, Wyatt SR, Kamei C, Horst S, Arroy N, Drummond I, et al. 2021. Single-cell transcriptome reveals insights into the development and function of the zebrafish ovary. bioRxiv 2021.12.01.470669. http://biorxiv.org/content/early/2021/12/01/2021.12.01.470669.abstract http://biorxiv.org/content/early/2021/12/03/2021.12.01.470669.abstract.

66. Martin VJ, Littlefield CL, Archer WE, Bode HR. 1997. Embryogenesis in Hydra. Biol Bull 192: 345–363. https://www.journals.uchicago.edu/doi/10.2307/1542745.

67. Martinez DE. 1998. Mortality patterns suggest lack of senescence in hydra. Exp Gerontol 33: 217–225. http://www.ncbi.nlm.nih.gov/pubmed/9615920.

68. Martínez DE, Iñiguez AR, Percell KM, Willner JB, Signorovitch J, Campbell RD. 2010. Phylogeny and biogeography of Hydra (Cnidaria: Hydridae) using mitochondrial and nuclear DNA sequences. Mol Phylogenet Evol 57: 403–410.

69. McArthur E, Capra JA. 2021. Topologically associating domain boundaries that are stable across diverse cell types are evolutionarily constrained and enriched for heritability. Am J Hum Genet 108: 269–283. https://doi.org/10.1016/j.ajhg.2021.01.001.

70. McInnes L, Healy J, Melville J. 2018. UMAP: Uniform Manifold Approximation and Projection for Dimension Reduction. http://arxiv.org/abs/1802.03426.

71. Meers MP, Tenenbaum D, Henikoff S. 2019. Peak calling by Sparse Enrichment Analysis for CUT&RUN chromatin profiling. Epigenetics and Chromatin 12: 1–11. https://doi.org/10.1186/s13072-019-0287-4.

72. Münder S, Käsbauer T, Prexl A, Aufschnaiter R, Zhang X, Towb P, Böttger A. 2010. Notch signalling defines critical boundary during budding in Hydra. Dev Biol 344: 331–345. http://dx.doi.org/10.1016/j.ydbio.2010.05.517.

73. Murad R, Macias-Muñoz A, Wong A, Ma X, Mortazavi A. 2021. Coordinated Gene Expression and Chromatin Regulation during Hydra Head Regeneration. Genome Biol Evol 13: 1–17.

74. Musser JM, Schippers KJ, Nickel M, Mizzon G, Kohn AB, Pape C, Ronchi P, Papadopoulos N, Tarashansky AJ, Hammel JU, et al. 2021. Profiling cellular diversity in sponges informs animal cell type and nervous system evolution. Science (80-) 374: 717–723.

75. Piraino S, Boero F, Aeschbach B, Schmid V. 1996. Reversing the life cycle: Medusae transforming into polyps and cell transdifferentiation in Turritopsis nutricula (Cnidaria, Hydrozoa). Biol Bull 190: 302–312.

76. Putnam NH, Srivastava M, Hellsten U, Dirks B, Chapman J, Salamov A, Terry A, Shapiro H, Lindquist E, Kapitonov V V., et al. 2007. Sea anemone genome reveals ancestral eumetazoan gene repertoire and genomic organization. Science (80-) 317: 86–94.

77. Ramírez F, Bhardwaj V, Arrigoni L, Lam KC, Grüning BA, Villaveces J, Habermann B, Akhtar A, Manke T. 2018. High-resolution TADs reveal DNA sequences underlying genome organization in flies. Nat Commun 9. http://dx.doi.org/10.1038/s41467-017-02525-w.

78. Rao SS, Huntley MH, Durand NC, Stamenova EK, Bochkov ID, Robinson JT, Sanborn AL, Machol I, Omer AD, Lander ES, et al. 2014. A 3D map of the human genome at kilobase resolution reveals principles of chromatin looping. Cell 159: 1665–1680. http://www.ncbi.nlm.nih.gov/pubmed/25497547.

79. Reddy PC, Gungi A, Ubhe S, Galande S. 2020. Epigenomic landscape of enhancer elements during Hydra head organizer formation. Epigenetics and Chromatin 13: 1–16. https://doi.org/10.1186/s13072-020-00364-6.

80. Richards GS, Rentzsch F. 2015. Regulation of Nematostella neural progenitors by SoxB, Notch and bHLH genes. Dev 142: 3332–3342.

81. Robert NSM, Sarigol F, Zimmermann B, Meyer A, Voolstra CR, Simakov O. 2022. Emergence of distinct syntenic density regimes is associated with early metazoan genomic transitions. BMC Genomics 23: 1–14. https://doi.org/10.1186/s12864-022-08304-2.

82. Schaible R, Scheuerlein A, Danko MJ, Gampe J, Martinez DE, Vaupel JW. 2015. Constant mortality and fertility over age in Hydra. Proc Natl Acad Sci U S A 112: 15701–15706. http://www.ncbi.nlm.nih.gov/pubmed/26644561.

83. Schenkelaars Q, Perez-Cortes D, Perruchoud C, Galliot B. 2020. The polymorphism of Hydra microsatellite sequences provides strain-specific signatures. PLoS One 15: 1–20. http://dx.doi.org/10.1371/journal.pone.0230547.

84. Schwaiger M, Schonauer A, Rendeiro AF, Pribitzer C, Schauer A, Gilles AF, Schinko JB, Renfer E, Fredman D, Technau U. 2014. Evolutionary conservation of the eumetazoan gene regulatory landscape. Genome Res 24: 639–650. https://www.ncbi.nlm.nih.gov/pubmed/24642862.

85. Schwentner M, Bosch TCG. 2015. Revisiting the age, evolutionary history and species level diversity of the genus Hydra (Cnidaria: Hydrozoa). Mol Phylogenet Evol 91: 41–55. http://dx.doi.org/10.1016/j.ympev.2015.05.013.

86. Siebert S, Farrell JA, Cazet JF, Abeykoon Y, Primack AS, Schnitzler CE, Juliano CE. 2019. Stem cell differentiation trajectories in Hydra resolved at single-cell resolution. Science (80-) 365: eaav9314. http://www.sciencemag.org/lookup/doi/10.1126/science.aav9314.

87. Simakov O, Bredeson J, Berkoff K, Marletaz F, Mitros T, Schultz DT, O’Connell BL, Dear P, Martinez DE, Steele RE, et al. 2022. Deeply conserved synteny and the evolution of metazoan chromosomes. Sci Adv 8. https://www.science.org/doi/10.1126/sciadv.abi5884.

88. Slater GSC, Birney E. 2005. Automated generation of heuristics for biological sequence comparison. BMC Bioinformatics 6: 1–11.

89. Smith CL, Mayorova TD. 2019. Insights into the evolution of digestive systems from studies of Trichoplax adhaerens. Cell Tissue Res 377: 353–367.

90. Sogabe S, Hatleberg WL, Kocot KM, Say TE, Stoupin D, Roper KE, Fernandez-Valverde SL, Degnan SM, Degnan BM. 2019. Pluripotency and the origin of animal multicellularity. Nature 570: 519–522. http://dx.doi.org/10.1038/s41586-019-1290-4.

91. Stuart T, Butler A, Hoffman P, Hafemeister C, Papalexi E, Mauck WM, Hao Y, Stoeckius M, Smibert P, Satija R. 2019. Comprehensive Integration of Single-Cell Data. Cell 177: 1888–1902.e21. https://doi.org/10.1016/j.cell.2019.05.031.

92. Tagle DA, Koop BF, Goodman M, Slightom JL, Hess DL, Jones RT. 1988. Embryonic ε and γ globin genes of a prosimian primate (Galago crassicaudatus). Nucleotide and amino acid sequences, developmental regulation and phylogenetic footprints. J Mol Biol 203: 439–455.

93. Tarashansky AJ, Musser JM, Khariton M, Li P, Arendt D, Quake SR, Wang B. 2021. Mapping single-cell atlases throughout metazoa unravels cell type evolution. Elife 10: 1–24.

94. Technau U, Rudd S, Maxwell P, Gordon PMK, Saina M, Grasso LC, Hayward DC, Sensen CW, Saint R, Holstein TW, et al. 2005. Maintenance of ancestral complexity and non-metazoan genes in two basal cnidarians. Trends Genet 21: 633–639.

95. Tournière O, Dolan D, Richards GS, Sunagar K, Columbus-Shenkar YY, Moran Y, Rentzsch F. 2020. NvPOU4/Brain3 Functions as a Terminal Selector Gene in the Nervous System of the Cnidarian Nematostella vectensis. Cell Rep 30: 4473–4489.e5.

96. Trembley A. 1744. Mémoires, pour servir à l’histoire d’un genre de polypes d’eau douce, à bras en forme de cornes. Chez Jean and Herman Verbeek.

97. Vogg MC, Beccari L, Iglesias Ollé L, Rampon C, Vriz S, Perruchoud C, Wenger Y, Galliot B. 2019. An evolutionarily-conserved Wnt3/β-catenin/Sp5 feedback loop restricts head organizer activity in Hydra. Nat Commun 10: 312. http://dx.doi.org/10.1038/s41467-018-08242-2.

98. Vogg MC, Ferenc J, Buzgariu WC, Perruchoud C, Papasaikas P, Sanchez PGL, Nuninger C, Delucinge-Vivier C, Rampon C, Beccari L, et al. 2021. The transcription factor Zic4 acts as a transdifferentiation switch. bioRxiv 2021.12.22.473838. http://biorxiv.org/content/early/2021/12/23/2021.12.22.473838.abstract.

99. Waltman L, Van Eck NJ. 2013. A smart local moving algorithm for large-scale modularity-based community detection. Eur Phys J B 86.

100. Weimer BR. 1928. The Physiological Gradients in Hydra I. Reconstitution and Budding in Relation to Length of Piece and Body Level in Pelmatohydra oligactis. Physiol Zool 1: 183–230. http://www.jstor.org/stable/30151045.

101. Wittlieb J, Khalturin K, Lohmann JU, Anton-Erxleben F, Bosch TC. 2006. Transgenic Hydra allow in vivo tracking of individual stem cells during morphogenesis. Proc Natl Acad Sci U S A 103: 6208–6211. https://www.ncbi.nlm.nih.gov/pubmed/16556723.

102. Wong WY, Simakov O, Bridge DM, Cartwright P, Bellantuono AJ, Kuhn A, Holstein TW, David CN, Steele RE, Martínez DE. 2019. Expansion of a single transposable element family is associated with genome-size increase and radiation in the genus Hydra. Proc Natl Acad Sci U S A 116: 22915–22917.

103. Zheng H, Xie W. 2019. The role of 3D genome organization in development and cell differentiation. Nat Rev Mol Cell Biol 20: 535–550. http://dx.doi.org/10.1038/s41580-019-0132-4.

104. Zimmermann B, Robb SM, Genikhovich G, Fropf WJ, Weilguny L, He S, Chen S, Lovegrove-Walsh J, Hill EM, Ragkousi K, et al. 2020. Sea anemone genomes reveal ancestral metazoan chromosomal macrosynteny. bioRxiv 2020.10.30.359448. https://doi.org/10.1101/2020.10.30.359448.

