## Supplementary material for "New *Hydra* genomes reveal conserved principles of hydrozoan transcriptional regulation": Cazet_Supplement

**Table of Contents:**

Supplemental Materials and Methods

Figures S1 to S19

Tables S1 to S2

Legends for Supplementary Data S1 to S15

References

### Supplemental Materials and Methods

#### Data Availability

We have generated a new genome portal, available at research.nhgri.nih.gov/HydraAEP/, that allows users to interact with and download the data generated in this study. A BLAST server is available to search for genes of interest in the *H. oligactis* and strain AEP *H. vulgaris* gene models. The portal includes an interactive genome browser for visualizing gene models, repetitive regions, ATAC-seq and CUT&Tag peaks, ATAC-seq and CUT&Tag read density, and sequence conservation across the AEP assembly. The website also features an interactive ShinyCell portal (Ouyang et al. 2021) for viewing the AEP-aligned *Hydra* single-cell atlas.

Step-by-step descriptions of all computational analyses conducted as part of this study, including all relevant code, formatted both as markdown and HTML documents are available at github.com/cejuliano/brown_hydra_genomes.

The raw sequencing data and assembled genomic sequences generated by this study has been deposited to NCBI under the BioProject ID PRJNA816482. We have also made all raw sequencing reads, scripts, and processed data files associated with this study available for download through the genome portal at research.nhgri.nih.gov/HydraAEP/download/index.cgi?dl=fa.

Importantly, due to data loss, we no longer have access to the basecall quality scores for the PacBio sequencing data. Because SRA requires that all submitted sequencing data include quality scores, we were unable to upload the PacBio data to NCBI. However, the PacBio data is available at research.nhgri.nih.gov/HydraAEP/download/index.cgi?dl=fa, and the basecall quality scores are not necessary for fully reproducing the results presented in this study.

#### ***Hydra* strains and animal care**

All *Hydra* strains were cultured using standard methods (Lenhoff and Brown 1970). The AEP strain of *H. vulgaris* was generated from a cross between the PA1 strain isolated by Dr. Carolyn Teragawa from a pond on the Haverford College campus near Philadelphia, Pennsylvania and the CA7 strain isolated by Drs. Lynne Littlefield and Carolyn Teragawa at Boulder Creek, near Susanville, California (Martin et al. 1997). The DNA used for generating the strain AEP *H. vulgaris* assembly was isolated from a clonally propagated line (the “Kiel” AEP line; courtesy of Thomas Bosch) that was generated from a self-cross of the original AEP line. The DNA used for generating the *H. oligactis* assembly was isolated from the Innsbruck female12 strain, a clonally propagated line originating from a single polyp collected from Lake Piburger See in Tyrol, Austria.

In addition to the Kiel AEP strain, the following lines were used for generating RNA-seq libraries: a transgenic line with an actin::EGFP transgene integrated into the ectodermal lineage and an actin::DsRed2 transgene integrated into the endodermal lineage (“watermelon” line)

(Glauber et al. 2015), a transgenic line with an actin::DsRed2 transgene integrated into the ectodermal lineage and an actin::EGFP transgene integrated into the endodermal lineage (" inverse watermelon" line) (Glauber et al. 2015), a transgenic line with an EF1α::EGFP transgene integrated into the endodermal lineage ("enGreen1" line; courtesy of Rob Steele and Catherine Dana), and a transgenic line with a transgene containing EGFP and DsRed2 in an operon configuration with expression driven by the actin promoter integrated into the ectodermal lineage ("operon" line) (Dana et al. 2012).

***Hydra vulgaris* strain AEP genome sequencing**

To generate high molecular weight (HMW) genomic DNA (gDNA) libraries for sequencing and assembling the strain AEP *H. vulgaris* genome, we used thirty whole adult polyps from a clonally propagated population belonging to the Kiel AEP line as input. The tissue was flash frozen in liquid nitrogen and HMW gDNA was purified using a Qiagen Gentra Puregene kit following standard manufactures instructions for mouse tail tissue (Qiagen Cat # 158445; Hilden, Germany). We then performed a Phenol/Chloroform purification using 5PRIME Phase Lock Gels (Quantabio Cat # 2302830; Beverly, Massachusetts) and precipitated the DNA by adding 0.4X 5M ammonium acetate and 3X ice cold ethanol. The DNA pellet was washed twice with 70% ethanol and resuspended in elution buffer (10mM Tris, pH 8.0). We used a Pippin Pulse gel electrophoresis system (Sage Sciences, Beverly, MA) to verify the DNA integrity and a NanoDrop spectrophotometer (ThermoFisher Scientific, Waltham, Massachusetts) to verity the DNA purity.

To generate the Oxford Nanopore library, HMW gDNA was gently sheared to 70kb-100kb using a Megaruptor 2 (Diagenode Cat # B06010002; Denville, New Jersey) and the library was prepared using the Oxford Nanopore Ligation Sequencing Kit (Oxford Nanopore Technologies Cat # LSK-109; Oxford, United Kingdom) following standard manufacturer's instructions except for extended incubation times for DNA damage repair, end repair, ligation, and bead elution. 850ng of the final library was loaded on PromethION R9.4.1 flow cells and the data were collected for sixty-four hours. Basecalling was performed live during the run with guppy v1.8.1.

A HMW gDNA PacBio library was generated using a SMRTbell *Express* Template Prep Kit 2.0 (PacBio Cat # 100-938-900; Menlo Park, California) following standard manufacturer's instructions. The library was then sequenced on a PacBio Sequel II sequencer using a 1M v3 SMRT Cell (PacBio Cat # 101-531-000).

To generate the 10X chromium library, HMW gDNA was loaded onto a Chromium Genome Chip (10X Genomics Cat # 120257; Pleasanton, California) and the library was prepared using Chromium Genome Library & Gel Bead Kit v.2 (10X Genomics Cat # 120258) and Chromium Controller (10X Genomics Cat # 120270) according to manufacturer's instructions with one modification. Briefly, gDNA was combined with Master Mix, Genome Gel Beads, and partitioning oil to create Gel Bead-in-Emulsions (GEMs) on a Chromium Genome Chip. The GEMs were isothermally amplified and barcoded DNA fragments were recovered for Illumina library construction. The post-GEM DNA was quantified using a Bioanalyzer 2100 with an Agilent High sensitivity DNA kit (Agilent Cat # 5067-4626; Santa Clara, California). Prior to Illumina library construction, the GEM amplification product was sheared on an E220 Focused-Ultrasonicator (Covaris Cat # 500239; Woburn, MA) to approximately 375 bp (50 seconds at peak power = 175, duty factor = 10, and cycle/burst = 200). Then, the sheared GEMs were converted to a sequencing library following the 10X standard operating procedure. The library was quantified by qPCR with a Kapa Library Quant kit (Roche Cat # 07960140001; Basel, Switzerland) and sequenced on a HiSeqX10 (Illumina, San Diego, CA) using 2 x 150 bp reads.

For generating the Hi-C library, we used 10 whole flash frozen adult polyps as input. The library was generated using the Arima Hi-C Kit (Arima Genomics Cat # A510008; San Diego, California) following the standard manufacturer's protocol for small animal tissue with the following modification: the frozen tissue was ground using a mortar and pestle for 1 minute in fixation buffer and was subsequently left for 19 minutes at room temperature. The proximally-ligated DNA was fragmented using Covaris E220 (Covaris Cat # 500239) and the biotinylated fragments were enriched. NGS library was prepared using KAPA Hyper prep kit (Roche Cat # 07962363001) and the library was sequenced on an Illumina NovaSeq 6000 using 2 x 150 bp reads.

**Whole-animal RNA-seq**

To aid in annotating and benchmarking our AEP genome assembly, we generated and sequenced several whole-animal RNA-seq libraries using multiple strain AEP-derived lines. In total, there were 13 libraries: one from the watermelon line, one from the inverse watermelon line, one from the enGreen1 line, one from the operon line, three from male Kiel AEP polyps, three from female Kiel AEP polyps, and three from Kiel AEP polyps that were not producing gametes.

For the watermelon, inverse watermelon, enGreen1, and operon RNA-seq libraries, total RNA was purified using a standard Trizol extraction protocol. RNA-seq libraries were then prepared using a TruSeq stranded mRNA kit (Illumina Cat # RS-122-2201) according to the manufacturer's recommended protocol with the following modifications: the RNA was sheared for only 1.5 minutes and the resulting fragments were size selected using a LabChip XT DNA 750 (PerkinElmer Cat # 760541; Waltham, Massachusetts) to be ~500 bp prior to the final PCR enrichment step. The libraries were then sequenced on an Illumina HiSeq2000 using 2 x 100 bp reads.

For the Kiel AEP libraries, total RNA was purified using a standard Trizol extraction protocol. Contaminating DNA was then removed by performing a DNAse digest using the QIAGEN DNAse set (QIAGEN Cat # 79254). A final purification was then performed using the Zymogen RNA Clean and Concentrator Kit (Zymo Research Cat # R1017; Irvine, California) according to the standard manufacturer's protocol. RNA-seq libraries were then generated using the Kapa mRNA-seq Hyper kit (Kapa Biosystems Cat # KK8581; Kapa Biosystems, Cape Town, South Africa). The libraries were then sequenced on a HiSeq4000 using 1 x 50 bp reads. We also performed additional sequencing for one biological replicate from both the male and female Kiel AEP libraries, which were sequenced on an Illumina HiSeq4000 using 2 x 150 bp reads.

To perform the alignment benchmarking analysis presented in Fig S5, the single end Kiel AEP RNA-seq reads were first processed with Trimmomatic (Bolger et al. 2014) to remove stretches of low-quality base-calls and contaminating adapter sequence. The data was then aligned to both the strain AEP and strain 105 *H. vulgaris* genome assemblies using the RSEM (Li and Dewey 2011) implementation of STAR (Dobin et al. 2013). The code for this alignment benchmarking analysis is included in the supplemental file 03_aepGenomeAnnotation.md.

#### *Hydra vulgaris* strain AEP genome assembly

A step-by-step description of the strain AEP *H. vulgaris* genome assembly methodology, including all relevant code, is provided in the markdown document *01_aepGenomeAssembly* available at github.com/cejuliano/brown_hydra_genomes.

The initial draft assembly was generated from the Oxford Nanopore data using Canu (Koren et al. 2017). We then mapped the 10X linked-read data to the draft genome and polished the assembly using Pilon (Walker et al. 2014). For this and all subsequent steps involving the 10X data, we used the 10X Long Ranger pipeline for genome alignment. Following the polishing step, we cut contigs in predicted mis-assembled regions with Tigmint (Jackman et al. 2018) using the 10X data. We then used the 10X data to identify and collapse duplicated contigs in the assembly using Purge Haplotigs (Roach et al. 2018). Deduplicated contigs were scaffolded with ARCS (Yeo et al. 2018) using the 10X data, and gaps introduced by the scaffolding were filled with PBJelly (English et al. 2012) using the Oxford Nanopore and PacBio data. To generate pseudo-chromosome scaffolds, we aligned the Hi-C data using Juicer (Durand et al. 2016) and scaffolded the assembly using the 3d-dna pipeline (Dudchenko et al. 2017). This was followed by an additional gap-filling step with PBJelly using the Oxford Nanopore and PacBio data. To finalize the assembly sequence, we performed another round of Pilon error correction using the 10X, PacBio, and Oxford Nanopore data. Minimap2 (Li 2018) was used for aligning the long-read data to the genome for the Pilon correction.

The resulting assembly is 901 Mb in length and contains 15 pseudo-chromosome scaffolds, consistent with the haploid chromosome number in *Hydra* (Rahat et al. 1985; Zacharias et al. 2004). Like the strain 105 *H. vulgaris* genome assembly, the AEP assembly is roughly 20-25% smaller than empirical genome size estimates (~1.06-1.22 Gb for the AEP strain) (Chapman et al. 2010; Zacharias et al. 2004), which is likely due to intrinsic difficulties in resolving long and repetitive stretches of heterochromatin. Nonetheless, the contiguity and completeness of the AEP assembly is comparable to the best currently available hydrozoan genomes (Fig. 1B and Table S1). Compared to the recently updated chromosome-level assembly of the strain 105 *H. vulgaris* genome (Simakov et al. 2022), the AEP assembly contains ~10% more sequence (900.9 Mb, compared to 819.4 Mb in the 105 v3 assembly) and a similar number of intact single-copy orthologs predicted from genomic sequence using BUSCO (866, compared to 862 in the 105 v3 assembly; Table S1).

**Genome repeat annotation**

A step-by-step description of the repeat annotation methodology used for the *H. oligactis* and *H. vulgaris* genomes, including all relevant code, is provided in the markdown document *02_repeatMasking* available at github.com/cejuliano/brown_hydra_genomes.

To compensate for the lack of well-annotated repeat families available for *Hydra*, we used RepeatModeler2 (Flynn et al. 2020) to predict repeat families ab initio for the *H. oligactis* and strain AEP *H. vulgaris* genome assemblies. We used RepeatMasker (repeatmasker.org) to identify repetitive regions in the strain AEP and strain 105 *H. vulgaris* genome assemblies as well as the *H. oligactis* assembly. For masking repeats in the strain AEP and strain 105 *H. vulgaris* genome assemblies, we used both the strain AEP *H. vulgaris* RepeatModeler2 repeats as well as the Dfam eumetazoan repeat database as repeat libraries when running RepeatMasker. For masking repeats in the *H. oligactis* genome assembly, we used both the *H. oligactis* RepeatModeler2 repeats as well as the Dfam eumetazoan repeat database as repeat libraries when running RepeatMasker. We then used utility scripts included with RepeatMasker to calculate sequence divergence for predicted repeat instances and to generate the repeat landscape plots presented in Fig S1.

Consistent with previous characterizations of brown *Hydra* genomes, we find that the AEP genome is highly A/T rich (~72%) and repetitive (Wong et al. 2019; Chapman et al. 2010). We estimate that ~71% of the AEP genome is repetitive, with ~6% being simple/low-complexity repeats and ~65% originating from transposable elements (TEs) (Fig. S1A-C). These estimates are slightly higher than the strain 105 genome (~57% TEs and Fig. S1D-F) (Chapman et al.

2010). As with the 105 strain, class II TEs—particularly the hAT, CMC, and Mariner families—make up most TE sequences in the AEP genome, although a sizable minority are derived from L2 and CR1 LINE retrotransposons (Fig. S1A).

#### *Hydra vulgaris* strain AEP genome gene annotation

A step-by-step description of the strain AEP *H. vulgaris* genome gene annotation methodology, including all relevant code, is provided in the markdown document *03_aepGenomeAnnotation* available at github.com/cejuliano/brown_hydra_genomes.

We generated an initial set of gene models for the strain AEP *H. vulgaris* genome using the BRAKER2 gene prediction pipeline (Brůna et al. 2021). As input into the pipeline, we included the AEP genome sequence with all repetitive regions soft-masked, a custom database of metazoan proteomes, and a whole-animal RNA-seq dataset (described in the "Whole-animal RNA-seq" section above) that was aligned to the soft-masked genome using STAR (Dobin et al. 2013). To supplement the BRAKER2 predictions, we designed a custom annotation pipeline that used exonerate (Slater and Birney 2005) to generate gene models using transcript sequences from a previously published transcriptome (Siebert et al. 2019) and a manually curated database of *Hydra* transcript sequences from GenBank. We collapsed duplicated/overlapping gene models in the combined BRAKER2 and exonerate gene predictions by selecting the gene model that had the highest alignment score following a BLAST search against the same custom protein database that was used to generate the BRAKER2 predictions. We then filtered out all gene models that had interrupted reading frames, were shorter than 50 amino acids, or were predicted by InterProScan (Blum et al. 2021; Jones et al. 2014) to contain one or more transposase domains. To improve UTR and splice isoform annotations in our gene predictions, we used the Trinity genome-guided assembly pipeline (Grabherr et al. 2011) to generate a transcriptome from the genome-aligned whole-animal RNA-seq data that was originally used as input for the BRAKER2 pipeline. We aligned this transcriptome to the AEP assembly and used this alignment to update the merged exonerate and BRAKER2 gene models with PASA (Haas et al. 2003), resulting in the final set of gene predictions presented in this study.

Our AEP annotation pipeline identified 28,917 protein coding genes that encode 37,784 predicted transcripts. Although the total gene number is ~14% lower than that observed in the 105 assembly annotations, the AEP annotation contains ~12% more complete single-copy orthologs as predicted using BUSCO (Fig. 1B and Table S1), demonstrating an improvement in both accuracy and sensitivity. Furthermore, the AEP assembly gene predictions are the first *H. vulgaris* gene models to include UTRs, with ~48% (13,901) of gene models containing 5' UTRs and ~46% (13,183) containing 3' UTRs. Overall, the AEP gene predictions are comparable to our previously published AEP transcriptome in both the number of predicted transcripts and the number of complete single-copy orthologs (Table S1) (Siebert et al. 2019), suggesting that our gene annotations have largely captured the transcriptomic repertoire of *H. vulgaris*.

To generate functional annotations for the AEP gene models, we performed a BLAST search against the UniProt protein database (Bateman et al. 2021), predicted protein domains using InterProScan, and identified orthologs in 43 other metazoans using OrthoFinder (Emms and Kelly 2019). The combined results from these annotation analyses are included in Supplemental Data *S*1. To identify putative TFs in the AEP gene models, we filtered the InterProScan predictions using a custom set of keywords and GO terms related to transcriptional regulation and DNA binding activity (see 03_aepGenomeAnnotation.md for details; gene IDs of putative TFs listed in Supplemental Data S1).

#### ***Hydra oligactis* genome sequencing**

For generating a draft genome for *H. oligactis*, we prepared two HMW gDNA libraries using the Innsbruck female12 strain of *H. oligactis*. For the first library, HMW gDNA was extracted from 10 whole adult polyps using the Circulomics NanoBind BigTissue kit (Circulomics Cat # NB-900-701-01; Baltimore, Maryland) according to the manufacturer's "Dounce" protocol (Circulomics document # EXT-DHH-001) with the following modifications: we used intact animals instead of finely minced tissue, we homogenized the tissue in 500 µl Buffer CT instead of 750 µl, animals were homogenized using a pestle in a 1.5 ml microcentrifuge tube for 2 minutes instead of using a dounce homogenizer, and the homogenate was pelleted at 1500 G instead of 3000 G. We removed short DNA using the Short Read Eliminator (Circulomics Cat # SS-100-101-01) and Short Read Eliminator XS (Circulomics Cat # SS-100-121-01) kits according to the manufacturer's standard protocol and eluted the samples overnight. We prepared the sequencing library using the Oxford Nanopore Ligation Sequencing Kit (Oxford Nanopore Technologies Cat # LSK-109) according to the standard manufacturer's protocol with the modification that the first two 5-minute incubations were extended to be 30 minutes each. The final library was eluted in 26 µl elution buffer and the library was loaded twice onto an Oxford Nanopore MinION sequencer, with DNAse from the Flow Cell Wash Kit (Oxford Nanopore Technologies Cat # EXP-WSH003) being used to remove gDNA carryover between runs.

The second HMW gDNA library was generated as described above with a few modifications. First, 100 instead of 10 whole animals were used as input. We also made additional modifications to the NanoBind protocol. We prolonged the proteinase K digestion from 30 minutes to 150 minutes, adding another 10 µl proteinase K and another 75 µl Buffer CLE3 90 minutes into the digestion. We also used 30 µl of RNAse A instead of 20 µl. Instead of using a Nanobind disk for DNA extraction as described in the standard protocol, we used the following approach: the lysate was centrifuged at 10,000 G for 5 minutes at room temperature, the resulting pellet was washed with 400 µl Buffer CW1 and centrifuged at 10,000 G for 5 minutes, the pellet was then washed with 500 µl Buffer CW2 and centrifuged at 10,000 G for 5 minutes, the supernatant was removed and the pellet air-dried for 1 minute, and DNA was eluted in 70 µl Elution Buffer. Short gDNA fragment elimination and library preparation was performed as described for the first library. The library was eluted in 60 µl and was loaded onto the MinION sequencer a total of five times. The total coverage of all sequencing libraries was ~17X (2.4 million reads with an N50 of 22.7 kb).

***Hydra oligactis* assembly and annotation**

A step-by-step description of the *H. oligactis* genome assembly and gene annotation methodology, including all relevant code, is provided in the markdown document *04_oligactisDraftGenome* available at github.com/cejuliano/brown_hydra_genomes.

We generated an initial draft assembly for *H. oligactis* with Flye (Kolmogorov et al. 2019) using reads from the two combined Oxford Nanopore libraries described above. The errors in the assembly were then polished with Medaka (github.com/nanoporetech/medaka) using the Nanopore data. To generate a preliminary set of gene models for the draft assembly, we first used previously published whole-animal RNA-seq data from *H. oligactis* (Sun et al. 2020; Rathje et al. 2020) to generate a de novo transcriptome using Trinity (Grabherr et al. 2011). We then aligned this transcriptome to a repeat-masked version of the *H. oligactis* draft genome using minimap2 (Li 2018). Finally, we ran the BRAKER2 gene prediction pipeline (Brůna et al. 2021), providing as input the repeat-masked *H. oligactis* genome sequence and the genome-mapped Trinity transcriptome.

The *oligactis* assembly is 1274 Mb in length, or ~88% of the empirically estimated genome size (Zacharias et al. 2004). The assembly is ~51-fold more contiguous than the previously available draft genome for *H. oligactis* (N50 of 274.9 kb, compared to previous N50 of 5.4 kb) and has ~27-fold fewer total contigs (16,314 contigs, compared to 447,335 contigs in the previous assembly; Fig 1B) (Vogg et al. 2019). The new *H. oligactis* draft genome is also more complete, with nearly double the number of intact single-copy orthologs (841, compared to 444 in the previous assembly) (Table S1). The A/T and repeat composition (~72% and ~74% respectively) were similar to *H. vulgaris*, although the *H. oligactis* assembly had a slightly higher abundance of repetitive elements (Fig. 1B and Fig. S1*G-I*). We identified 60,590 genes, which is likely an over-estimation of the genome's genic content given that hydrozoan genomes typically contain between 20,000 and 30,000 genes (Leclère et al. 2019; Hamada et al. 2020; Chapman et al. 2010). Nonetheless, the high BUSCO completeness of these gene models (86.2%) suggests that they accurately capture most of the genic content of the *H. oligactis* genome. Thus, we present the first annotated draft genome of *H. oligactis* that is of comparable quality to other published hydrozoan genomes and suitable for systematic comparative analyses.

**ATAC-seq**

Whole animal ATAC-seq was performed in triplicate on adult bud-free strain AEP *H. vulgaris* polyps using a previously described protocol (Corces et al. 2017; Siebert et al. 2019). All steps of the ATAC-seq protocol were performed using chilled solutions on ice unless otherwise indicated. For each replicate, 5 whole bud-free adult polyps that had been starved for two days were transferred to a sterile 1.5 ml microcentrifuge tube and briefly washed with 1 ml of *Hydra* dissociation medium (DM) (3.6 mM KCl, 6 mM $CaCl_2$, 1.2 mM $MgSO_4$, 6 mM sodium citrate, 6 mM sodium pyruvate, 6 mM glucose, 12.5 mM TES buffer, adjusted to pH 6.9) (Gierer et al. 1972). The polyps were then homogenized in 1 ml DM using ~50 strokes of a tight-fitting glass dounce. The homogenate was transferred into a sterile 1.5 ml microcentrifuge and spun down at 500 G for 5 minutes in a centrifuge chilled to 4˚C. The cell pellet was resuspended in 50µl resuspension buffer (RSB) (10 mM Tris-HCl, 10 mM NaCl, 3 mM MgCl2, pH 7.4) containing 0.1% Tween-20, 0.1% NP-40, and 0.01% digitonin. Lysis proceeded for 3 minutes and was subsequently halted by adding 1 ml RSB containing 0.1% Tween-20. Nuclear density in the lysate was quantified by loading 19 µl of the resuspension and 1 ul of 20mM Hoechst 33342 (ThermoFisher Scientific Cat # 62249; Waltham, Massachusetts) onto a Fuchs-Rosenthal hemocytometer. An aliquot of the resuspended lysate containing ~50,000 nuclei was then transferred to a fresh 1.5 ml microcentrifuge tube and was subsequently spun down for 10 minutes at 500 G in a centrifuge chilled to 4˚C. The crude nuclear pellet was then resuspended in 50 ul tagmentation buffer (1X TD buffer [Illumina Cat # 20034197], 33% phosphate-buffered saline, 0.01% digitonin, 0.1% Tween-20, 5 ml TDE1 [Illumina Cat # 20034197]) and shaken at 1000 rpm for 30 min at 37˚C. Tagmentation was halted by adding 250 µl of PB buffer from a QIAGEN MinElute PCR Purification Kit (QIAGEN Cat # 28004; Hilden, Germany).

Tagmented DNA was purified using a QIAGEN MinElute PCR Purification Kit using the standard manufacturer's instructions. The libraries were eluted in 21 µl water and amplified for an initial five PCR cycles using 2X NEBNext master mix (NEB Cat # M0541S; Ipswitch, MA) following the cycling parameters specified in the original ATAC-seq protocol (Buenrostro et al.

2013, 2015). The number of additional PCR cycles following this initial amplification was then determined by performing qPCR on an aliquot of the pre-amplified libraries as described in the original ATAC-seq protocol. Biological replicate 1 received 1 additional cycle of PCR (for a total of 6), replicate 2 received 3 additional cycles (for a total of 8), and replicate 3 received 4 additional cycles (for a total of 9). Two rounds of post-PCR clean-up were performed using Agencourt AMPure XP beads (Beckman Coulter Cat # A63881; Pasadena, California) following the standard manufacturer's protocol. During this step we selected for DNA fragments between 100 and 700 bp in size. Library concentration was quantified using the Qubit dsDNA HS Assay Kit (ThermoFisher Scientific Cat # Q32851) and fragment size distributions were determined using the Bioanalyzer High-Sensitivity DNA kit (Agilent Cat # 5067-4626). The libraries were then pooled at roughly equimolar proportions and sequenced on an Illumina NextSeq 500 using 2 x 75 bp reads.

**CUT&Tag**

CUT&Tag targeting H3K4me1, H3K4me3, and H3K27me3 were each performed in triplicate using a modified version of the originally published CUT&Tag protocol (Kaya-Okur et al. 2019) that was adapted for use in *Hydra*. Each CUT&Tag replicate consisted of 40 whole, bud-free strain AEP *H. vulgaris* polyps that had been fed once weekly and then starved for two days prior to the experiment. Unless otherwise specified, all steps were performed at room temperature without agitation. The polyps were collected in a 1.5 ml microcentrifuge tube, washed once with 1 ml DM, and then homogenized in 1 ml DM using 40 strokes of a tight-fitting glass dounce. The homogenate was passed through a 70 µm filter and centrifuged for 5 minutes at 1000 G. The resulting pellet was resuspended in 1 ml of lysis buffer (20mM HEPES, pH 7.5, 150 mM NaCl, 0.5 mM spermidine, 1X cOmplete protease inhibitor [Roche Cat # 11836153001], 2 mM EDTA, 0.1% tween-20, 0.1% NP-40, and 0.01% digoxygenin) and incubated for 5 minutes. The lysate was centrifuged for 5 minutes at 1300 G to produce a crude nuclear pellet, which was then resuspended in 1 ml of wash buffer (20mM HEPES, pH 7.5, 150 mM NaCl, 0.5 mM spermidine, 1X cOmplete protease inhibitor) and divided evenly into 4 1.5 ml microcentrifuge tubes. The volume of each tube was then brought to 1 ml using wash buffer. 10 µl of 5mg/ml Concanavalin A coated magnetic beads (Bangs Laboratories Cat # BP531; Fishers, Indiana) that had first been washed twice in bead activation buffer (20 mM HEPES, pH 7.5, 10 mM KCl, 1 mM $CaCl_2$, and 1 mM $MnCl_2$) was added to each tube. The bead-nuclei suspensions were then incubated for 10 minutes on a rotator and the supernatant was subsequently removed using a magnet stand. Bead-bound nuclei were resuspended in 50 µl solutions of either 1:1000 negative control rabbit IgG (EpiCypher Cat # 13-0042; Durham, North Carolina), 1:100 rabbit α-H3K4me1 (Abcam Cat # ab8895; Cambridge, United Kingdom), 1:100 rabbit α-H3K4me3 (Active Motif Cat # 39060; Carlsbad, California), or 1:50 rabbit α-H3K27me3 (Cell Signaling Technology Cat # 9733T; Danvers, Massachusetts) diluted in antibody buffer (1% bovine serum albumin and 2 mM EDTA in wash buffer). The nuclei were incubated in the primary antibody solutions for 2 hours. This was followed by a 1-hour incubation in 50 µl of anti-rabbit secondary antibodies (EpiCypher Cat # 13-0047) diluted 1:100 in antibody buffer. The nuclei were then quickly washed three times in 1ml wash buffer, resuspended in 50 µl of 1x pAG-Tn5 (EpiCypher Cat # 15-1017) diluted in high-salt buffer (20mM HEPES, pH 7.5, 300 mM NaCl, 0.5 mM spermidine, 1X cOmplete protease inhibitor), and incubated for 1 hour. Next, excess pAG-Tn5 was removed using three quick 1 ml washes with high-salt buffer and the nuclei were resuspended in 150 µl of tagmentation buffer (high-salt buffer with 10 mM $MgCl_2$ added). Tagmentation was then allowed to proceed for 1 hour at 37˚C. Tagmentation was stopped by adding 5 µl 0.5 mM EDTA, 1.5 µl 10% SDS, and 2.5 µl proteinase K (ThermoFisher Scientific Cat # EO0492) to each sample and incubating at 55˚C for 1 hour.

Tagmented DNA was purified using a Zymogen Oligo Clean & Concentrator Kit (Zymo Research Cat # D4060; Irvine, California) following the standard manufacturer's protocol. The libraries were eluted in 21 µl water and amplified using 2X NEBNext master mix following the cycling parameters described in the original CUT&Tag protocol (Kaya-Okur et al. 2019) for a total of 13 cycles. We then used Agencourt AMPure XP beads to perform two rounds of post-PCR clean-up and to select for DNA fragment sizes between 100 and 700 base pairs. We quantified the concentration of our libraries using the Qubit dsDNA HS Assay Kit and we determined their fragment size distributions using the Bioanalyzer High-Sensitivity DNA kit. When measuring the concentrations of our purified libraries, we found that our negative control samples were too dilute to effectively validate their size and concentration for pooling. We therefore performed another five rounds of PCR amplification on the three negative control libraries followed by two additional rounds of AMPure bead cleanup. Finally, libraries were pooled at roughly equimolar concentrations and sequenced on an Illumina NextSeq 500 using 2 x 75 bp reads.

**Cis-regulatory element annotation**

A step-by-step description of the *Hydra* cis-regulatory element annotation methodology, including all relevant code, is provided in the markdown document *08_creIdentification* available at github.com/cejuliano/brown_hydra_genomes.

To analyze the ATAC-seq data collected from whole strain AEP *H. vulgaris* polyps, we first filtered the raw reads using Trimmomatic (Bolger et al. 2014) to remove stretches of low-quality base-calls and contaminating adapter sequence. The filtered reads were then aligned to the AEP assembly using Bowtie2 (Langmead and Salzberg 2012). To remove mitochondrial reads, we also aligned the ATAC-seq data to the *Hydra* mitochondrial genome (Voigt et al. 2008) and subsequently discarded any reads that aligned to the mitochondrial and nuclear genome references using Picard Tools (broadinstitute.github.io/picard/). We next identified and removed PCR duplicates from the aligned data using Samtools (Li et al. 2009) and Picard Tools. We then called peaks for each ATAC-seq biological replicate using MACS2 (Zhang et al. 2008). To generate a consensus peakset of biologically reproducible ATAC-seq peaks, we first calculated irreproducible discovery rate (IDR) (Li et al. 2011) peak scores for each pairwise combination of biological replicates (three in total). We defined a reproducible peak as one that received an IDR score ≤ 0.1 for at least two pairwise comparisons between biological replicates.

We identified 50,151 ATAC-seq peaks, 12,807 H3K4me1 peaks, 1,969 H3K4me3 peaks, and 3,744 H3K27me3 peaks (Supplemental Data S2). The number of ATAC-seq peaks we identified in the AEP assembly is similar to previously published *Hydra* ATAC-seq datasets generated using strain 105 animals (Siebert et al. 2019; Cazet et al. 2021). However, the number of peaks from our CUT&Tag libraries likely underrepresent the true number of genomic regions enriched for each respective histone modification. Thus, although we have demonstrated for the first time that CUT&Tag can successfully be applied to a cnidarian model, the protocol will require further optimization to improve sensitivity in the future. The establishment of CUT&Tag in *Hydra* offers substantial benefits over alternative chromatin mapping techniques, namely ChIP-seq, as CUT&Tag requires approximately two orders of magnitude fewer animals as input compared to equivalent *Hydra* ChIP-seq experiments (Reddy et al. 2020).

To analyze the CUT&Tag data collected from whole strain AEP *H. vulgaris* polyps, we first used Trimmomatic to remove stretches of low-quality base-calls and contaminating adapter sequence. We then aligned the data to the AEP assembly using Bowtie2. PCR duplicates were then identified and removed using Samtools. We then called peaks for the H3K4me1 H3K4me3 and H3K27me3 data with SEACR (Meers et al. 2019) using the IgG data as the background signal. To identify biologically reproducible peaks, we again performed IDR and selected peaks with an IDR score ≤ 0.1 for at least two of the three pairwise comparisons between biological replicates.

We used UROPA (Kondili et al. 2017) to annotate all ATAC-seq and CUT&Tag peaks based on the nearest TSS. We used deepTools (Ramírez et al. 2016) to generate the correlation heatmap globally comparing the aligned CUT&Tag and ATAC-seq data, to generate the data tracks used to depict read density along the AEP assembly, and to characterize the distribution of ATAC-seq and CUT&Tag data in and around genes. Individual plots visualizing the CUT&Tag, ATAC-seq, and sequence conservation data were generated using Gviz (Hahne and Ivanek 2016) and pyGenomeTracks (Lopez-Delisle et al. 2021).

#### *Hydra* single-cell atlas mapping and annotation

A step-by-step description of the single-cell RNA-seq atlas mapping and annotation methodology, including all relevant code, is provided in the markdown document *05_hydraAtlasReMap* available at github.com/cejuliano/brown_hydra_genomes.

We aligned the raw *Hydra* single-cell atlas sequencing data (previously deposited under BioProject PRJNA497966) to the AEP genome transcript models using the Drop-seq Tools alignment pipeline (github.com/broadinstitute/Drop-seq). Following mapping, we next determined which cell barcodes to include in downstream analyses. Because most beads in a Drop-seq experiment are not exposed to a lysed cell, only a small minority of sequenced cell barcodes are associated with a genuine single-cell transcriptome. Instead, most barcodes have low read counts attributable to contamination from ambient RNA. To differentiate between cell barcodes containing true single-cell transcriptomes and barcodes containing only transcriptomic noise, we generated plots that depicted the cumulative read fraction of cell barcodes ordered by read depth from highest to lowest. The curves generated by these plots have an elbow—an inflection point where the cumulative read fraction rapidly plateaus. This inflection point demarcates the transition from true biological signal to noise. For our downstream analyses, we used only read count data from the cell barcodes that preceded the elbow in the cumulative read plot.

Subsequent clustering and visualizations of the scRNA-seq data were done using Seurat (Hao et al. 2021). Prior to clustering, we performed additional filtering to remove cell barcodes with fewer than 300 or greater than 7,500 unique molecular identifiers (UMIs) as well as barcodes with fewer than 500 or greater than 75,000 total reads. We also removed any genes that were found in fewer than 3 cells. After filtering, we normalized the data using sctransform (Hafemeister and Satija 2019) and corrected for batch effects using reciprocal PCR as implemented in Seurat. We then clustered the single-cell transcriptomes using the Louvain algorithm (Waltman and Van Eck 2013) and visualized the results using a UMAP dimensional reduction (McInnes et al. 2018). We annotated the clustered dataset using a panel of previously validated marker genes (Figs. S6 and S7A) (Siebert et al. 2019).

As with prior analyses of the *Hydra* scRNA-seq atlas (Siebert et al. 2019), we found that many individual cell transcriptomes simultaneously contained multiple transcripts known to have mutually exclusive expression patterns. These chimeric transcriptomes are referred to as "doublets" and can result from either technical or biological causes (Siebert et al. 2019; Macosko et al. 2015). For example, battery cells, a prominent source of doublets in *Hydra* scRNA-seq data, are tentacle ectodermal cells in which both neurons and nematocytes are stably embedded (Bode and Flick 1976; Yu et al. 1985; Hufnagel et al. 1985). Because these three cell types are tightly physically associated in battery cell complexes, they are resistant to dissociation and are frequently sequenced as a single cell (Siebert et al. 2019).

To systematically identify likely doublets, we identified markers associated with ectodermal, endodermal, neuronal, nematocyte, gland, and germ cells using a Wilcoxon Rank Sum test as implemented in Seurat. We then calculated a holistic score representing how highly each cell in the atlas expressed each set of cell type markers using the Seurat AddModuleScore function (Fig. S7B). Because most doublets in *Hydra* include at least one epithelial cell (Siebert et al. 2019), we defined a doublet as a cell with a score greater than 0.2 for both an epithelial module and any other cell type module (Fig. S7C). For the sake of clarity and simplicity, we chose to exclude all doublet transcriptomes from the finalized version of the AEP genome-mapped atlas; however, we provide an alternative version of the atlas with doublets included (available at research.nhgri.nih.gov/HydraAEP/download/index.cgi?dl=fa), as certain cell types (e.g., battery cells) may require the inclusion of doublets to be properly represented. We repeated the batch correction, clustering, and UMAP dimensional reduction after removing all predicted doublets and found two remaining clusters, one that contained endodermal/interstitial doublets and another that appeared to contain cells expressing stress markers, that we removed prior to finalizing the set of cells included in the doublet-free version of the atlas. We again repeated the clustering and UMAP dimensional reduction steps to generate the final atlas presented in the main text, which we annotated using the same panel of previously validated marker genes described above.

To identify groups of co-expressed gene in the single-cell atlas, we performed non-negative matrix factorization (NMF) as implemented in the cNMF python package (Kotliar et al. 2019) on the full (doublets included) single-cell expression matrix. NMF is a dimensional reduction technique that, when applied to gene expression data, groups co-expressed genes into modules referred to as metagenes. The number of metagenes identified by NMF, a value referred to as k, needs to be specified prior to performing the factorization. The optimal value for k cannot be determined objectively and instead needs to be estimated empirically by evaluating a range of k values. Therefore, we performed an initial parameter sweep using k values ranging from 15 to 90 by steps of 5. The results from NMF depend on how the analysis is initialized, so we performed 200 independent runs for each k value that could then be combined to generate a consensus factorization result. We then selected a k value that maximized reproducibility across independent runs while simultaneously minimizing the differences between the factorized data and the original expression data. Based on these criteria, we selected a k value of 55. Our initial sweep of k values used steps of 5, so to more precisely identify the optimal k value we performed another parameter sweep for k values ranging from 50 to 60 by steps of 1. After evaluating the reproducibility and fidelity of the results from the fine resolution sweep, we selected a final k value of 56. We then generated the final consensus factorization results after first discarding individual runs that contained irreproducible results (see 05_hydraAtlasReMap.md for details).

**In situ hybridization**

To generate labeled RNA probes for performing in situ hybridization, we cloned and sequenced PCR products for the *Hydra* genes *G017021* (*scleraxis*) and *G008733* that had been amplified from oligo-dT-primed cDNA generated from whole adult male and female *H. vulgaris* polyps (Kiel AEP line). Amplicons were generated using the following PCR primers: *G017021*-forward: AGTTTAAAATGCTCCAATCTATAAGG; *G017021*-reverse: TAATACGACTCACTATA GGGTGATCTTAAAAATGTAACGCAAAATG; *G008733*-forward: GCTTTAGGCGGCTCAACAAA; *G008733*-reverse: ATTTAGGTGACACTATAGAACCTTTGTTTACGCCAGCA. The reverse primer sequences for *G017021* and *G008733* included T7 and SP6 promoter sequences respectively, allowing us to use purified PCR products as templates for in vitro transcription reactions using the Roche DIG RNA Labeling Kit (Roche Cat # 11175025910). The resulting DIG-labeled RNA products were then purified using the Zymogen RNA Clean & Concentrator-25 kit (Zymo Research Cat # R1017) and stored at -80˚C until use.

To perform in situ hybridization on whole *Hydra* polyps, we used a slightly modified version of a previously published protocol (Bode et al. 2009). For each in situ, 15 whole adult strain AEP *H. vulgaris* polyps that had been starved for two days were transferred to a 1.5 ml microcentrifuge tube, relaxed at room temperature (RT) for 1 minute in 1 ml *Hydra* medium (HM) containing 2% urethane, and then fixed in 1 ml HM containing 4% paraformaldehyde (PFA) at 4˚C overnight. All subsequent steps were performed at RT in 1 ml of solution with gentle rocking agitation unless otherwise indicated. Following overnight fixation, the fixative was removed with three quick washes in PBT (0.1% tween-20 in phosphate buffered saline, pH 7.4). The tissue was then bleached by transferring the samples gradually to 100% MeOH using 5-minute washes first in 33% MeOH in PBT then in 66% MeOH. The samples were then incubated in 100% MeOH for 1 hour. To maximize bleaching, the samples were incubated overnight in fresh 100% MeOH at -20˚C. The tissue was rehydrated using 1 wash with 66% MeOH, 1 wash with 33% MeOH in PBT, and three washes in PBT for 5 minutes each. The tissue was then permeabilized in 10 µg/ml proteinase K in PBT for 5 minutes. Proteinase activity was halted with a quick wash in 4 mg/ml glycine in PBT followed by a 10-minute wash in fresh glycine solution. The glycine solution and any residual proteinase K was then removed with three 5-minute washes in PBT. The samples were then washed twice in 0.1 M triethanolamine in PBT, once in 0.1 M triethanolamine in PBT containing 3 µl/ml acetic anhydride, once in 0.1 M triethanolamine in PBT containing 6 µl/ml acetic anhydride, then three times in PBT, all for 5 minutes each. Next, the tissue was refixed for 1 hour using 4% PFA in PBT. The fixative was removed with three 5-minute PBT washes followed by two 5-minute washes in 2X SSC (300 mM NaCl and 30 mM sodium citrate). In preparation for probe hybridization, the samples were incubated in 50% 2X SSC/50% hybridization solution (HS; 50% formamide, 5x SSC [750 mM NaCl and 75 mM sodium citrate], 1x Denhardt's solution, 100 µg/mL heparin, 0.1% Tween-20, and 0.1% Chaps) for 10 minutes, starting first at RT then gradually transitioning to hybridization temperature (56˚C). All subsequent pre-hybridization and hybridization steps were carried out at 56˚C. The tissue was incubated in HS for 10 minutes and then in HS containing 200 µg/ml yeast RNA for 2 hours. To prepare the DIG-labeled probes for hybridization, we added ~750 ng of probe to modified HS (50% formamide and 5x SSC) and denatured secondary RNA structures by incubating the solution at 85˚C for 5 minutes. The probe solution was then added to the sample tubes after first being diluted in fresh HS containing 200 µg/ml yeast RNA to a final probe concentration of ~3 ng/ul. The samples were then left to hybridize for ~60 hours with no agitation.

Excess probe was removed using a sequence of single, 5-minute washes in HS, 75% HS/25% 2X SSC, 50% HS/50% 2X SSC, and then 25% HS/75% 2X SSC at 56˚C. The samples were then washed twice with 2X SSC containing 0.1% CHAPS for 30 minutes each, with the first wash occurring at 56˚ C and the second at 37˚ C. Unbound probe was digested by treating the tissue with 20 µg/ml RNase A in 2X SSC containing 0.1% CHAPS for 30 minutes at 37˚C without agitation. RNase A was then removed using two 10-minute washes at 37˚C and two 30-minutes washes at 55˚C in 2X SSC containing 0.1% CHAPS. The samples were then transitioned back to RT and washed three times with MABT (100 mM maleic acid, 150 mM NaCl, 0.1% Tween 20, pH 7.5) for five minutes each. Non-specific protein interactions in the tissue were then blocked with a two-hour incubation in blocking solution (MABT containing 1% BSA and 20% sheep serum) at 4˚C. The samples were then resuspended in a 1:2000 dilution of Anti-Digoxigenin-AP (Roche Cat # 11093274910) in blocking solution and incubated overnight at 4˚C without agitation.

Following antibody binding, the samples were transitioned back to RT and excess antibodies were removed with eight 20-minute washes in MABT. The tissue was then washed once in NTMT (100 mM NaCl, 100 mM Tris, 50 mM MgCl2, 0.1% Tween-20, pH 9.5) for 5 minutes. During this NTMT wash, the samples were transitioned to six-well plates. The NTMT was then replaced with 20 µl/ml NBT/BCIP solution (Roche Cat # 11681451001) in NTMT. The staining reaction proceeded for an empirically determined time (~1-2 hours) and was subsequently stopped using three quick PBT washes. To reduce non-specific signal in the tissue, the tissue was transitioned into 100% EtOH using 5-minute washes first in 33% EtOH in PBT then in 66% EtOH. The tissue was then incubated in 100% EtOH until the staining in the tissue turned from purple to blue (~30 minutes). The tissue was then rehydrated using single 5-minute washes in 66% EtOH then 33% EtOH in PBT. Finally, residual EtOH was removed using three quick PBT washes. The in situs were documented using a Leica DM5000B microscope (camera Leica DFC310FX), a Leica M165C digital stereo microscope (camera MC170HD), or a Zeiss Axiophot microscope (camera Leica DFC 550).

**Characterization of gene age in the *Hydra* single-cell atlas**

A step-by-step description for our methodology for characterizing the cell-type-specific transcriptional patterns associated with gene age, including all relevant code, is provided in the markdown document *06_geneAge* available at github.com/cejuliano/brown_hydra_genomes.

To estimate the age for each *Hydra* gene model, we adopted a phylostratigraphic approach (Domazet-Lošo et al. 2007). We used the orthology predictions generated from our OrthoFinder analysis (see "AEP genome gene annotation") to identify the most recent clade that contained all orthologs of each *Hydra* gene (i.e. the "clade of origin"). We defined gene age to be the age of each gene's clade of origin. For example, if a gene in *Hydra* had orthologs throughout cnidaria, but lacked any orthologs outside of cnidaria, then Cnidaria would be considered that gene's clade of origin. Therefore, the gene likely first emerged after the split of Bilateria and Cnidaria but before the split of Anthozoa and Medusozoa.

We next used these gene age predictions to characterize the relationship between gene age and cell-type specific transcription in our *Hydra* single-cell atlas. To do this, we first generated lists of genes that were present in the transcriptomes of each cell type in our atlas by identifying all genes with an average expression level above 0.05 normalized counts per cell for each cell type. Then, to exclude ubiquitously expressed genes that do not vary across different cell types, we used the Seurat FindVariableFeatures function to identify 7,500 genes with high or intermediate levels of variability across the *Hydra* atlas and excluded genes from our cell type transcriptomic profiles if they were not found in this variable gene list. To calculate the relative enrichment of each age across *Hydra* cell types, we calculated the odds that a gene expressed in a certain cell type will be of a certain age. We found that the transcriptomes of all cell types were heavily skewed towards ancient genes that predate Metazoa, likely reflecting the essential and deeply conserved functions of ancient genes. However, cell-type-specific enrichment patterns did emerge when we normalized the enrichment profiles across cell types. We calculated single-cell transcriptomic age index values by applying a previously described formula (Domazet-Lošo and Tautz 2010) to the normalized *Hydra* atlas single-cell gene expression matrix.

**Genomic sequence conservation analysis**

A step-by-step description of the single-cell RNA-seq atlas mapping and annotation methodology, including all relevant code, is provided in the markdown document *07_genomeConservation* available at github.com/cejuliano/brown_hydra_genomes.

We generated a cross-species whole-genome alignment of the *C. hemisphaerica*, *H. viridissima*, *H. oligactis*, strain 105 *H. vulgaris*, and strain AEP *H. vulgaris* genome assemblies using Progressive Cactus (Armstrong et al. 2020). To facilitate the alignment, repetitive regions in each genome were soft-masked prior to running Progressive Cactus, causing them to be largely excluded from the resulting alignment. To quantify sequence conservation rates in across the AEP assembly using the resulting alignment, we used a custom Python script to count the number of non-AEP genomes with the same nucleotide for every position of the AEP assembly that was included in the whole-genome alignment. For visualizing the sequence conservation results (as in Fig 2B), we smoothed the per-base conservation results using a 100 bp moving window. We used deepTools (Ramírez et al. 2016) to characterize the distribution of conservation rates around the AEP assembly gene models.

To identify putative conserved transcription factor binding sites (TFBS) in the AEP assembly, we first used FIMO (Grant et al. 2011) to identify putative binding sites in all four *Hydra* genomes in our alignment using a custom database of non-redundant vertebrate, insect, and nematode binding motif sequences from the JASPAR database (Fornes et al. 2020). To generate a control dataset, we also performed TFBS prediction using a version of our custom motif sequence data base where the nucleotide order of each motif had been shuffled. We then used the Hierarchical Alignment API (Hickey et al. 2013) in conjunction with our cross-species genome alignment to convert the coordinates of all non-AEP TFBS coordinates to their equivalent coordinates in the AEP assembly. This allowed us to determine if a given TFBS in the AEP assembly was also present in other *Hydra* genomes. We considered a TFBS in the AEP assembly to be conserved if it was present in the strain 105 *H. vulgaris* assembly and at least one other *Hydra* genome. To further filter our conserved TFBS list to sites that were most likely to be functionally relevant, we eliminated any predicted binding sites that did not fall within an ATAC-seq peak or that overlapped protein coding sequence. To identify motif sequences from our custom database that showed evidence of conservation in *Hydra*, we used a chi-square test, as implemented in R, to identify motifs with significantly (FDR ≤ 0.01) higher conservation rates than shuffled controls.

To identify putatively conserved CREs, we used deepTools (Ramírez et al. 2016) to calculate the average level of sequence conservation for each ATAC-seq and CUT&Tag peak in the AEP assembly. We calculated these sequence conservation rates using pairwise comparisons between the AEP assembly and each non-AEP assembly in our whole-genome alignment, such that each peak received four separate conservation scores (e.g., one score for the AEP-105 alignment, one score for the AEP-*oligactis* alignment, etc.). We then used k-means clustering, as implemented in R, to partition peaks into two populations—a high-scoring population and a low-scoring population—for each pairwise species comparison. We defined a peak as conserved if it was classified as high scoring in at least two pairwise comparisons. To characterize the distribution of conserved enhancer-like CREs around genes in the AEP-assembly (presented in Fig 2 C,D), we used UROPA (Kondili et al. 2017) to calculate the distance from each H3K4me1 and ATAC-seq peak to the nearest TSS. To remove possible core promoter peaks from this analysis, we disregarded all H3K4me1 and ATAC-seq peaks that overlapped a H3K4me3 peak prior to visualizing the TSS distance distribution.

#### Prediction of transcriptional regulators in *Hydra*

A step-by-step description of the *Hydra* transcriptional regulator analysis, including all relevant code, is provided in the markdown document *10_hydraRegulators* available at github.com/cejuliano/brown_hydra_genomes.

To identify motifs enriched in the putative regulatory regions of genes belonging to cell-type-specific gene co-expression programs in the *Hydra* single-cell atlas, we used gene set enrichment analysis (GSEA) as implemented in the fgsea R package (Korotkevich et al. 2021; Subramanian et al. 2005). GSEA requires two inputs: 1) a set binary of classifications that groups together genes associated with a feature or process of interest (i.e., a gene set), and 2) a set of continuous scores that can be used to rank genes. To test for enrichment, GSEA evaluates if the members of a given gene set show a non-random distribution in their score rankings (i.e., if the gene set is biased towards having higher or lower scores). If a gene set has a non-random distribution, it indicates that the feature or process that was used to group those genes (e.g., the presence of a specific motif in nearby regulatory regions) is associated with the metric used to generate the gene rankings (e.g., a gene co-expression score for a specific cell type). The strength of this association is quantified using a metric called the normalized enrichment score, with higher scores indicating a stronger bias for the gene set to be associated with high gene ranks.

To perform a motif enrichment analysis using GSEA, we used our conserved TFBS predictions (described above in "Genomic sequence conservation analysis") to generate gene sets that grouped genes according to to the conserved binding motifs that were present in their putative regulatory regions, such that each motif was assigned a list of genes that were predicted to be regulated by the motif's cognate TF. For the continuous scores used to order genes in the GSEA, we used the *Hydra* atlas NMF gene scores (NMF described in "Single-cell atlas mapping and annotation"), which reflect how strongly the expression pattern of a gene matched the expression pattern associated with a given metagene. After performing GSEA for each metagene in the *Hydra* atlas, we discarded any enrichment scores that were not significant (adjusted P-value > 0.01) to reduce noise in the enrichment results. We then mapped these enrichment scores onto the *Hydra* atlas by generating single-cell enrichment scores for each motif. To do this, we used NMF cell scores, which reflect how well each metagene reflected a cell's overall transcriptomic profile, to calculate a weighted average enrichment score for each cell, with enrichment scores from highly scoring metagenes contributing more strongly than lowly scoring metagenes.

To identify the candidate transcription factors that could plausibly bind the motifs associated with each metagene, we first used metadata available through the JASPAR and UniProt databases to identify the Pfam DNA binding domains present in each motif's cognate TF.

We then generated a list of candidate regulators for each motif by identifying the AEP gene models that possessed the appropriate DNA binding motifs. To determine the most likely candidate regulators for each motif, we used the single-cell atlas to identify TFs whose expression was correlated with the enrichment pattern of their cognate motif.

A common problem that arises when performing correlation analyses using single cell RNA-seq data is the high frequency of 'dropouts', instances where low and moderately expressed genes are completely missed in a random subset of cell transcriptomes due to low sequencing depth. To mitigate this source of noise, and thus facilitate the comparison of motif enrichment and TF expression patterns, we used the *Hydra* atlas NMF results to generate an imputed version of the single-cell expression data. The results of a single-cell RNA-seq NMF analysis are two matrices, a gene score matrix and a cell score matrix, that approximate the original expression matrix when multiplied together. Importantly, this NMF-derived approximation eliminates the cell-to-cell heterogeneity caused by dropouts, thus facilitating single-cell expression correlation analyses.

Using the imputed read count matrix, we performed a correlation analysis to identify motifs whose enrichment pattern was correlated with the expression pattern of a TF that possessed the appropriate DNA binding domain. TFs with a motif enrichment correlation score ≥ 0.5 were deemed candidate regulators. We also reviewed possible regulator/motif pairs manually, allowing us to catch marginal cases where TFs were expressed in only a subset of cells where the target motif was enriched, causing them to fall slightly below our correlation score threshold (e.g., *zic1* and *zic4*).

To control for the possible contribution of sequence bias to our enrichment results, we repeated our GSEA and TF expression correlation analysis using shuffled versions of each motif (see 10_hydraRegulators.md for details). We found that while some shuffled motifs were significantly enriched in the *Hydra* atlas, the enrichment patterns of the shuffled motifs were overwhelmingly different from the enrichment patterns of their unshuffled counterparts. Specifically, the enrichment patterns of over 90% (832/907) of shuffled motifs had a correlation score ≤ 0 when compared to the enrichment patterns of the unshuffled motifs. This demonstrates that the enrichment patterns we observed using the unshuffled motifs were not driven primarily by sequence composition biases. We also found that the correlation scores between motifs and their candidate regulators were significantly higher when using unshuffled motifs when compared to shuffled motifs (student's t-test P-value ≤ 2.2e-16), suggesting the enrichment patterns for the unshuffled motifs better reflected the regulatory activity of *Hydra* TFs.

**Re-aligning the *Clytia* single-cell atlas**

A step-by-step description of the approach for generating new *Clytia* gene models and the subsequent re-alignment and clustering of the *Clytia* single-cell atlas, including all relevant code, is provided in the markdown document *11_clytiaAtlasReMap* available at github.com/cejuliano/brown_hydra_genomes.

The initial published version of the *Clytia* single cell RNA-seq atlas used a newly generated set of gene models for the original version of the *Clytia* genome as a reference for read mapping (Leclère et al. 2019; Chari et al. 2021). However, we used an updated version of the *Clytia* genome (available at metazoa.ensembl.org/Clytia_hemisphaerica_gca902728285) for our cross-species whole genome alignment. To maintain a consistent genome reference across analyses, and to maximize the completeness of the gene models used for mapping the single cell data, we generated a custom set of gene predictions for the updated version of the *Clytia* genome. To do this, we first generated a preliminary set of gene predictions by aligning both the new transcriptome generated in the *Clytia* single-cell atlas publication and the transcript models from the original *Clytia* genome publication to the updated Clytia genome using PASA. We then combined the PASA gene models with the gene models for the updated genome assembly using AGAT (github.com/NBISweden/AGAT). The resulting gene models were more complete than the pre-existing gene models for the updated genome assembly, as indicated by the increased number of complete single copy orthologs identified using BUSCO (Table S1). We then aligned the raw *Clytia* single-cell data to the newly generated transcript models using the 10X Cell Ranger pipeline. Following mapping, we selected the cell barcodes used for downstream analysis by retaining only those cells that were present in the original published version of the *Clytia* atlas.

We then clustered the re-mapped data using the Louvain algorithm as implemented in Seurat and found that our analysis recapitulated the cell type clustering results from the original publication (see 11_clytiaAtlasReMap.md for details), validating our mapping and clustering approach.

To characterize the cell-type-specificity of *Clytia* genes that were lost in the *Hydra* lineage, we first used the results from our OrthoFinder analysis (described above in "AEP genome gene annotation") to identify *Clytia* genes with orthologs in *Hydractinia echinata* (the other non-*Hydra* hydrozoan in our analysis) but with no orthologs in any of the *Hydra* proteomes in our analysis. We then generated a holistic score representing how strongly each cell in the *Clytia* atlas expressed these lost genes using the Seurat AddModuleScore function.

**Aligning the *Clytia* and *Hydra* single-cell atlases**

A step-by-step description of the *Clytia* and *Hydra* single-cell RNA-seq atlas alignment, including all relevant code, is provided in the markdown document *12_crossSpeciesAtlasAlignment* available at github.com/cejuliano/brown_hydra_genomes.

To align the *Clytia* and *Hydra* single cell atlases, we first identified all *Hydra* genes with unambiguous one-to-one orthologs in *Clytia* using the results from our OrthoFinder analysis (described above in "AEP genome gene annotation"). We then subset the *Clytia* and *Hydra* single-cell read count matrices to only include these one-to-one orthologs and converted all *Clytia* gene names to their *Hydra* equivalent. After the data was reformatted, we used reciprocal principal component analysis as implemented in Seurat to combine and align the *Hydra* and *Clytia* single-cell RNA-seq data. We then performed Louvain clustering on the aligned data and visualized the results using a UMAP dimensional reduction. We annotated the resulting clusters by propagating the cell type annotations associated with each cell barcode from the uncombined versions of the *Clytia* and *Hydra* atlases.

To quantify the transcriptional similarities between *Clytia* and *Hydra* cell types, we made use of a previously described alignment metric (Tarashansky et al. 2021). To calculate this alignment score, we performed a mutual nearest neighbor analysis (MNN) as implemented in the BiocNeighbors R package. This analysis identified all cross-species cell pairs where each member of the pair was among the other's 30 nearest cross-species neighbors in principal component space. We calculated the alignment score by determining the portion of total MNNs for a cell type of interest that belonged to each cell type in the other species. We retained all cross-species cell type pairs with an alignment score ≥ 0.05. We also calculated a single-cell divergence score, which measures the average distance between a cell and it's thirty nearest cross-species neighbors in principal component space. A smaller divergence score thus indicates that the transcriptomic profile of a given cell is more like the transcriptomic profiles of cells from the other species than cells with higher divergence scores.

To identify genes with conserved expression patterns in *Clytia* and *Hydra*, we first performed a high-resolution Louvain clustering analysis to generate 'pseudo-cells' that grouped together small sets of *Clytia* and *Hydra* cells with similar gene expression profiles. We then calculated average gene expression values for each species in each pseudo-cell. We designated a gene as having a conserved expression pattern if the pseudo-cell expression values in the two species had a correlation score > 0.65.

**Predicting conserved transcriptional regulators in *Clytia* and *Hydra***

A step-by-step description of the *Clytia* transcriptional regulator analysis and the comparison of candidate regulator predictions in *Hydra* and *Clytia*, including all relevant code, is provided in the markdown document *13_conservedRegulators* available at github.com/cejuliano/brown_hydra_genomes.

To identify cell-type-specific gene co-expression modules in *Clytia*, we performed NMF on the raw *Clytia* atlas single-cell expression matrix, following the same steps as described above for the *Hydra* single-cell atlas (see "Single-cell atlas mapping and annotation"). To identify the optimal number of metagenes, we first performed a broad sweep of k values from 15 to 90 by steps of 5. We observed a local maximum in the stability of the NMF results for k=40, prompting us to perform a second sweep of k values from 35 to 45 by steps of 1. Based on this fine resolution sweep, we chose a k value of 37. We then generated the final consensus factorization results after first discarding individual runs that contained irreproducible results.

Because cis-regulatory element annotations were not available for *Clytia*, we were unable to use the same motif enrichment approach as for our analysis in *Hydra*. Instead, to isolate presumptive promoter sequences we extracted all sequences that fell within 1 kb upstream of a TSS. Then, for each *Clytia* metagene, we generated a ranked list of these putative promoters with sequences that were near genes strongly associated with the metagene placed at the top of the list and sequences near genes that were weakly associated placed at the bottom of the list. We then used these ranked promoters as input for an Analysis of Motif Enrichment (AME) (McLeay and Bailey 2010). To map the AME results onto the *Clytia* single-cell atlas, we calculated single-cell weighted averages of the significant (E-value < 10) fold-enrichment results for each metagene using the NMF metagene cell scores.

To identify conserved regulators in *Hydra* and *Clytia*, we manually reviewed the expression patterns and associated motif enrichment patterns for all TFs that both had a conserved expression pattern in *Clytia* (see Aligning the *Clytia* and *Hydra* single-cell atlases") and were designated as candidate regulators in *Hydra* (see "Prediction of transcriptional regulators in *Hydra*"). We considered a TF to be a conserved regulator when both the expression of the TF and the enrichment of its cognate motif were localized to the same cell populations in *Clytia* and *Hydra* in the cross-species single-cell atlas.

To determine if the degree of overlap in motif enrichment patterns for the *Hydra* and *Clytia* atlases was greater than would be expected by chance, we repeated our analysis using shuffled versions of each motif. We then quantified the degree of overlap in motif enrichment patterns using the same pseudo-cell correlation approach described above (see "Aligning the *Clytia* and *Hydra* single-cell atlases"). We observed no highly correlated ($r \geq 0.5$) enrichment patterns when using shuffled motifs, whereas we found 13 highly correlated enrichment patterns when using unshuffled motifs (Supplemental Data S15). This suggests that the enrichment overlap we observed using unshuffled motifs are likely indicative of conserved TF function and are not driven purely by chance.

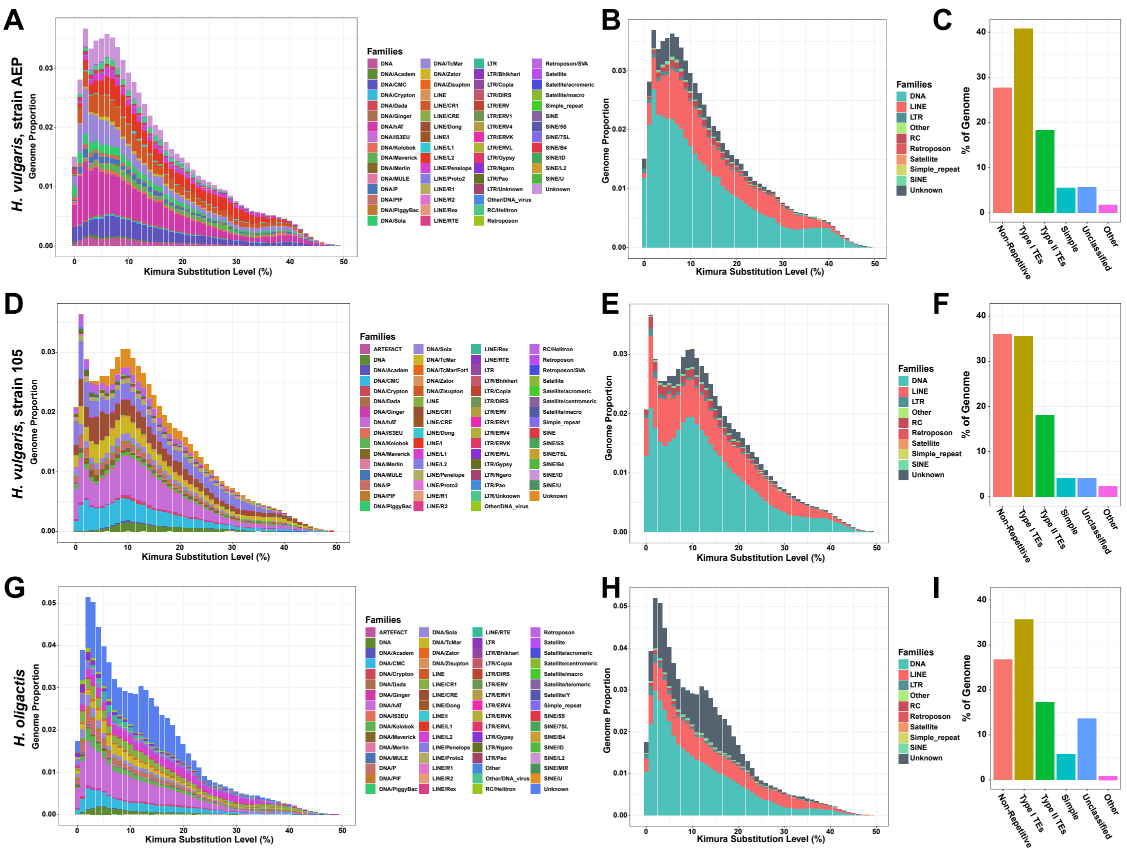


**Fig. S1.** Repeat composition of *Hydra* genomes. Summary plots of repeat composition in the (*A*-*C*) strain AEP *H. vulgaris*, (*D*-*F*) strain 105 *H. vulgaris*, and (*G*-*I*) *H. oligactis* genomes. Repeat landscapes are presented at the level of repeat subfamilies (*A*, *D*, and *G*) and broader repeat classes (*B*, *E*, and *H*). (*C*, *F*, and *I*) Total proportions for repetitive and non-repetitive elements across each genome.

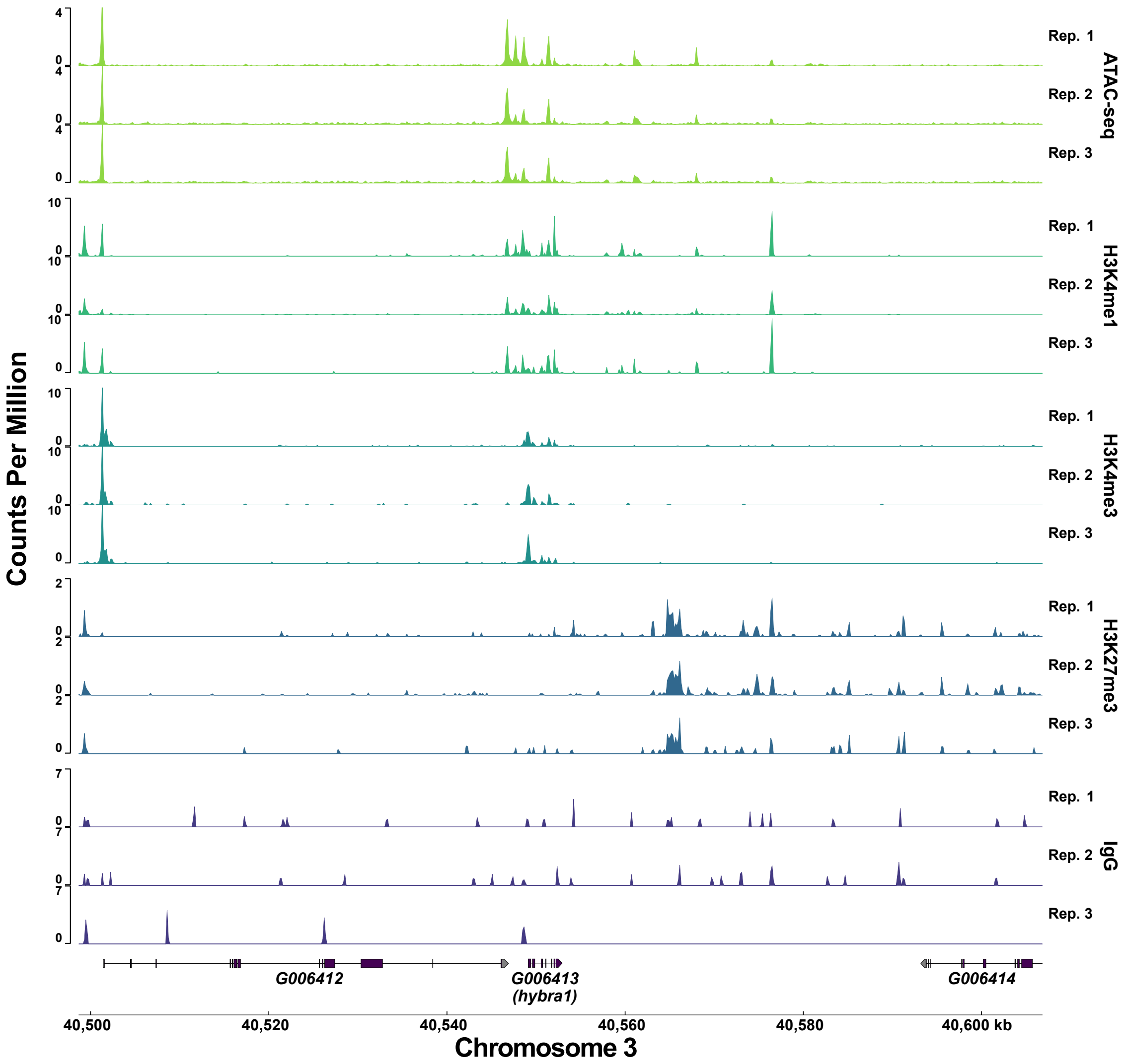


**Fig. S2.** Representative plot of all CUT&Tag and ATAC-seq biological replicates centered on the *brachyury1* (*bra1*) gene.

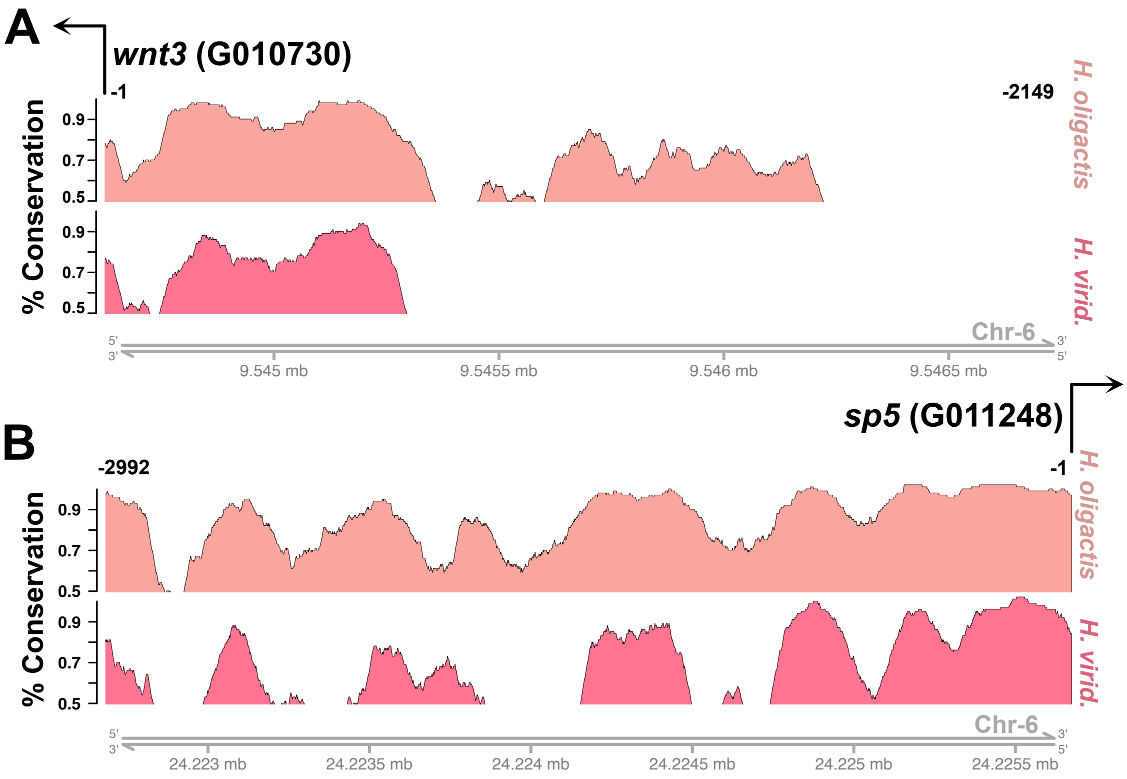


**Fig. S3.** Cross-species whole-genome alignments reveal conserved non-coding sequences in the strain AEP *H. vulgaris* genome. (*A* and *B*) 100 Bp moving window sequence conservation in sequence upstream of (*A*) *wnt3* and (*B*) *sp5* recapitulates previous results that used manual alignments (Vogg et al. 2019).

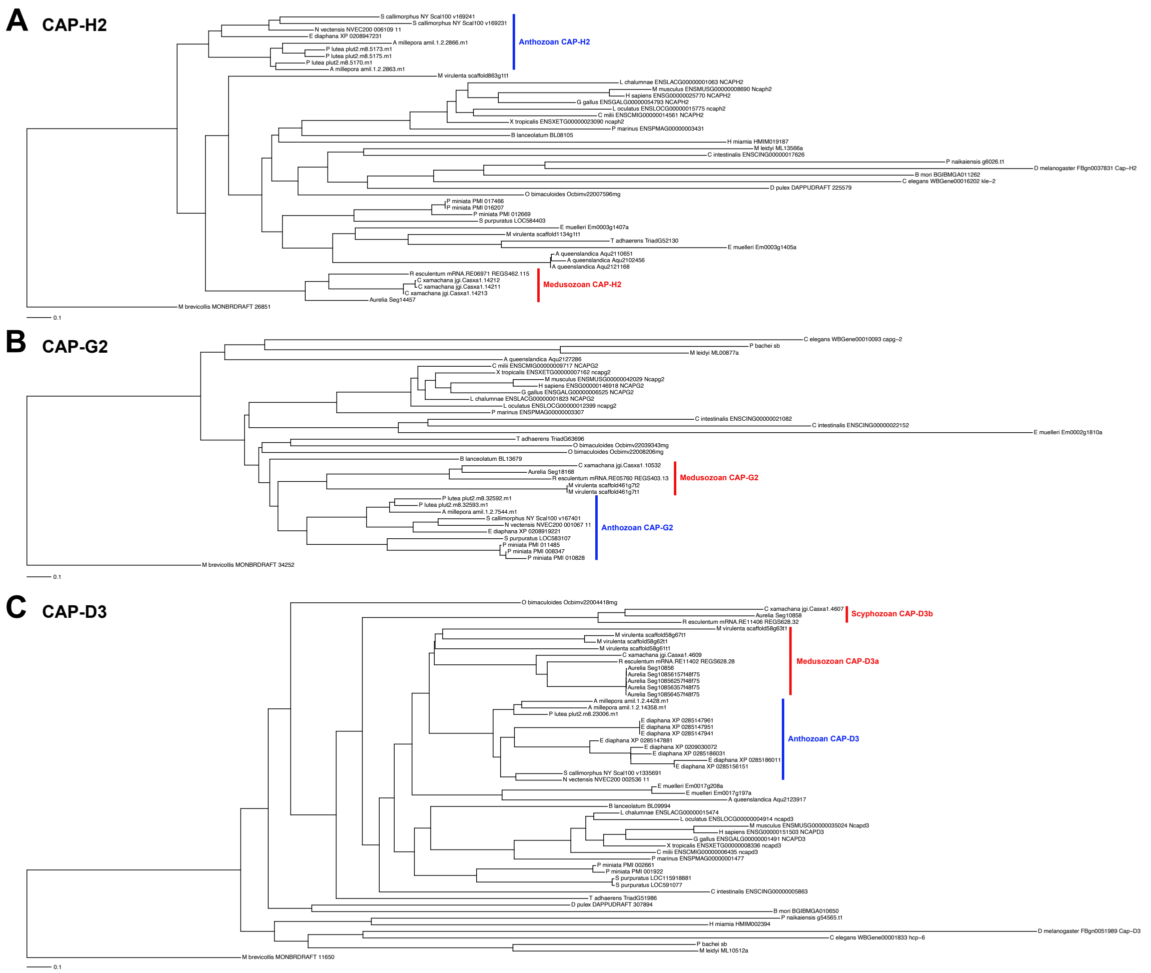


**Fig. S4.** Condensin II subunits are absent in hydrozoans. Phylogenies constructed by Orthofinder identify orthologs of the condensin II subunits (*A*) CAP-H2, (*B*) CAP-G2, and (*C*) CAP-D3 in anthozoans and non-hydrozoan medusozoans (Acraspeda), but not in hydrozoans.

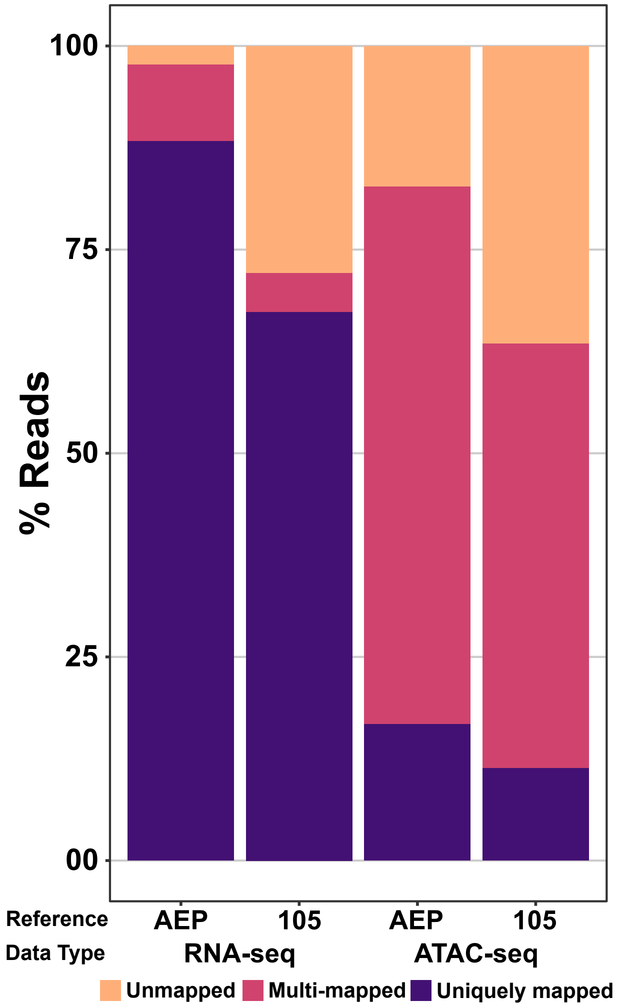


**Fig. S5.** Mapping efficiency of strain AEP *H. vulgaris* ATAC-seq and RNA-seq data are reduced when aligned to the strain 105 *H. vulgaris* genome reference.

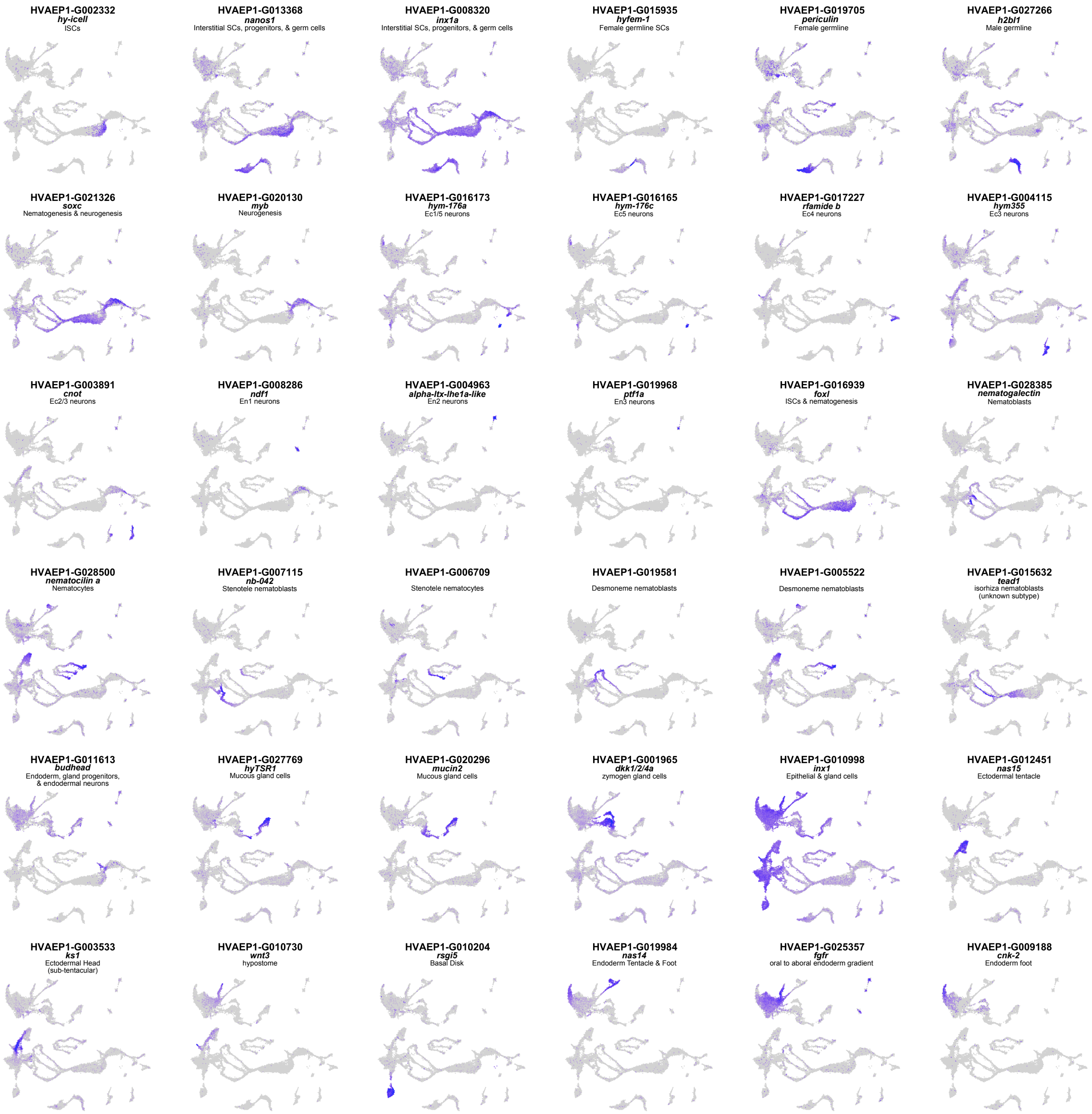


**Fig. S6.** Cluster annotation for the version of the strain AEP *H. vulgaris* single cell atlas that includes doublets using marker gene expression. All markers presented were validated in the initial atlas publication (Siebert et al. 2019).

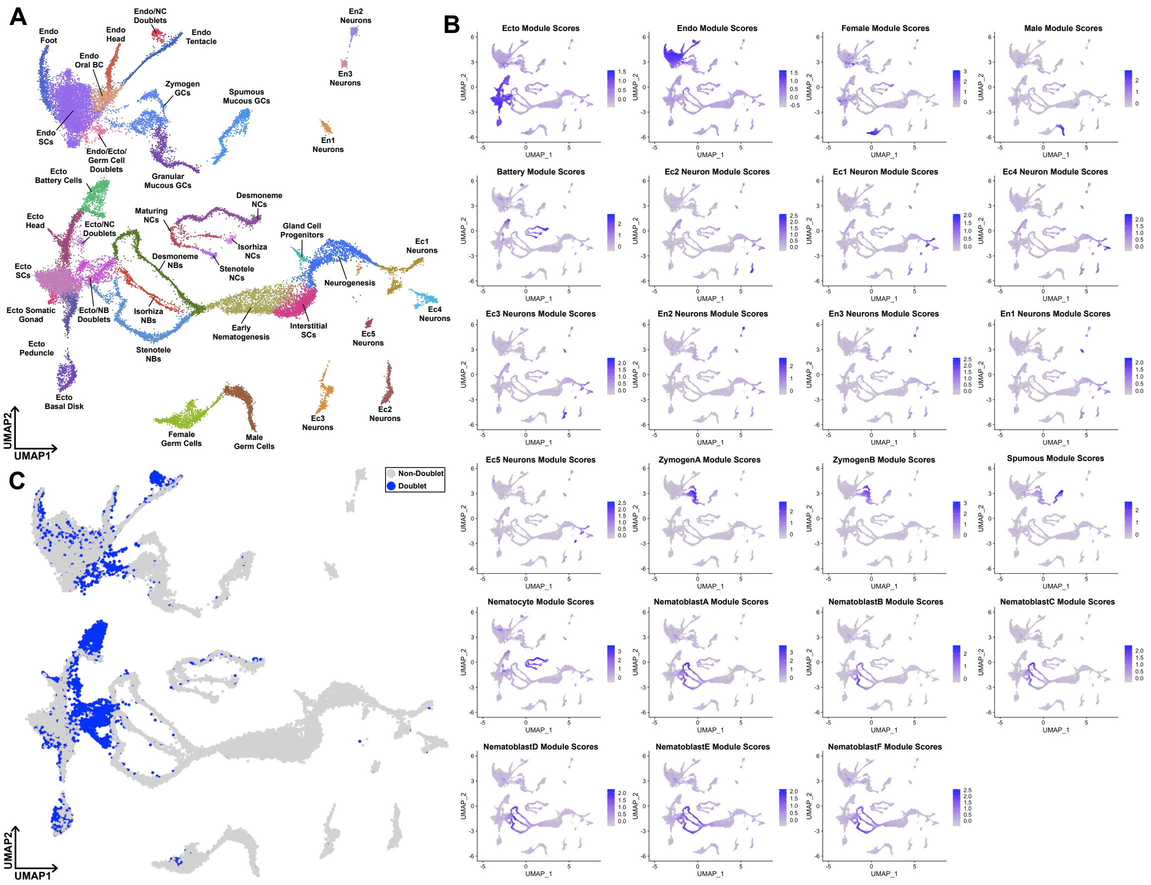


**Fig. S7.** Identification of doublets in the *Hydra* single-cell RNA-seq atlas. (*A*) Uniform Manifold Approximation and Projection (UMAP) dimensional reduction of the *Hydra* single-cell RNA-seq atlas mapped to the AEP reference genome prior to doublet removal. NCs, nematocytes; NBs, nematoblasts; SCs, stem cells; Ecto, ectodermal epithelial cells; Endo, endodermal epithelial cells; GCs, gland cells; Ec, neuron subtypes found in the ectoderm; En, neuron subtypes found in the endoderm. (*B*) Module scores for cell-type-specific gene expression programs as calculated by the Seurat AddModuleScore function. Cell-type-specific genes were identified using a Wilcoxon Rank Sum test as implemented in Seurat. (*C*) UMAP plot highlighting all cells identified as doublets. Doublets were defined as having a module score greater than 0.2 for both an epithelial module and any other cell type module.

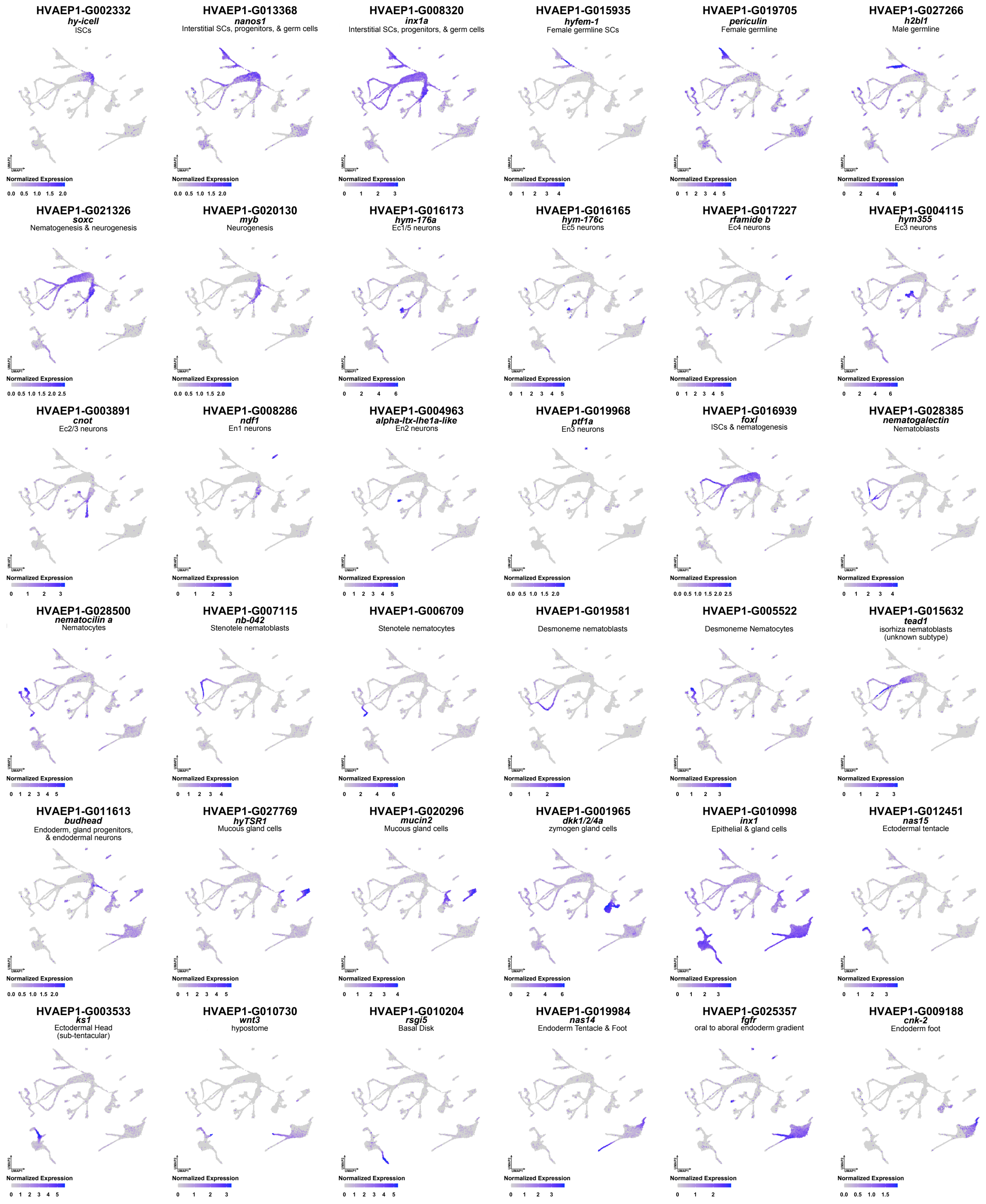


**Fig. S8.** Cluster annotation for the strain AEP *H. vulgaris* single cell atlas using marker gene expression. All markers presented were validated in the initial atlas publication (Siebert et al. 2019).

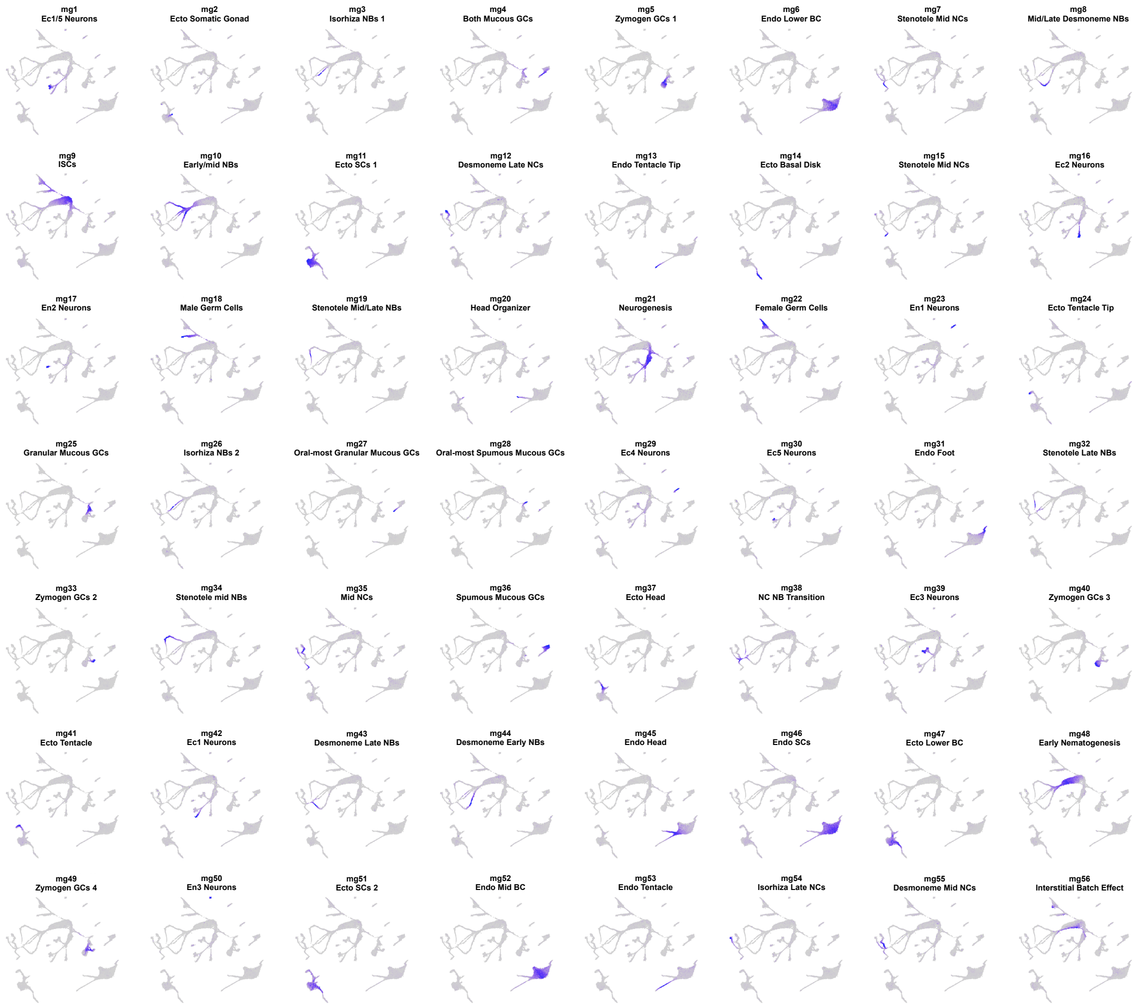


**Fig. S9.** Non-negative matrix factorization (NMF) identifies cell-type-specific co-expressed gene modules in the strain AEP *H. vulgaris* atlas. UMAP plots colored to highlight the cells expressing the 56 modules of co-expressed genes (i.e., metagenes) identified using NMF. More intense purple coloration indicates higher overall expression of a given metagene.

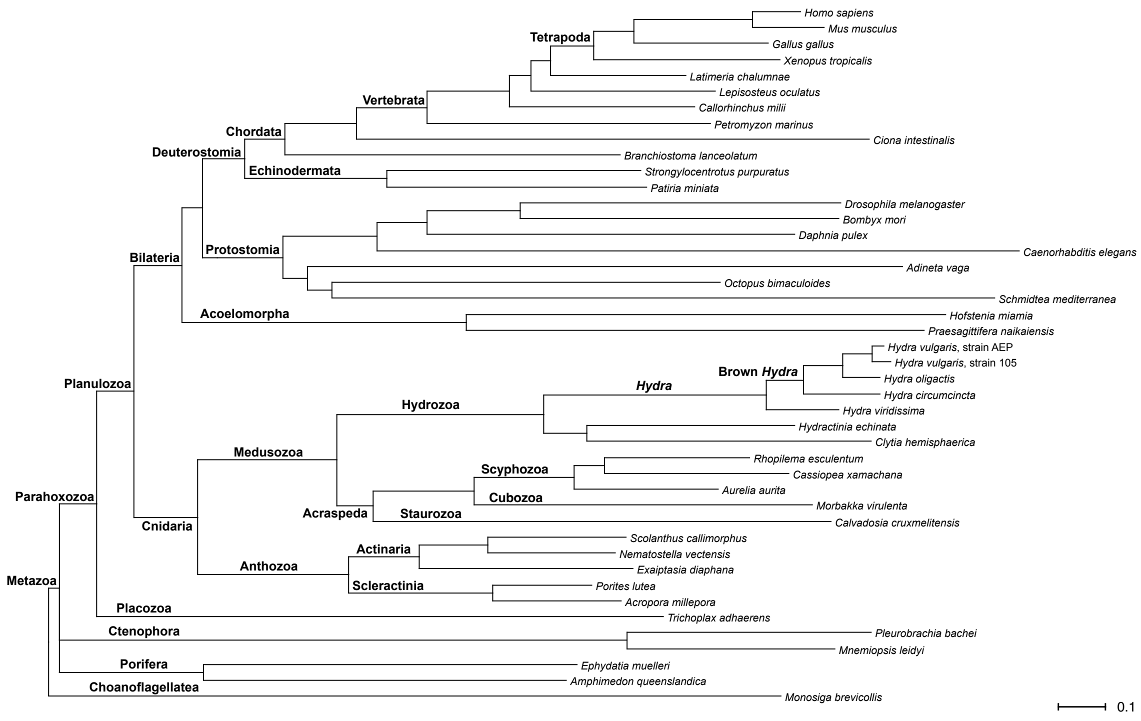


**Fig. S10.** Phylogeny of proteomes used in Orthofinder analysis. Proteome sources are provided in Table S2.

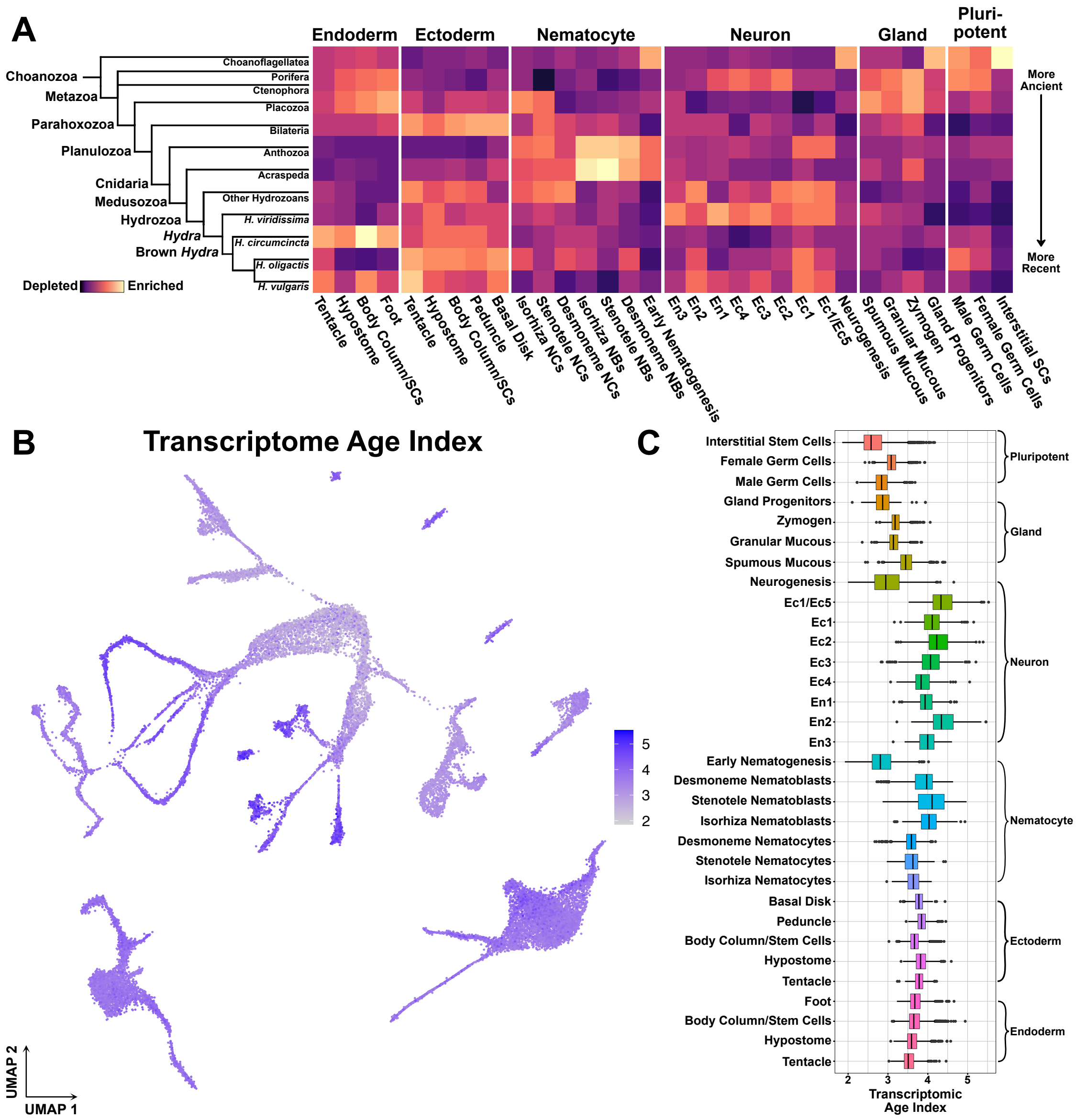


**Fig. S11.** Characterizing the relationship between gene age and cell-type-specific expression. (*A*) Heatmap depicting the relative enrichment of gene families by evolutionary age in the transcriptomes of different cell types suggest distinct evolutionary timelines. (*B-C*) Holistic quantification of single-cell transcriptome ages. The transcriptomic age index (TAI) is a weighted average that combines transcript abundance with gene age. High values of the resulting metric indicate a transcriptome is made up of relatively more recent genes and low values indicate a transcriptome is made up of relatively more ancient genes. (*B*) UMAP plot depicting TAI values for all single-cell transcriptomes in the *Hydra* cell atlas. (*C*) Boxplot of TAI values averaged by cell type.

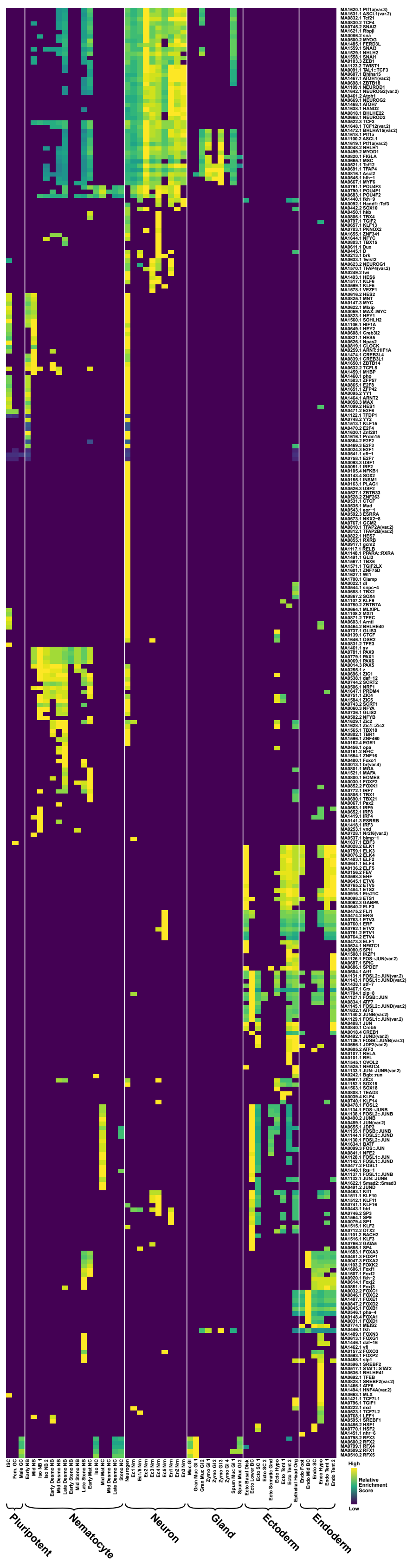
High
Relative Enrichment Score
Low
Pluripotent
Nematocyte
Neuron
Gland
Ectoderm
Endoderm

**Fig. S12.** Full motif enrichment results for the *Hydra* cell atlas that includes redundant motifs. Enrichment scores that were not significant (adjusted P-value > 0.01) were set to zero. Heatmap values are normalized by row (i.e. by motif). Motifs are referred to using both their unique JASPAR ID (formatted as MA####.#) and the abbreviated name of their corresponding TF.

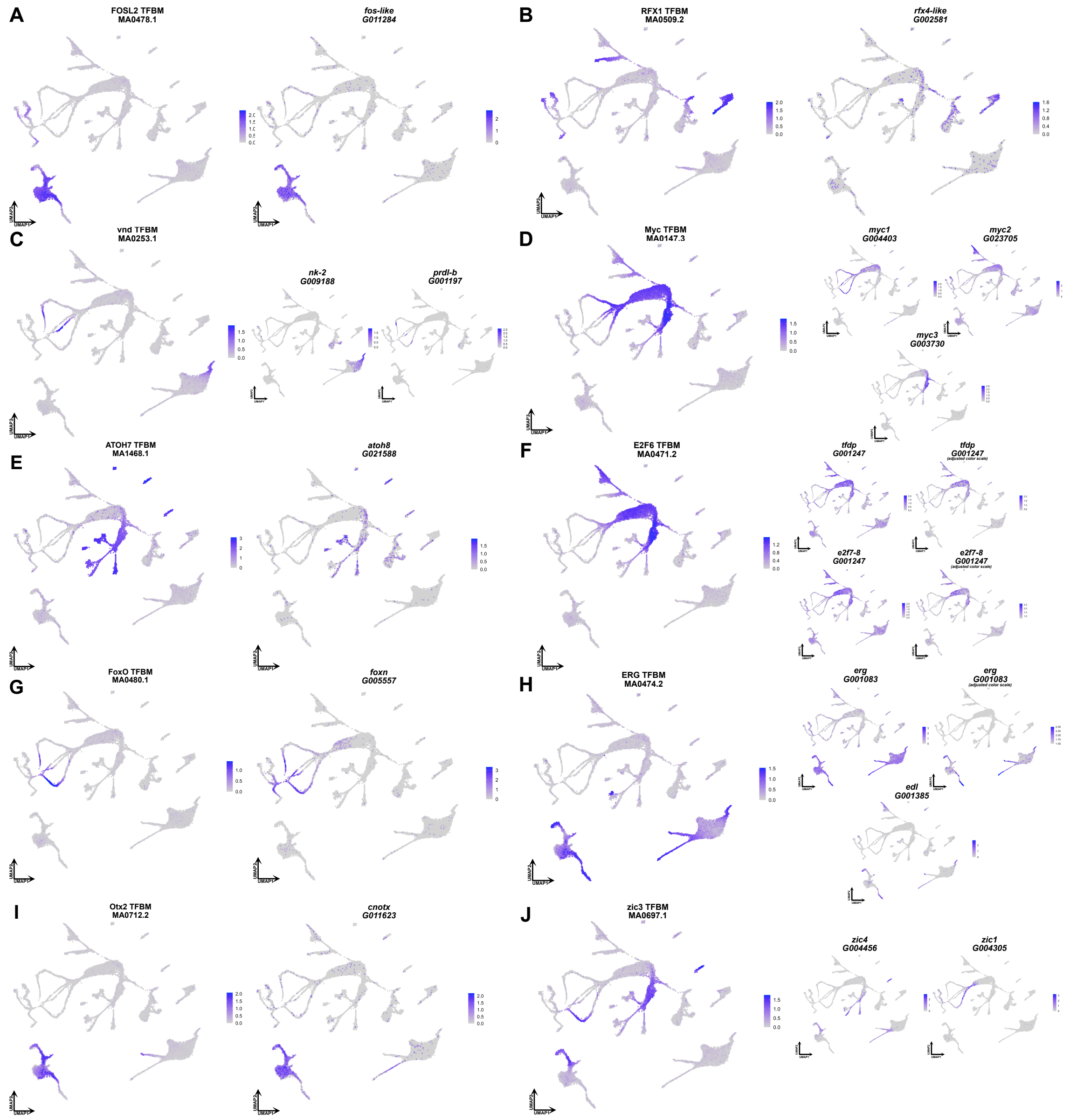


**Fig. S13.** Additional candidate regulators of gene co-expression in *Hydra*. Motif enrichment and gene expression correlation suggest that (*A*) *fos-like* is a regulator in ectodermal head and body column cells; (*B*) *rfx4-like* is a regulator in mucous gland cells; (*C*) the homeobox TFs *nk-2* and prdl-b are regulators in endodermal foot cells and nematoblasts respectively; (*D*) myc family transcription factors (TFs) are regulators in interstitial stem cells and progenitors; (*E*) *atoh8* is a regulator in mature and differentiating neurons; (*F*) e2f family TFs are regulators in interstitial stem cells, progenitors, and germ cells; (*G*) *foxN* is a regulator in late nematoblasts; (*H*) ets family TFs are regulators in epithelial cells at the extremities (i.e., tentacle and foot tissue); (*I*) *cnotx* is a regulator in ectodermal cells in the body column, head, and tentacles; and (*J*) zic family TFs are regulators in ectodermal tentacle cells, Ec4 neurons, and desmoneme nematoblasts. Note that for some gene expression plots (*tfdb*, *e2f7-8*, and *erg*) two plots with different color scales are presented to highlight cells with high expression levels. Color scales for motif plots refer to enrichment scores and normalized read counts in the gene expression plots.

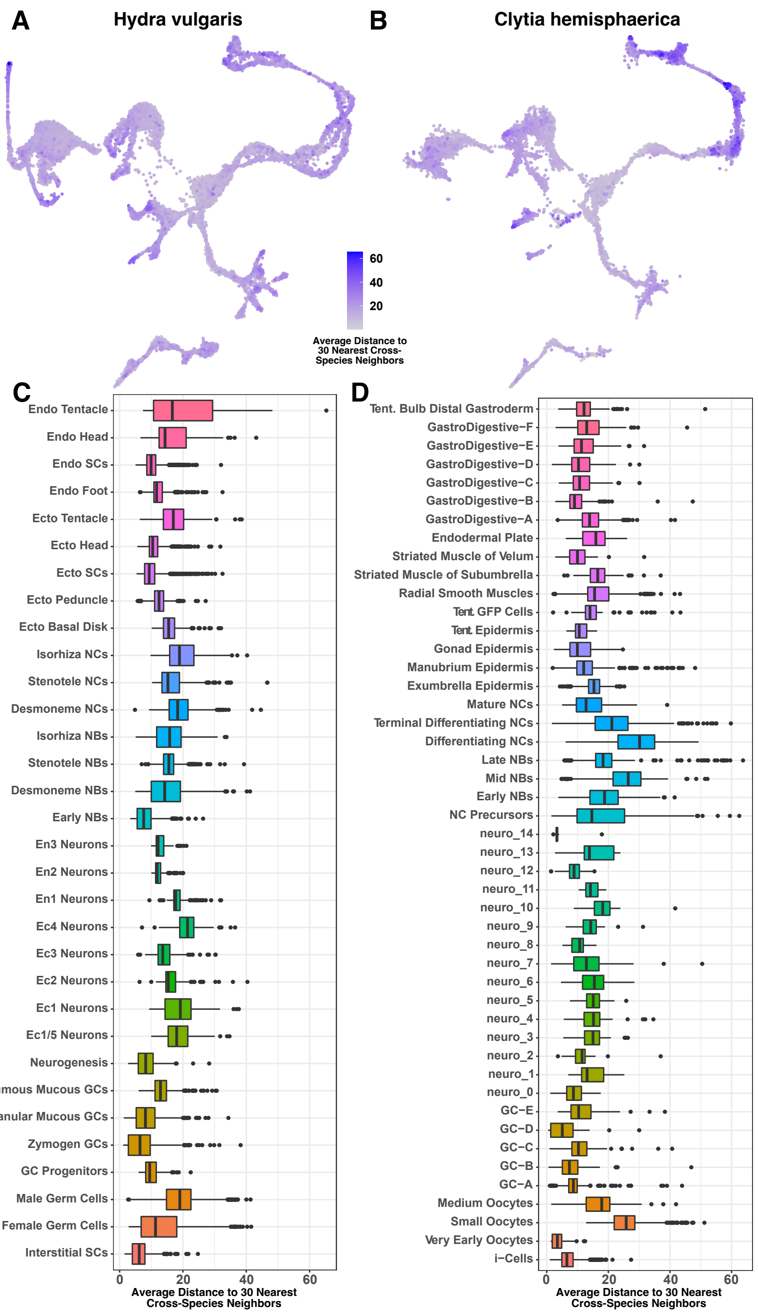


**Fig. S14.** Quantification of alignment distance in the cross-species *Hydra* and *Clytia* single cell atlas. (*A* and *B*) UMAP plots depicting the average distance between (*A*) *Hydra* and (*B*) *Clytia* cells and their 30 nearest cross-species nearest neighbors in aligned principal component space. Cells with lower distance values had transcriptional profiles that were more like cells from the other species. These values were calculated based only on one-to-one orthologs, and thus did not consider transcriptional differences based on genes unique to one of the species. (*C* and *D*)

Box plots showing the distribution of distance scores for (*C*) *Hydra* and (*D*) *Clytia* grouped by cell type.

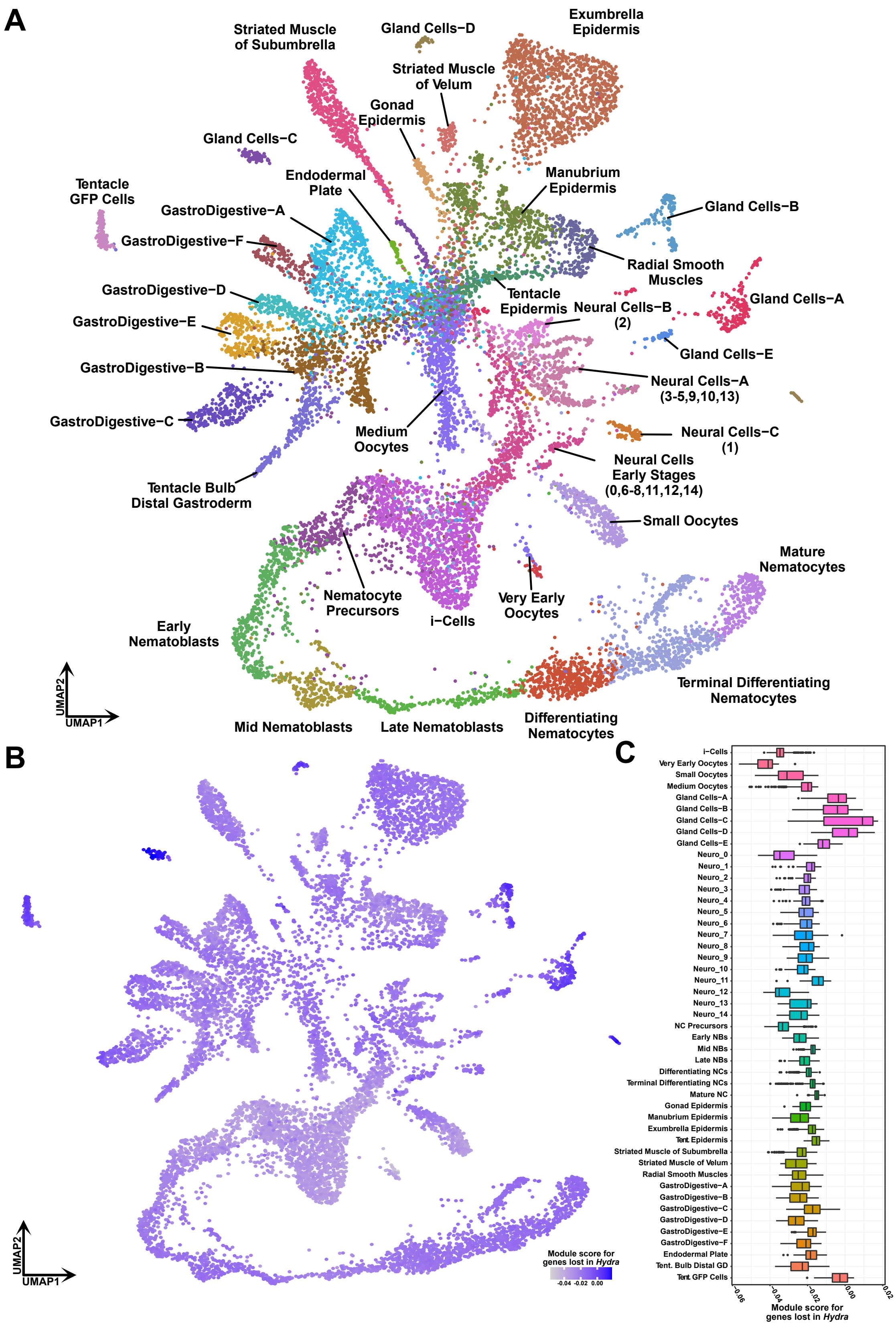
A
Striated Muscle of Subumbrella
Gland Cells−D
Exumbrella Epidermis
Striated Muscle of Velum
Gonad Epidermis
Gland Cells−C
Endodermal Plate
Manubrium Epidermis
Tentacle GFP Cells
GastroDigestive−A
Gland Cells−B
GastroDigestive−F
Radial Smooth Muscles
Tentacle Epidermis
GastroDigestive−D
Neural Cells−B (2)
Gland Cells−A
GastroDigestive−E
Gland Cells−E
GastroDigestive−B
Neural Cells−A (3-5,9,10,13)
GastroDigestive−C
Medium Oocytes
Neural Cells−C (1)
Neural Cells Early Stages (0,6-8,11,12,14)
Tentacle Bulb Distal Gastroderm
Small Oocytes
Mature Nematocytes
Nematocyte Precursors
Very Early Oocytes
i−Cells
Early Nematoblasts
UMAP2
UMAP1
Terminal Differentiating Nematocytes
Mid Nematoblasts
Late Nematoblasts
Differentiating Nematocytes
B
UMAP2
UMAP1
Module score for genes lost in Hydra
-0.04 -0.02 0.00
C
i−Cells
Very Early Oocytes
Small Oocytes
Medium Oocytes
Gland Cells−A
Gland Cells−B
Gland Cells−C
Gland Cells−D
Gland Cells−E
Neuro_0
Neuro_1
Neuro_2
Neuro_3
Neuro_4
Neuro_5
Neuro_6
Neuro_7
Neuro_8
Neuro_9
Neuro_10
Neuro_11
Neuro_12
Neuro_13
Neuro_14
NC Precursors
Early NBs
Mid NBs
Late NBs
Differentiating NCs
Terminal Differentiating NCs
Mature NC
Gonad Epidermis
Manubrium Epidermis
Exumbrella Epidermis
Tent. Epidermis
Striated Muscle of Subumbrella
Striated Muscle of Velum
Radial Smooth Muscles
GastroDigestive−A
GastroDigestive−B
GastroDigestive−C
GastroDigestive−D
GastroDigestive−E
GastroDigestive−F
Endodermal Plate
Tent. Bulb Distal GD
Tent. GFP Cells
-0.06 -0.04 -0.02 0.00 0.02
Module score for genes lost in Hydra

**Fig. S15.** Transcripts expressed in *Clytia* gland and tentacle GFP cells are enriched for genes lost in *Hydra*. (*A*) Original annotated UMAP from the initial *Clytia* atlas publication (Chari et al. 2021). Parenthetical numbers under neuron cluster names refer to neuron subtypes contained within each broad neuron type. Neuron subtype names are based on a neural sub-clustering analysis from the initial atlas publication. Subtypes were assigned to the neuronal cluster that contained the largest portion of cells from a given subtype. (*B-C*) Module scores in the *Clytia* single-cell RNA-seq atlas were calculated based on a weighted average of the expression of all genes lost in *Hydra*. (B) UMAP plot depicting module scores for all single-cell transcriptomes in the *Clytia* atlas. (C) Module scores pooled by cell type.

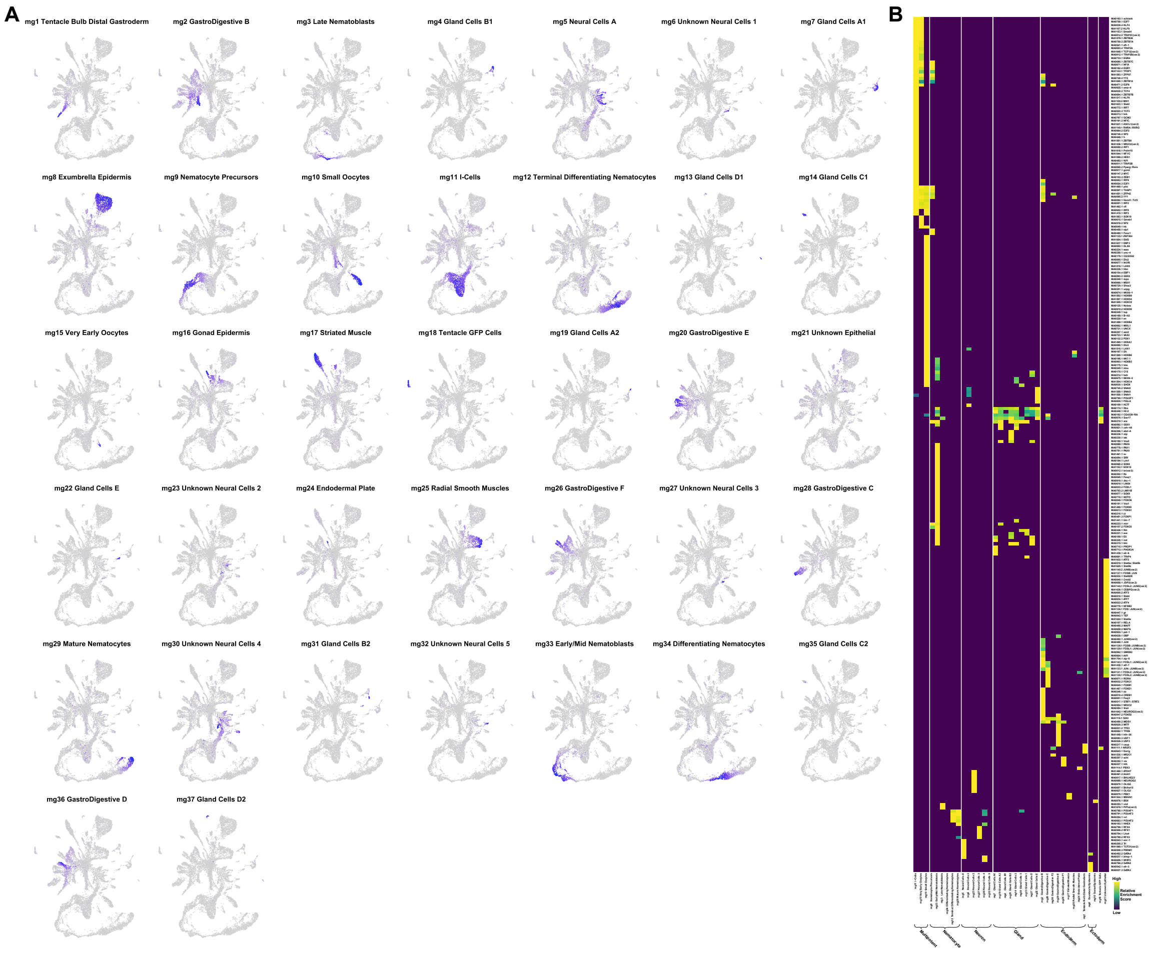


**Fig. S16.** Motif enrichment analysis in the *Clytia* single-cell medusa atlas. (*A*) UMAP plots from the original *Clytia* atlas publication (Chari et al. 2021) colored by non-negative matrix factorization (NMF) metagene expression. NMF identified 37 sets of co-expressed genes in the Clytia atlas, most of which could be readily assigned to previously annotated cell types. (*B*) Heatmap showing enrichment results for promoter proximal (≤ 1 kb from nearest TSS) sequences associated with the 37 metagenes identified by NMF. Sequences were assigned to metagenes based on gene weights generated as part of the standard NMF output. Values are presented only for enrichment results with an E-value < 10 (approximate adjusted p-value of 0.01). Motifs are referred to using both their unique JASPAR ID (formatted as MA####.#) and the abbreviated name of their corresponding TF.

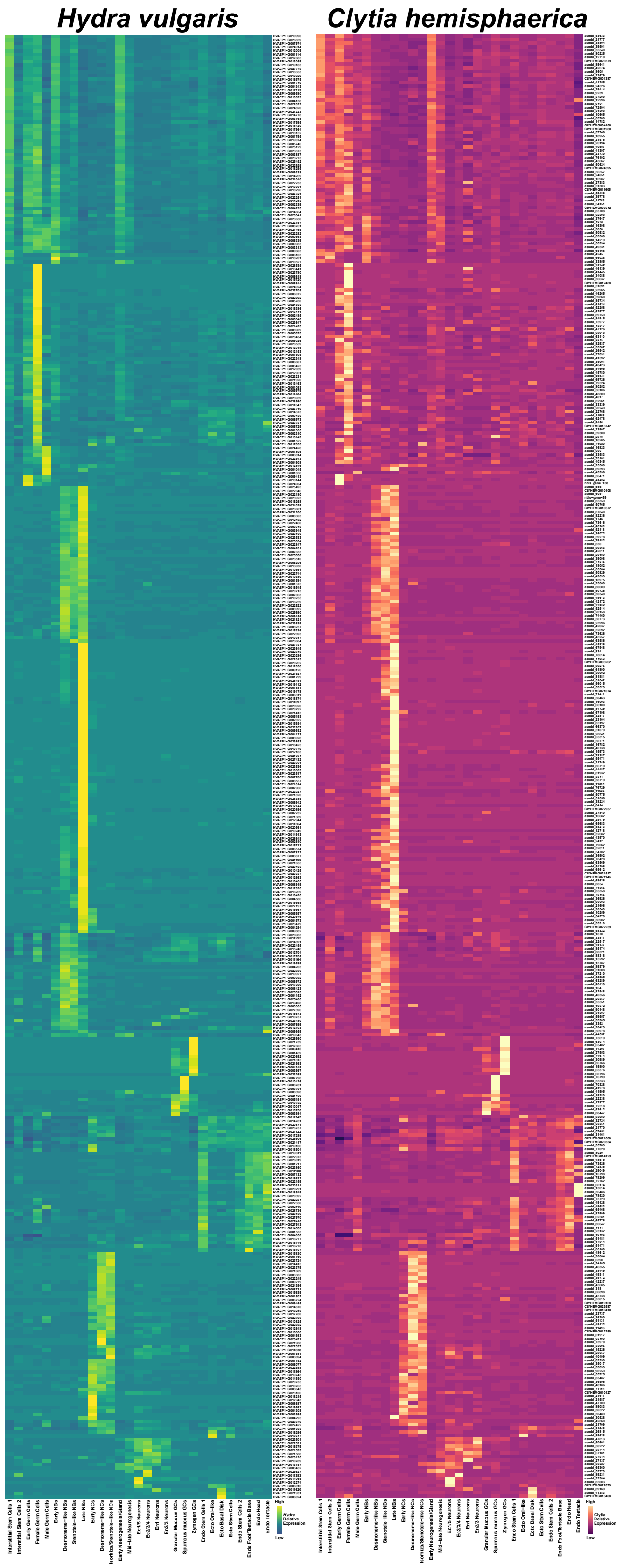

Hydra vulgaris
Clytia hemisphaerica
Interstitial Stem Cells 1
Interstitial Stem Cells 2
Early Germ Cells
Female Germ Cells
Male Germ Cells
Early NBs
Desmoneme−like NBs
Stenotele−like NBs
Late NBs
Early NCs
Desmoneme−like NCs
Isorhiza/Stenotele−like NCs
Early Neurogenesis/Gland
Mid−late Neurogenesis
Ec1/5 Neurons
Ec2/3/4 Neurons
En1 Neurons
En2/3 Neurons
Granular Mucous GCs
Spumous mucous GCs
Zymogen GCs
Endo Stem Cells 1
Ecto Oral−like
Ecto Basal Disk
Ecto Stem Cells
Endo Stem Cells 2
Endo Foot/Tentacle Base
Endo Head
Endo Tentacle
High
Hydra Relative Expression
Low
High
Clytia Relative Expression
Low

**Fig. S17.** Heatmap of orthologous gene pairs with similar cell-type-specific expression in *Hydra* and *Clytia* single-cell atlases. Gene pairs were classified as having similar expression patterns based on correlated expression (correlation score > 0.65) in the aligned cross-species principal component space. The clusters referred to in the heatmap column names refer to a fine-resolution cross-species Louvain clustering analysis presented in Fig S18.

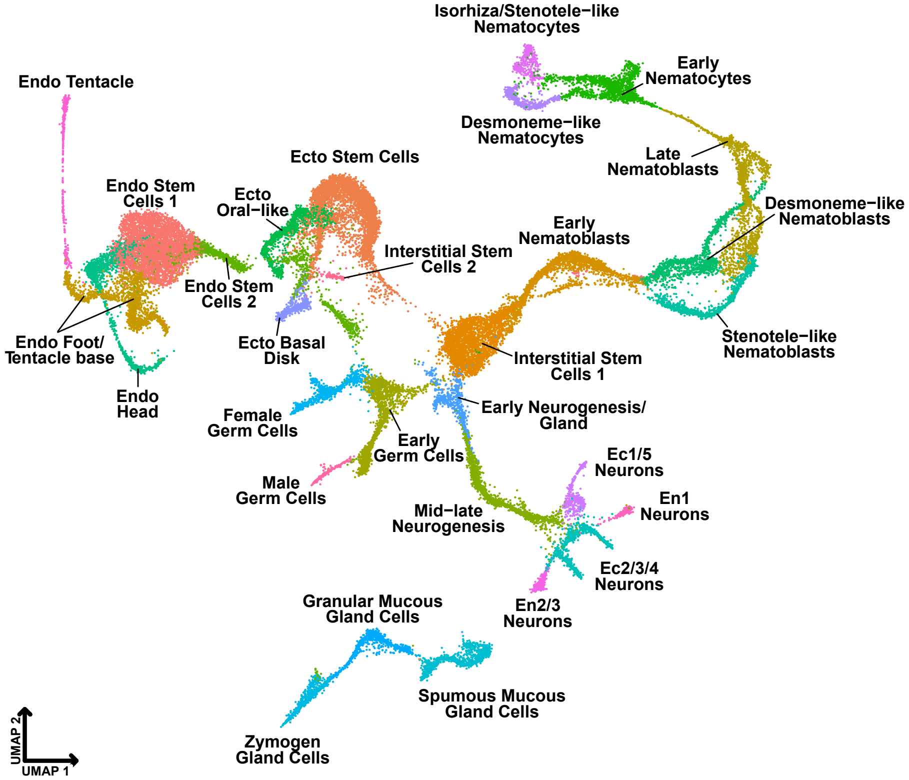


**Fig. S18.** Cross-species aligned *Clytia* and *Hydra* UMAP colored by the Louvain clusters used for the expression correlation heatmaps in Fig 4C and Fig S17.

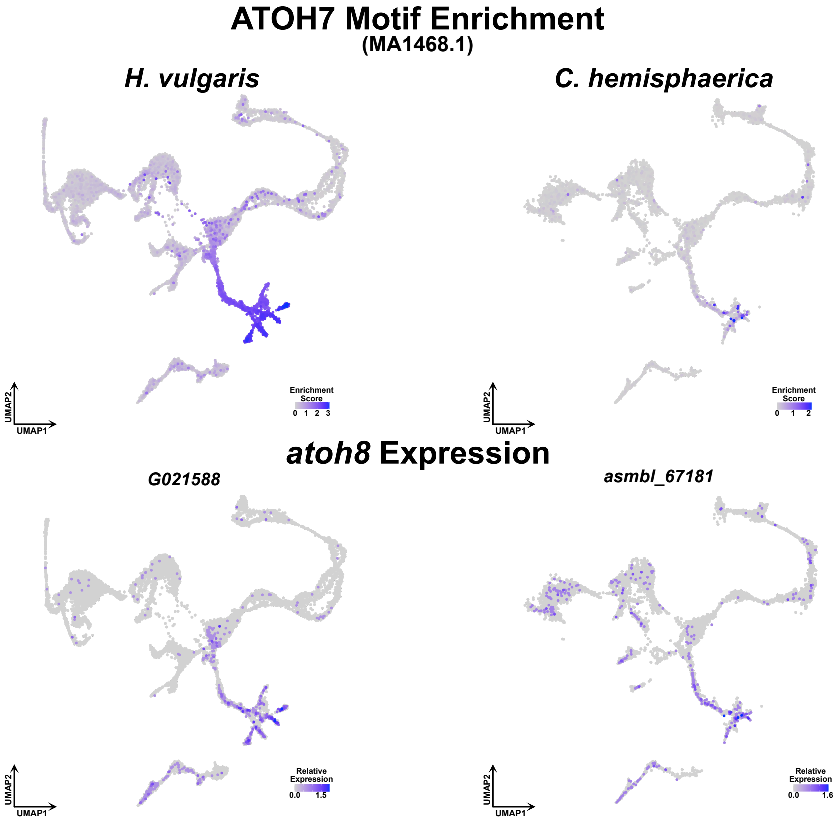


**Fig. S19.** Motif enrichment and gene expression patterns in the *Hydra* and *Clytia* cell atlases suggest *atoh8* is a conserved regulator of hydrozoan neurogenesis.

**Table S1.** Benchmarking Universal Single Copy Orthologs (BUSCOs) statistics for hydrozoan reference sequences.

| | *H. vulgaris*, strain AEP genome (This Study) | *H. vulgaris*, strain 105 genome v3 (Simakov et al., 2022) | *H. vulgaris*, strain 105 genome v2 (Chapman et al., 2010; arusha.nhgri.nih.gov/hydra) | *H. vulgaris*, strain AEP protein models (This Study) | *H. vulgaris*, strain 105 protein models (Chapman et al., 2010; arusha.nhgri.nih.gov/hydra) | *H. vulgaris*, strain AEP LRv2 Transcriptome (Siebert et al., 2019) | *H. oligactis* strain Innsbruck female12 genome (This study) | *H. oligactis*, strain CR genome (Vogg et al., 2019) | *H. oligactis* strain Innsbruck female12 protein models (This Study) | *H. viridissima* genome (Hamada et al., 2020) | *H. viridissima* protein models (Hamada et al., 2020) | *C. hemisphaerica* genome V2 (Leclère et al., 2019; marimba.obs-vlfr.fr/organism/Clytia/hemisphaerica) | *C. hemisphaerica* V2 protein models (Leclère et al., 2019; marimba.obs-vlfr.fr/organism/Clytia/hemisphaerica) | *C. hemisphaerica* protein models (this study) |
|---|---|---|---|---|---|---|---|---|---|---|---|---|---|---|
| **Genes** | NA | NA | NA | 28917 | 33820 | NA | NA | NA | 60590 | NA | 21467 | NA | 19133 | 30058 |
| **Isoforms** | NA | NA | NA | 37784 | 36059 | 38749 | NA | NA | 63777 | NA | 25978 | NA | 28659 | 52872 |
| **Complete BUSCOs** | 866 (90.7%) | 862 (90.3%) | 821 (86.1%) | 903 (94.7%) | 790 (82.8%) | 913 (95.7%) | 841 (88.2%) | 444 (46.5%) | 823 (86.2%) | 839 (88.0%) | 841 (88.2%) | 743 (77.9%) | 774 (81.1%) | 830 (87.0%) |
| **Complete and single-copy BUSCOs** | 859 (90.0%) | 857 (89.8%) | 808 (84.7%) | 890 (93.3%) | 767 (80.4%) | 876 (91.8%) | 718 (75.3%) | 357 (37.4%) | 731 (76.6%) | 828 (86.8%) | 830 (87.0%) | 708 (74.2%) | 688 (72.1%) | 792 (83.0%) |
| **Complete and duplicated BUSCOs** | 7 (0.7%) | 5 (0.5%) | 13 (1.4%) | 13 (1.4%) | 23 (2.4%) | 37 (3.9%) | 123 (12.9%) | 87 (9.1%) | 92 (9.6%) | 11 (1.2%) | 11 (1.2%) | 35 (3.7%) | 86 (9.0%) | 38 (4.0%) |
| **Fragmented BUSCOs** | 39 (4.1%) | 41 ( 4.3%) | 61 (6.4%) | 16 (1.7%) | 86 (9.0%) | 9 (0.9%) | 63 (6.6%) | 290 (30.4%) | 74 (7.8%) | 59 (6.2%) | 41 (4.3%) | 92 (9.6%) | 28 (2.9%) | 30 (3.1%) |
| **Missing BUSCOs** | 49 (5.2%) | 51 (5.4%) | 72 (7.5%) | 35 (3.6%) | 78 (8.2%) | 32 (3.4%) | 50 (5.2%) | 220 (23.1%) | 57 (6.0%) | 56 (5.8%) | 72 (7.5%) | 119 (12.5%) | 152 (16.0%) | 94 (9.9%) |

**Table S2.** List of sources for proteomes used in the OrthoFinder analysis. * indicates transcriptomes that were translated into protein sequence before being used in the analysis.

| Species | Citation | Download Source |
|---|---|---|
| *Acropora millepora* | Ying et al., 2019 | ncbi.nlm.nih.gov/genome/2652 |
| *Adineta vaga* | Flot et al., 2013 | metazoa.ensembl.org/Adineta_vaga/Info/Index |
| *Amphimedon queenslandica* | Srivastava et al., 2010; Fernandez-Valverde et al., 2015 | metazoa.ensembl.org/Amphimedon_queenslandica/Info/Index |
| *Aurelia aurita* | Gold et al., 2019 | davidadlergold.faculty.ucdavis.edu/jellyfish/ |
| *Bombyx mori* | Xia et al., 2008; Duan et al., 2009 | metazoa.ensembl.org/Bombyx_mori/Info/Index |
| *Branchiostoma lanceolatum* | Marlétaz et al., 2018 | metazoa.ensembl.org/Branchiostoma_lanceolatum/Info/Index |
| *Caenorhabditis elegans* | Harris et. al., 2019; C. elegans Sequencing Consortium, 1998 | uswest.ensembl.org/Caenorhabditis_elegans/Info/Index |
| *Callorhinchus milii* | Venkatesh et al., 2014 | uswest.ensembl.org/Callorhinchus_milii/Info/Index |
| *Calvadosia cruxmelitensis* | Odhera et al., 2019 | github.com/josephryan/Ohdera_et_al_2018/tree/master/AA_Files |
| *Cassiopea xamachana* | Odhera et al., 2019 | mycocosm.jgi.doe.gov/Casxa1/Casxa1.home.html |
| *Ciona intestinalis* | Satou et al., 2008 | uswest.ensembl.org/Ciona_intestinalis/Info/Index |
| *Clytia hemisphaerica* | Leclère et al., 2019; marimba.obs-vlfr.fr/organism/Clytia/hemisphaerica | Final protein models were generated by this study |
| *Daphnia pulex* | Colbourne et al., 2011 and Colbourne et al., 2005 | metazoa.ensembl.org/Daphnia_pulex/Info/Annotation/ |
| *Drosophila melanogaster* | Adams et al., 2000 and Clark et al., 2007 | uswest.ensembl.org/Drosophila_melanogaster/Info/Index |
| *Ephydatia muelleri* | Kenny et al., 2020 | spaces.facsci.ualberta.ca/ephybase/ |
| *Exaiptasia diaphana* | Baumgarten et al., 2015 | www.ncbi.nlm.nih.gov/genome/92231 |
| *Gallus gallus* | Hillier et al., 2004 | uswest.ensembl.org/Gallus_gallus/Info/Index |
| *Hofstenia miamia* | Gehrke et al., 2019 | metazoa.ensembl.org/Hofstenia_miamia/Info/Index |
| *Homo sapiens* | International Human Genome Sequencing Consortium, 2001; Venter et al., 2001; Genome Reference Consortium | uswest.ensembl.org/Homo_sapiens/Info/Index |
| *Hydra circumcincta** | Wong et al., 2019 | Obtained by contacting study authors |
| *Hydra oligactis** | Wong et al., 2019 | Obtained by contacting study authors |
| *Hydra viridissima* | Hamada et al., 2020 | marinegenomics.oist.jp/hydra_viridissima_a99/viewer/download?project_id=82 |
| *Hydra vulgaris, strain 105* | Chapman et al., 2010 | arusha.nhgri.nih.gov/hydra/download/genemodels_proteins/hydra2.0_genemodels.aa.gz |
| *Hydractinia echinata** | NA | research.nhgri.nih.gov/hydractinia/ |
| *Latimeria chalumnae* | NA | uswest.ensembl.org/Latimeria_chalumnae/Info/Index |
| *Lepisosteus oculatus* | Braasch et al., 2016 | uswest.ensembl.org/Lepisosteus_oculatus/Info/Index |
| *Mnemiopsis leidyi* | Ryan et al., 2013 and Moreland et al., 2020 | research.nhgri.nih.gov/mnemiopsis/download/download.cgi?dl=proteome |
| *Monosiga brevicollis* | King et al., 2008 | protists.ensembl.org/Monosiga_brevicollis_mx1_gca_000002865/Info/Index |
| *Morbakka virulenta* | Khalturin et al., 2019 | marinegenomics.oist.jp/morbakka_virulenta/viewer/download?project_id=70 |
| *Mus Musculus* | Mouse Genome Sequencing Consortium, 2002; Genome Reference Consortium | uswest.ensembl.org/Mus_musculus/Info/Index |
| *Nematostella vectensis* | Zimmermann et al., 2020 | simrbase.stowers.org/starletseaanemone |
| *Octopus bimaculoides* | Albertin et al., 2015 | metazoa.ensembl.org/Octopus_bimaculoides/Info/Index |
| *Patiria miniata* | Kudtarkar and Cameron, 2017 | legacy.echinobase.org/Echinobase/PmDownload |
| *Petromyzon marinus* | Smith et al., 2013 | uswest.ensembl.org/Petromyzon_marinus/Info/Index |
| *Pleurobrachia bachei* | Moroz et al., 2014 | neurobase.rc.ufl.edu/Pleurobrachia/download |
| *Porites lutea* | Robbins et al., 2019 | plut.reefgenomics.org/download/ |
| *Praesagittifera naikaiensis* | Arimoto et al., 2019 | gigadb.org/dataset/100564 |
| *Rhopilema esculentum* | Li et al., 2020 | gigadb.org/dataset/view/id/100720/File_page |
| *Schmidtea mediterranea* | Robb et al., 2015 | planosphere.stowers.org/smedgd |
| *Scolanthus callimorphus* | Zimmermann et al., 2020 | simrbase.stowers.org/wormanemone |
| *Strongylocentrotus purpuratus* | Sea Urchin Genome Sequencing Consortium 2006 and Cameron et al., 2009 | metazoa.ensembl.org/Strongylocentrotus_purpuratus/Info/Index |
| *Trichoplax adhaerens* | Srivastava et al., 2008 | metazoa.ensembl.org/Trichoplax_adhaerens/Info/Index |
| *Xenopus tropicalis* | Hellsten et al., 2010 | uswest.ensembl.org/Xenopus_tropicalis/Info/Index |

**Supplemental Data S1.** Excel worksheet containing functional annotation data for the *H. vulgaris* strain AEP gene models. The first tab contains the combined results from our OrthoFinder, InterProScan, and BLAST annotation approaches. For these annotations, the longest isoform was used as the representative sequence for each gene. For the OrthoFinder results, the assigned orthogroup is included along with predicted orthologs from a select set of well-annotated bilaterian species. Specifically, the bilaterian orthologs were drawn from the *Homo sapiens*, *Mus musculus*, *Xenopus tropicalis*, *Drosophila melanogaster*, and *Caenorhabditis elegans* proteomes. For each strain AEP gene model, we only included orthologs from one of the five species, prioritizing the species based using the following hierarchy: 1) *H. sapiens*, 2) *M. musculus*, 3) *X. tropicalis*, 4) *D. melanogaster*, and 5) *C. elegans* (e.g., if the AEP gene had no orthologs in *H. sapiens*, we included the orthologs from *M. musculus*. If there were no orthologs in either *H. sapiens* or *M. musculus* we included the orthologs from *X. tropicalis*, etc.). The table also contains the Pfam and PANTHER predictions from the InterProScan analysis. The "*Hydra* GenBank BLAST Hit" column contains the best BLAST hits, when available, in a custom database of manually deposited *Hydra* transcript sequences from GenBank. Finally, the table includes the best BLAST hit, when available, for each gene model in the UniProtKB protein database. The second tab in the worksheet contains the InterProScan predictions for all putative transcription factors in the strain AEP gene models.

**Supplemental Data S2.** Excel workbook containing consensus peak coordinates for ATAC-seq and CUT&Tag datasets. The workbook includes peak sets for all biologically reproducible peaks (irreproducible discovery rate $\leq 0.1$) as well as peak sets that are only those reproducible peaks that were also conserved in at least two other *Hydra* genomes. Conservation status was determined by using k-means clustering to partition peaks into two populations (conserved and non-conserved) based on the percent of conserved bases for each non-AEP *Hydra* genome. The first six columns in the peak tables are identical to BED-formatted genome-coordinate files. H3K4me1, H3K4me3, and ATAC-seq peak lists also include three additional columns, generated via the UROPA annotation pipeline (Kondili et al. 2017), that provide information on the nearest gene to each peak.

**Supplemental Data S3.** Table containing motif conservation analysis results. The ‘Conserved Motif Hits’ column represents the number of predicted instances of each motif in the AEP genome that were conserved in at least two other non-AEP genomes. The ‘Total Motif Hits’ column represents the total number of predicted instances for each motif in the AEP genome. ‘Motif Conservation Rate’ represents the ratio between ‘Conserved Motif Hits’ and ‘Total Motif Hits’. The ‘Conserved Shuffled Motif Hits’, ‘Total Shuffled Motif Hits’, and ‘Shuffled Motif Conservation Rate’ columns are the same metrics as described above applied to a shuffled version of the same motif. Shuffled motifs are non-functional sequences with identical lengths and sequence biases as their non-shuffled equivalents. The ‘Log-Odds Ratio’ column represents the log-transformed ratio of the odds that an instance of the non-shuffled motif will be conserved compared to the same odds for the shuffled control motif. Positive values indicate higher rates of conservation in the non-shuffled motif. The ‘Rank’ column represents how strongly a motif was enriched relative to all other motifs in the analysis, with the smallest values corresponding to the highest log-odds ratio and the highest values corresponding to the lowest log-odds ratio. The ‘P-value’ and ‘FDR’ columns contain the results of a chi-square test comparing the conservation rates of shuffled and non-shuffled versions of the motif. The ‘Enrichment Result’ summarizes the classification for each motif based on the results of the chi-square test. We classified a motif as ‘enriched’ if the chi-square FDR was ≤ 0.01 and the log-odds ratio was > 0 and ‘depleted’ if the chi-square FDR was ≤ 0.01 and the log-odds ratio was < 0. A motif was classified as ‘neutral’ if the chi-square FDR was > 0.01.

**Supplemental Data S4.** Genome coordinates for all predicted motif instances in the AEP genome that were conserved in at least two other non-AEP *Hydra* genomes. The format is identical to the coordinate files in Supplemental Data S2.

**Supplemental Data S5.** BED-formatted genome coordinate file for chromatin contact domains generated by the hicFindTADs function from the HiCExplorer pipeline (Ramírez et al. 2018). The fifth column contains the insulation score at the start of each domain. The insulation score measures chromatin interaction levels, with lower values indicating a localized decrease in contact frequency.

**Supplemental Data S6.** List of cell-type-specific markers for the strain AEP *H. vulgaris* single-cell RNA-seq atlas. Markers were found by comparing single cell transcriptomes of a given cell-type to all other cells in the atlas using a Wilcoxon Rank Sum test as implemented in Seurat. Markers were excluded if the estimated $\log_2$ fold-change was less than 1. Cluster names in the 'Target Cluster' column use the following abbreviations: Ec, ectoderm; En, endoderm; I, interstitial; SC, stem cell; BodyCol, body column; NB, nematoblast; NC, nematocyte; Nem, nematogenesis; N, neuron; Neuro, neurogenesis; Gl, gland cell; GC, germ cell; Progen, progenitor; Zymo, zymogen; SpumMuc, spumous mucous; Fem, female; ISC, interstitial stem cell; Desmo, desmoneme; Steno, stenotele; Iso, isorhiza.

**Supplemental Data S7.** Metagene by gene matrix generated by NMF analysis of the *Hydra* cell atlas. Values in the matrix are Z-scores that measure how enriched a gene was in each metagene. Positive values represent stronger associations between a gene and a metagene.

**Supplemental Data S8.** Cell by metagene matrix generated by NMF analysis of the *Hydra* cell atlas. Values in the matrix reflect the extent to which each metagene contributes to the overall transcriptomic profile of each single-cell transcriptome, with higher values reflecting a stronger relative contribution. Values in this matrix are unitless.

**Supplemental Data S9.** Excel workbook containing the phylostratigraphically estimated age for all *Hydra* genes that were assigned to an orthogroup in our OrthoFinder analysis. The table includes the gene model ID ("*Hydra* Gene ID"), the most ancient clade that contained all predicted orthologs of that gene ("Clade of Origin"), and the orthofinder-assigned node ID for the clade of origin ("Orthofinder Node ID").

**Supplemental Data S10.** Motif by cell-type matrices containing the enrichment results used to generate the heatmaps in Fig. 3J and Fig. S12. Values in the matrix are derived from normalized enrichment scores (NES) calculated using a gene set enrichment analysis (GSEA) (Subramanian et al. 2005; Korotkevich et al. 2021), with higher values indicating stronger enrichment for that motif in the specified cell type. Cell types with non-significant enrichment results (adjusted P-value > 0.01) were set to zero to reduce noise.

**Supplemental Data S11.** Table of candidate regulators of gene co-expression in *Hydra*. Each row of the table represents a different candidate regulator. Genes were designated as candidate regulators of a motif if its expression correlated with the motif's enrichment pattern in the single cell atlas and the gene possessed a DNA binding domain that could bind the motif. Along with the regulator gene ID, each row includes all motifs with enrichment patterns that were correlated with the gene's expression pattern (correlation score > 0.5) listed in decreasing order, functional annotations based on bilaterian orthologs identified by Orthofinder, all Pfam protein domains predicted by InterProScan, and GenBank accession numbers for the gene based on best BLAST hits against a curated list of *Hydra* GenBank entries.

**Supplemental Data S12.** Metagene by gene matrix generated by NMF analysis of the *Clytia* cell atlas. Values in the matrix are Z-scores that measure how enriched a gene was in each metagene. Positive values represent stronger associations between a gene and a metagene.

**Supplemental Data S13.** Cell by metagene matrix generated by NMF analysis of the *Clytia* cell atlas. Values in the matrix reflect the extent to which each metagene contributes to the overall transcriptomic profile of each single-cell transcriptome, with higher values reflecting a stronger relative contribution. Values in this matrix are unitless.

**Supplemental Data S14.** Excel worksheet containing results from the cross-species transcriptional regulation analysis. The 'Clytia Motif Enrichment' tab contains the output from the *Clytia* metagene Analysis of Motif Enrichment (AME). The 'Mg Promoters w/ Motif' column represents the number of genes that belong to a given metagene that had a predicted instance of the target motif. The '% MG Promoters w/ Motif' represents this number as a percentage of the total number of genes belonging to a given metagene. The 'Non-mg Promoters w/ Motif' and '% Non-mg Promoters w/ Motif' are equivalent metrics that instead refer to the genes that were not part of the target metagene. The 'Fold-Enrichment' was calculated by dividing the '% Mg Promoters w/ Motif' by the '% Non-mg Promoters w/ Motif'. All motifs with an E-value > 10 were excluded from AME results tables. The 'Cross-Species Motif Cor' tab contains enrichment pattern correlation scores for all motifs that were enriched in both the *Hydra* and *Clytia* atlases. High correlation scores indicate a motif was enriched in similar cell types in the two species. The 'Cross-Species Shuf Motif Cor' tab contains enrichment pattern correlation scores for shuffled, non-functional motif sequences.

**Supplemental Data S15.** Expression correlation scores for orthologous gene pairs in the *Hydra* and *Clytia* cell atlases. Ortholog pairs were deemed to have similar expression in *Clytia* and *Hydra* if their pseudo-cell correlation score was ≥ 0.65. In addition to the *Hydra* and *Clytia* gene IDs and the pseudo-cell correlation score, each row also contains functional annotations based on bilaterian orthologs identified by Orthofinder (both the abbreviated name and the Ensembl ID) and GenBank accession numbers for the gene based on best BLAST hits against a curated list of *Hydra* GenBank entries.

**References**

Armstrong J, Hickey G, Diekhans M, Fiddes IT, Novak AM, Deran A, Fang Q, Xie D, Feng S, Stiller J, et al. 2020. Progressive Cactus is a multiple-genome aligner for the thousandgenome era. *Nature* **587**: 246–251. http://dx.doi.org/10.1038/s41586-020-2871-y.

Bateman A, Martin MJ, Orchard S, Magrane M, Agivetova R, Ahmad S, Alpi E, Bowler-Barnett EH, Britto R, Bursteinas B, et al. 2021. UniProt: The universal protein knowledgebase in 2021. *Nucleic Acids Res* **49**: D480–D489.

Blum M, Chang H-Y, Chuguransky S, Grego T, Kandasaamy S, Mitchell A, Nuka G, Paysan-Lafosse T, Qureshi M, Raj S, et al. 2021. The InterPro protein families and domains database: 20 years on. *Nucleic Acids Res* **49**: D344–D354.

Bode H, Lengfeld T, Hobmayer B, Holstein TW. 2009. Detection of Expression Patterns in Hydra Pattern Formation. In *Wnt Signaling* (ed. E. Vincan), pp. 69–84, Humana Press, Totowa, NJ https://doi.org/10.1007/978-1-60327-469-2_7.

Bode HR, Flick KM. 1976. Distribution and dynamics of nematocyte populations in Hydra attenuata. *J Cell Sci* **21**: 15–34.

Bolger AM, Lohse M, Usadel B. 2014. Trimmomatic: a flexible trimmer for Illumina sequence data. *Bioinformatics* **30**: 2114–2120. https://www.ncbi.nlm.nih.gov/pubmed/24695404.

Brůna T, Hoff KJ, Lomsadze A, Stanke M, Borodovsky M. 2021. BRAKER2: Automatic eukaryotic genome annotation with GeneMark-EP+ and AUGUSTUS supported by a protein database. *NAR Genomics Bioinforma* **3**: 1–11.

Buenrostro JD, Giresi PG, Zaba LC, Chang HY, Greenleaf WJ. 2013. Transposition of native chromatin for fast and sensitive epigenomic profiling of open chromatin, DNA-binding proteins and nucleosome position. *Nat Methods* **10**: 1213–1218. https://www.ncbi.nlm.nih.gov/pubmed/24097267.

Buenrostro JD, Wu B, Chang HY, Greenleaf WJ. 2015. ATAC-seq: A Method for Assaying Chromatin Accessibility Genome-Wide. *Curr Protoc Mol Biol* **109**: 21 29 1–9. https://www.ncbi.nlm.nih.gov/pubmed/25559105.

Cazet JF, Cho A, Juliano C. 2021. Generic injuries are sufficient to induce ectopic Wnt organizers in Hydra. *Elife* **10**. https://elifesciences.org/articles/60562.

Chapman JA, Kirkness EF, Simakov O, Hampson SE, Mitros T, Weinmaier T, Rattei T, Balasubramanian PG, Borman J, Busam D, et al. 2010. The dynamic genome of Hydra.

*Nature* **464**: 592–596. https://www.ncbi.nlm.nih.gov/pubmed/20228792.

Chari T, Weissbourd B, Gehring J, Ferraioli A, Leclère L, Herl M, Gao F, Chevalier S, Copley RR, Houliston E, et al. 2021. Whole-animal multiplexed single-cell RNA-seq reveals transcriptional shifts across Clytia medusa cell types. *Sci Adv* **7**: 2021.01.22.427844. https://www.biorxiv.org/content/10.1101/2021.01.22.427844v1%0Ahttps://www.biorxiv.org/content/10.1101/2021.01.22.427844v1.abstract.

Corces MR, Trevino AE, Hamilton EG, Greenside PG, Sinnott-Armstrong NA, Vesuna S, Satpathy AT, Rubin AJ, Montine KS, Wu B, et al. 2017. An improved ATAC-seq protocol reduces background and enables interrogation of frozen tissues. *Nat Methods* **14**: 959–962. https://www.ncbi.nlm.nih.gov/pubmed/28846090.

Dana CE, Glauber KM, Chan TA, Bridge DM, Steele RE. 2012. Incorporation of a Horizontally Transferred Gene into an Operon during Cnidarian Evolution. *PLoS One* **7**.

Dobin A, Davis CA, Schlesinger F, Drenkow J, Zaleski C, Jha S, Batut P, Chaisson M, Gingeras TR. 2013. STAR: Ultrafast universal RNA-seq aligner. *Bioinformatics* **29**: 15–21.

Domazet-Lošo T, Brajković J, Tautz D. 2007. A phylostratigraphy approach to uncover the genomic history of major adaptations in metazoan lineages. *Trends Genet* **23**: 533–539. https://linkinghub.elsevier.com/retrieve/pii/S0168952507002995.

Domazet-Lošo T, Tautz D. 2010. A phylogenetically based transcriptome age index mirrors ontogenetic divergence patterns. *Nature* **468**: 815–819.

Dudchenko O, Batra SS, Omer AD, Nyquist SK, Hoeger M, Durand NC, Shamim MS, Machol I, Lander ES, Aiden AP, et al. 2017. De novo assembly of the Aedes aegypti genome using Hi-C yields chromosome-length scaffolds. *Science (80- )* **356**: 92–95.

Durand NC, Shamim MS, Machol I, Rao SSP, Huntley MH, Lander ES, Aiden EL. 2016. Juicer Provides a One-Click System for Analyzing Loop-Resolution Hi-C Experiments. *Cell Syst* **3**: 95–98. http://dx.doi.org/10.1016/j.cels.2016.07.002.

Emms DM, Kelly S. 2019. OrthoFinder: phylogenetic orthology inference for comparative genomics. *Genome Biol* **20**: 238. https://genomebiology.biomedcentral.com/articles/10.1186/s13059-015-0721-2.

English AC, Richards S, Han Y, Wang M, Vee V, Qu J, Qin X, Muzny DM, Reid JG, Worley KC, et al. 2012. Mind the Gap: Upgrading Genomes with Pacific Biosciences RS Long-Read Sequencing Technology. *PLoS One* **7**: 1–12.

Flynn JM, Hubley R, Goubert C, Rosen J, Clark AG, Feschotte C, Smit AF. 2020. RepeatModeler2 for automated genomic discovery of transposable element families. *Proc Natl Acad Sci U S A* **117**: 9451–9457.

Fornes O, Castro-Mondragon JA, Khan A, van der Lee R, Zhang X, Richmond PA, Modi BP, Correard S, Gheorghe M, Baranašić D, et al. 2020. JASPAR 2020: update of the open-access database of transcription factor binding profiles. *Nucleic Acids Res* **48**: D87–D92. https://doi.org/10.1093/nar/gkz1001.

Gierer A, Berking S, Bode H, David CN, Flick K, Hansmann G, Schaller H, Trenkner E. 1972. Regeneration of Hydra from Reaggregated Cells. *Nat New Biol* **239**: 98–101. http://www.nature.com/articles/newbio239098a0.

Glauber KM, Dana CE, Park SS, Colby DA, Noro Y, Fujisawa T, Chamberlin AR, Steele RE, Glauber KM, Dana CE, et al. 2015. A small molecule screen identifies a novel compound that induces a homeotic transformation in Hydra, (Development (Cambridge), (2015) 142, 4788-4796). *Dev* **142**: 2081.

Grabherr MG, Haas BJ, Yassour M, Levin JZ, Thompson DA, Amit I, Adiconis X, Fan L, Raychowdhury R, Zeng Q, et al. 2011. Full-length transcriptome assembly from RNA-Seq data without a reference genome. *Nat Biotechnol* **29**: 644–652.

Grant CE, Bailey TL, Noble WS. 2011. FIMO: Scanning for occurrences of a given motif. *Bioinformatics* **27**: 1017–1018.

Haas BJ, Delcher AL, Mount SM, Wortman JR, Smith RK, Hannick LI, Maiti R, Ronning CM, Rusch DB, Town CD, et al. 2003. Improving the Arabidopsis genome annotation using maximal transcript alignment assemblies. *Nucleic Acids Res* **31**: 5654–5666.

Hafemeister C, Satija R. 2019. Normalization and variance stabilization of single-cell RNA-seq data using regularized negative binomial regression. *Genome Biol* **20**: 296. http://www.ncbi.nlm.nih.gov/pubmed/31870423.

Hahne F, Ivanek R. 2016. Visualizing Genomic Data Using Gviz and Bioconductor. In *Statistical Genomics: Methods and Protocols* (eds. E. Mathé and S. Davis), pp. 335–351, Springer New York, New York, NY https://doi.org/10.1007/978-1-4939-3578-9_16.

Hamada M, Satoh N, Khalturin K. 2020. A Reference Genome from the Symbiotic Hydrozoan, Hydra viridissima. *G3 (Bethesda)* **10**: 3883–3895.

Hao Y, Hao S, Andersen-Nissen E, Mauck WM, Zheng S, Butler A, Lee MJ, Wilk AJ, Darby C, Zager M, et al. 2021. Integrated analysis of multimodal single-cell data. *Cell* **184**: 3573-3587.e29. https://doi.org/10.1016/j.cell.2021.04.048.

Hickey G, Paten B, Earl D, Zerbino D, Haussler D. 2013. HAL: A hierarchical format for storing and analyzing multiple genome alignments. *Bioinformatics* **29**: 1341–1342.

Hufnagel LA, Kass-Simon G, Lyon MK. 1985. Functional organization of battery cell complexes in tentacles of Hydra attenuata. *J Morphol* **184**: 323–341.

Jackman SD, Coombe L, Chu J, Warren RL, Vandervalk BP, Yeo S, Xue Z, Mohamadi H, Bohlmann J, Jones SJM, et al. 2018. Tigmint: correcting assembly errors using linked reads from large molecules. *BMC Bioinformatics* **19**: 393. http://www.ncbi.nlm.nih.gov/pubmed/30367597.

Jones P, Binns D, Chang HY, Fraser M, Li W, McAnulla C, McWilliam H, Maslen J, Mitchell A, Nuka G, et al. 2014. InterProScan 5: Genome-scale protein function classification. *Bioinformatics* **30**: 1236–1240.

Kaya-Okur HS, Wu SJ, Codomo CA, Pledger ES, Bryson TD, Henikoff JG, Ahmad K, Henikoff S. 2019. CUT&Tag for efficient epigenomic profiling of small samples and single cells. *Nat Commun* **10**: 1–10.

Kolmogorov M, Yuan J, Lin Y, Pevzner PA. 2019. Assembly of long, error-prone reads using repeat graphs. *Nat Biotechnol* **37**: 540–546. http://dx.doi.org/10.1038/s41587-019-0072-8.

Kondili M, Fust A, Preussner J, Kuenne C, Braun T, Looso M. 2017. UROPA: a tool for Universal RObust Peak Annotation. *Sci Rep* **7**: 2593. https://www.ncbi.nlm.nih.gov/pubmed/28572580.

Koren S, Walenz BP, Berlin K, Miller JR, Bergman NH, Phillippy AM. 2017. Canu: Scalable and accurate long-read assembly via adaptive κ-mer weighting and repeat separation. *Genome Res* **27**: 722–736.

Korotkevich G, Sukhov V, Budin N, Shpak B, Artyomov MN, Sergushichev A. 2021. Fast gene set enrichment analysis. *bioRxiv*. https://www.biorxiv.org/content/early/2021/02/01/060012.

Kotliar D, Veres A, Nagy MA, Tabrizi S, Hodis E, Melton DA, Sabeti PC. 2019. Identifying gene expression programs of cell-type identity and cellular activity with single-cell RNA-Seq. *Elife* **8**: 1–26.

Langmead B, Salzberg SL. 2012. Fast gapped-read alignment with Bowtie 2. *Nat Methods* **9**: 357–359. https://www.ncbi.nlm.nih.gov/pubmed/22388286.

Leclère L, Horin C, Chevalier S, Lapébie P, Dru P, Peron S, Jager M, Condamine T, Pottin K, Romano S, et al. 2019. The genome of the jellyfish Clytia hemisphaerica and the evolution of the cnidarian life-cycle. *Nat Ecol Evol* **3**: 801–810. http://dx.doi.org/10.1038/s41559-019-0833-2.

Lenhoff HM, Brown RD. 1970. Mass culture of hydra: an improved method and its application to other aquatic invertebrates. *Lab Anim* **4**: 139–154.

Li B, Dewey CN. 2011. RSEM: accurate transcript quantification from RNA-Seq data with or without a reference genome. *BMC Bioinformatics* **12**: 323. https://www.ncbi.nlm.nih.gov/pubmed/21816040.

Li H. 2018. Minimap2: Pairwise alignment for nucleotide sequences. *Bioinformatics* **34**: 3094–3100.

Li H, Handsaker B, Wysoker A, Fennell T, Ruan J, Homer N, Marth G, Abecasis G, Durbin R. 2009. The Sequence Alignment/Map format and SAMtools. *Bioinformatics* **25**: 2078–2079.

Li QH, Brown JB, Huang HY, Bickel PJ. 2011. Measuring Reproducibility of High-Throughput Experiments. *Ann Appl Stat* **5**: 1752–1779.

Lopez-Delisle L, Rabbani L, Wolff J, Bhardwaj V, Backofen R, Grüning B, Ramírez F, Manke T. 2021. pyGenomeTracks: reproducible plots for multivariate genomic datasets. *Bioinformatics* **37**: 422–423.

Macosko EZ, Basu A, Satija R, Nemesh J, Shekhar K, Goldman M, Tirosh I, Bialas AR, Kamitaki

N, Martersteck EM, et al. 2015. Highly Parallel Genome-wide Expression Profiling of Individual Cells Using Nanoliter Droplets. *Cell* **161**: 1202–1214. http://ac.els-cdn.com/S0092867415005498/1-s2.0-S0092867415005498-main.pdf?_tid=0e4f0f52-99a6-11e6-98a6-00000aab0f27&acdnat=1477285096_f5a41a99ac302d13e90633d9f464cdf4.

Martin VJ, Littlefield CL, Archer WE, Bode HR. 1997. Embryogenesis in Hydra. *Biol Bull* **192**: 345–363. https://www.journals.uchicago.edu/doi/10.2307/1542745.

McInnes L, Healy J, Saul N, Großberger L. 2018. UMAP: Uniform Manifold Approximation and Projection. *J Open Source Softw* **3**: 861.

McLeay RC, Bailey TL. 2010. Motif Enrichment Analysis: A unified framework and an evaluation on ChIP data. *BMC Bioinformatics* **11**.

Meers MP, Tenenbaum D, Henikoff S. 2019. Peak calling by Sparse Enrichment Analysis for CUT&RUN chromatin profiling. *Epigenetics and Chromatin* **12**: 1–11. https://doi.org/10.1186/s13072-019-0287-4.

Ouyang JF, Kamaraj US, Cao EY, Rackham OJL. 2021. ShinyCell: simple and sharable visualization of single-cell gene expression data. *Bioinformatics* **37**: 3374–3376.

Rahat A, Rahat M, Searle JB. 1985. A simple method for the preparation of hydra chromosome spreads: introducing chromosome counts into hydra taxonomy. *Experientia* **41**: 282–283. http://link.springer.com/10.1007/BF02002638.

Ramírez F, Bhardwaj V, Arrigoni L, Lam KC, Grüning BA, Villaveces J, Habermann B, Akhtar A, Manke T. 2018. High-resolution TADs reveal DNA sequences underlying genome organization in flies. *Nat Commun* **9**. http://dx.doi.org/10.1038/s41467-017-02525-w.

Ramírez F, Ryan DP, Grüning B, Bhardwaj V, Kilpert F, Richter AS, Heyne S, Dündar F, Manke T. 2016. deepTools2: a next generation web server for deep-sequencing data analysis. *Nucleic Acids Res* **44**: W160–W165. https://academic.oup.com/nar/article-lookup/doi/10.1093/nar/gkw257.

Rathje K, Mortzfeld B, Hoeppner MP, Taubenheim J, Bosch TCG, Klimovich A. 2020. *Dynamic interactions within the host-associated microbiota cause tumor formation in the basal metazoan Hydra*. http://dx.doi.org/10.1371/journal.ppat.1008375.

Reddy PC, Gungi A, Ubhe S, Galande S. 2020. Epigenomic landscape of enhancer elements during Hydra head organizer formation. *Epigenetics and Chromatin* **13**: 1–16. https://doi.org/10.1186/s13072-020-00364-6.

Roach MJ, Schmidt SA, Borneman AR. 2018. Purge Haplotigs: allelic contig reassignment for third-gen diploid genome assemblies. *BMC Bioinformatics* **19**: 460. http://www.ncbi.nlm.nih.gov/pubmed/30497373.

Siebert S, Farrell JA, Cazet JF, Abeykoon Y, Primack AS, Schnitzler CE, Juliano CE. 2019. Stem cell differentiation trajectories in Hydra resolved at single-cell resolution. *Science (80- )* **365**: eaav9314. http://www.sciencemag.org/lookup/doi/10.1126/science.aav9314.

Simakov O, Bredeson J, Berkoff K, Marletaz F, Mitros T, Schultz DT, O'Connell BL, Dear P, Martinez DE, Steele RE, et al. 2022. Deeply conserved synteny and the evolution of metazoan chromosomes. *Sci Adv* **8**. https://www.science.org/doi/10.1126/sciadv.abi5884.

Slater GSC, Birney E. 2005. Automated generation of heuristics for biological sequence comparison. *BMC Bioinformatics* **6**: 1–11.

Subramanian A, Tamayo P, Mootha VK, Mukherjee S, Ebert BL, Gillette MA, Paulovich A, Pomeroy SL, Golub TR, Lander ES, et al. 2005. Gene set enrichment analysis: A knowledge-based approach for interpreting genome-wide expression profiles. *Proc Natl Acad Sci U S A* **102**: 15545–15550.

Sun S, White RR, Fischer KE, Zhang Z, Austad SN, Vijg J. 2020. Inducible aging in Hydra oligactis implicates sexual reproduction, loss of stem cells, and genome maintenance as major pathways. *GeroScience* **42**: 1119–1132.

Tarashansky AJ, Musser JM, Khariton M, Li P, Arendt D, Quake SR, Wang B. 2021. Mapping single-cell atlases throughout metazoa unravels cell type evolution. *Elife* **10**: 1–24.

Vogg MC, Beccari L, Iglesias Ollé L, Rampon C, Vriz S, Perruchoud C, Wenger Y, Galliot B. 2019. An evolutionarily-conserved Wnt3/β-catenin/Sp5 feedback loop restricts head organizer activity in Hydra. *Nat Commun* **10**: 312. http://dx.doi.org/10.1038/s41467-018-08242-2.

Voigt O, Erpenbeck D, Worheide G. 2008. A fragmented metazoan organellar genome: the two mitochondrial chromosomes of Hydra magnipapillata. *BMC Genomics* **9**: 350. http://bmcgenomics.biomedcentral.com/articles/10.1186/1471-2164-9-350.

Walker BJ, Abeel T, Shea T, Priest M, Abouelliel A, Sakthikumar S, Cuomo CA, Zeng Q, Wortman J, Young SK, et al. 2014. Pilon: an integrated tool for comprehensive microbial variant detection and genome assembly improvement. *PLoS One* **9**: e112963. https://bmcbioinformatics.biomedcentral.com/articles/10.1186/s12859-018-2425-6.

Waltman L, Van Eck NJ. 2013. A smart local moving algorithm for large-scale modularity-based community detection. *Eur Phys J B* **86**.

Wong WY, Simakov O, Bridge DM, Cartwright P, Bellantuono AJ, Kuhn A, Holstein TW, David CN, Steele RE, Martínez DE. 2019. Expansion of a single transposable element family is associated with genome-size increase and radiation in the genus Hydra. *Proc Natl Acad Sci U S A* **116**: 22915–22917.

Yeo S, Coombe L, Warren RL, Chu J, Birol I. 2018. ARCS: Scaffolding genome drafts with linked reads. *Bioinformatics* **34**: 725–731.

Yu S, Westfall JA, Dunne JF. 1985. Light and electron microscopic localization of a monoclonal antibody in neurons in situ in the head region of Hydra. *J Morphol* **184**: 183–193.

Zacharias H, Anokhin B, Khalturin K, Bosch TCG. 2004. Genome sizes and chromosomes in the basal metazoan Hydra. *Zoology* **107**: 219–227.

Zhang Y, Liu T, Meyer CA, Eeckhoute J, Johnson DS, Bernstein BE, Nussbaum C, Myers RM, Brown M, Li W, et al. 2008. Model-based Analysis of ChIP-Seq (MACS). *Genome Biol* **9**: R137. http://genomebiology.biomedcentral.com/articles/10.1186/gb-2008-9-9-r137.
